# Exploring the extracellular regulation of the tumor angiogenic interaction network using a systems biology model

**DOI:** 10.1101/581884

**Authors:** Ding Li, Stacey D. Finley

**Affiliations:** Department of Biomedical Engineering, University of Southern California, Los Angeles, CA, USA; Department of Biomedical Engineering; Mork Family Department of Chemical Engineering and Materials Science; Department of Biological Sciences, University of Southern California, Los Angeles, CA, USA

## Abstract

Tumor angiogenesis is regulated by pro- and anti-angiogenic factors. Anti-angiogenic agents target the interconnected network of angiogenic factors to inhibit neovascularization, which subsequently impedes tumor growth. Due to the complexity of this network, optimizing anti-angiogenic cancer treatments requires detailed knowledge at a systems level. In this study, we constructed a tumor tissue-based model to better understand how the angiogenic network is regulated by opposing mediators at the extracellular level. We consider the network comprised of two pro-angiogenic factors: vascular endothelial growth factor (VEGF) and basic fibroblast growth factor (FGF2), and two anti-angiogenic factors: thrombospondin-1 (TSP1) and platelet factor 4 (PF4). The model’s prediction of angiogenic factors’ distribution in tumor tissue reveals the localization of different factors and indicates the angiogenic state of the tumor. We explored how the distributions are affected by the secretion of the pro- and anti-angiogenic factors, illustrating how the angiogenic network is regulated in the extracellular space. Interestingly, we identified a counterintuitive result that the secretion of the anti-angiogenic factor PF4 can enhance pro-angiogenic signaling by elevating the levels of the interstitial and sur-face-level pro-angiogenic species. This counterintuitive situation is pertinent to the clinical setting, such as the release of anti-angiogenic factors in platelet activation or the administration of exogenous PF4 for anti-angiogenic therapy. Our study provides mechanistic insights into this counterintuitive result and highlights the role of heparan sulfate proteoglycans in regulating the interactions between angiogenic factors. This work complements previous studies aimed at understanding formation of angiogenic complexes in tumor tissue and helps in the development of anti-cancer strategies targeting angiogenesis.

## 1 INTRODUCTION

Angiogenesis, the growth of new blood microvessels from pre-existing microvasculature, plays a crucial role in tumor development^1^. Tumor growth relies on angiogenesis to enable waste exchange and provide oxygen and nutrients from the surrounding environment. Several angiogenic factors that affect the extent of tumor vascularization have been identified and are commonly categorized as pro- and anti-angiogenic factors. Pro-angiogenic factors, including vascular endothelial growth factor-A (VEGF) and fibroblast growth factor 2 (FGF2), bind to their respective receptors to induce pro-angiogenic signaling promoting cell proliferation, cell migration and blood vessel formation^2,3^. On the other side, anti-angio-genic factors, like thrombospodin-1 (TSP1) and platelet factor 4 (PF4), inhibit pro-angiogenic signaling and induce anti-angiogenic signaling to oppose angiogenesis^4,5^. Considering the importance of angiogenesis in tumor development, anti-angiogenic therapies are designed to target the signaling of angiogenic factors to inhibit neovascularization and tumor growth^6^. Single-agent anti-angiogenic therapies that target a particular angiogenic factor in the network were the first angi-ogenesis-inhibiting therapies studied. These include antibodies or small molecules targeting pro-angiogenic factors^7^ and peptide mimetics of anti-angiogenic factors^8^. However, these single-agent anti-angiogenic therapies showed limited success in the clinic due to toxicity, low efficacy, or the development of resistance^6,9^. These drawbacks have promoted efforts to develop combination therapies administering multiple anti-angiogenic agents that simultaneously target various angiogenic species in the network^10–13^.

Due to the intrinsic complexity of the network regulating tumor angiogenesis, optimizing anti-angiogenic cancer treatment, specifically combination anti-angiogenic therapy, requires detailed knowledge and a holistic view at a systems level. Computational systems biology models offer powerful tools to systematically study tumor angiogenesis and optimize anti-angiogenic tumor therapy. Various types of systems biology models have been constructed to investigate new anti-angiogenic therapies^14^. Models of intracellular signaling of angiogenic factors characterize the biochemical events inside the cell initiated by ligand binding to signaling receptors on the cell surface. These models help in the identification of new intracellular drug targets. At the extracellular level, models of the extracellular species’ reaction network are used to understand the distribution of angiogenic factors in tumor tissue^15^ and in the whole body^16^. By linking to the kinetics of anti-angiogenic drugs, models that capture extracellular interactions can be used to study therapeutics that modulate the distribution of angiogenic factors, which directly affects angiogenic signaling^17,18^. To better understand the effects of targeting angiogenic factors in the tumor, we built a new tissue-based systems biology model characterizing the extracellular network that involves four main angiogenic factors regulating tumor angiogenesis, including VEGF, FGF2, TSP1 and PF4.

Our modeling work expanded previous models by incorporating angiogenic factors that were previously omitted from the models, as well as other significant mediators. Thus, our model enables a systematic study of the extracellular regulation of multiple angiogenic factors. The extracellular distribution of VEGF alone was firstly investigated in a computational setting with a tissue-based model^15^. Then this physiologically relevant and molecularly detailed model was extended to include TSP1, a potent endogenous anti-angiogenic factor, to explore the balance of pro- and anti-angiogenic factor in tumor tissue^19^. In the present work, we further expand the model to include the pro-angiogenic factor, FGF2, and an additional anti-angiogenic factor, PF4. These species are reported to interact with VEGF and TSP1 and significantly impact on tumor angiogenesis. FGF2 is reported to synergistically enhance the pro-angiogenic signal with VEGF^20,21^. On the other hand, upregulation of the FGF2 pathway can result in resistance to anti-VEGF therapy^10,13^. PF4, like the other anti-angiogenic factor TSP1, binds to VEGF and FGF2 to reduce pro-angiogenic signaling^22,23^. Therefore, incorporating FGF2 and PF4 provides a more complete view of the angiogenic interaction network and a more comprehensive understanding of tumor angiogenic state, as compared to previous models. In addition, PF4, TSP1, VEGF and FGF2 each bind to heparin, competing for the heparan sulfate (HS) binding sites in heparan sulfate proteoglycans (HSPG) on the cell surface and in the extracellular matrix and basement membrane^24^. The secretion of PF4 and TSP1 leading to displacement of VEGF and FGF2 from HS binding sites is an important mechanism of tumor angiogenesis regulation. Specifically, PF4 is known to interrupt the HSPG-mediated formation of pro-angiogenic complexes to inhibit VEGF and FGF2 signaling^25,26^. To account for the regulation of HSPG, our model includes two distinct species with HS binding sites, one of which, the surface-level HSPG, is not explicitly accounted for in previous tumor tissue-based models^15,19^.

With the newly constructed model, we firstly profiled the distribution of these four angiogenic factors in tumor tissue and systematically investigated how the secretion of different angiogenic factors affects the balance of pro- and anti-angiogenic signaling. Furthermore, we generate insights explaining two specific counterintuitive phenomena: (1) the secretion of PF4 increases the levels of free VEGF and FGF2 in tumor tissue and (2) the secretion of PF4 promotes the formation of VEGF signaling complexes. We found HSPG’s level directly affects these counterintuitive results in different ways, emphasizing the important role of HSPGs in the regulation of angiogenic factor signaling. Lastly, we apply the model to simulate a controlled release of PF4 in tumor tissue, and our results indicate that the HSPG level in the tumor microenvironment might affect the response to platelet activation and recombinant PF4 anti-angiogenic therapy. Overall, we establish a new computational framework to understand the extracellular distribution of angiogenic factors in tumor tissue and generate new insights into the regulation of the angiogenic factors’ interaction network, which are difficult to examine through experimental study alone.

## 2 METHODS

### The tumor tissue model of angiogenic factors

We constructed a molecularly detailed model that describes the extracellular network of four main angiogenic factors in tumor tissue (Fig.1). The modeling approach is consistent with previous works^15,19^. The system is represented by a set of coupled nonlinear ordinary differential equations (ODEs) to characterize a well-mixed tumor tissue. For the model structure, the extracellular spaces in tumor tissue are divided into three regions: the surface of endothelial cells, the surface of tumor cells and interstitial space. The interstitial space between tumor and endothelial cells is comprised of extracellular matrix (ECM) and the basement membranes surrounding the tumor cells (TBM) and the endothelial cells (EBM). The soluble species are secreted by both cell types and can be removed from the system through degradation in the interstitial space or internalization with receptors at the cell surface.

**Figure 1.**
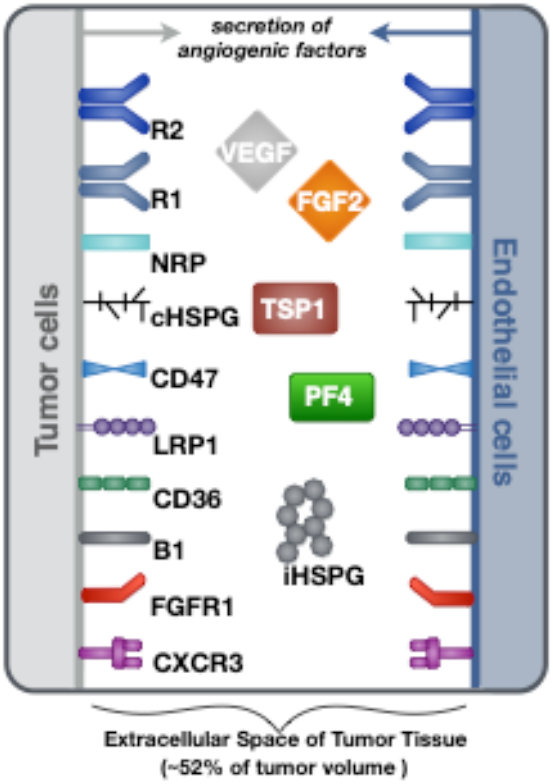
Schematic of the tumor tissue based model. The compartmental model describes the reaction network of four major angiogenic factors (VEGF, FGF2, TSP1 and PF4) in the extracellular space of tumor tissue. Angiogenic receptors are expressed on the cell surfaces. Soluble angiogenic factors exist in the interstitial space, and they bind to cell surface receptors to form signaling complexes. Two different heparan sulfate proteoglycans are included in the model, including cell surface HSPG (cHSPG) and interstitial HSPG (iHSPG).

Ten soluble species are present in the model (Fig. 2, Legend I). Physiologically, in tumor tissue, VEGF_121_, VEGF_165_, FGF2, TSP1, MMP3 and proMMP9 are mainly produced through the secretion from tumor cells and endothelial cells, while PF4 is stored in the α-granules of the platelets and is released through platelet activation. Thus, in the model, the source of PF4 in tumor tissue is represented by a generic production rate. In addition, VEGF_114_, inactive TSP1 and active MMP9 are formed through cleavage. Nine relevant receptors are present on the cell surface (Fig. 2, Legend II), including VEGF receptors (VEGFR1, VEGFR2, Neuropilin-1), TSP1 receptors (CD47, CD36, LRP1, α_x_β_1_ integrins), FGF2 receptor (FGFR1) and PF4 receptor (CXCR3). Receptors are assumed to be uniformly distributed on the cell surface and are recycled back to surface to maintain a constant total number for each type of receptor. In addition, we include the heparan sulfate proteoglycans (HSPGs) in the model, which are important modulators of angiogenic signaling. HSPGs are glycoproteins, which have a protein core and one or more covalently attached heparan sulfate (HS) chains. Two types of HSPGs are present in the model (Fig. 2, Legend III). One is the interstitial heparan sulfate proteoglycans (iHSPG) that are present in the ECM, EBM and TBM. The other one is the cell-surface heparan sulfate proteoglycans (cHSPG). The iHSPG serves as a reservoir for angiogenic factors, while the cHSPG mainly functions as a co-receptor participating in the formation of complexes to modulate angiogenic signaling.

**Figure 2.**
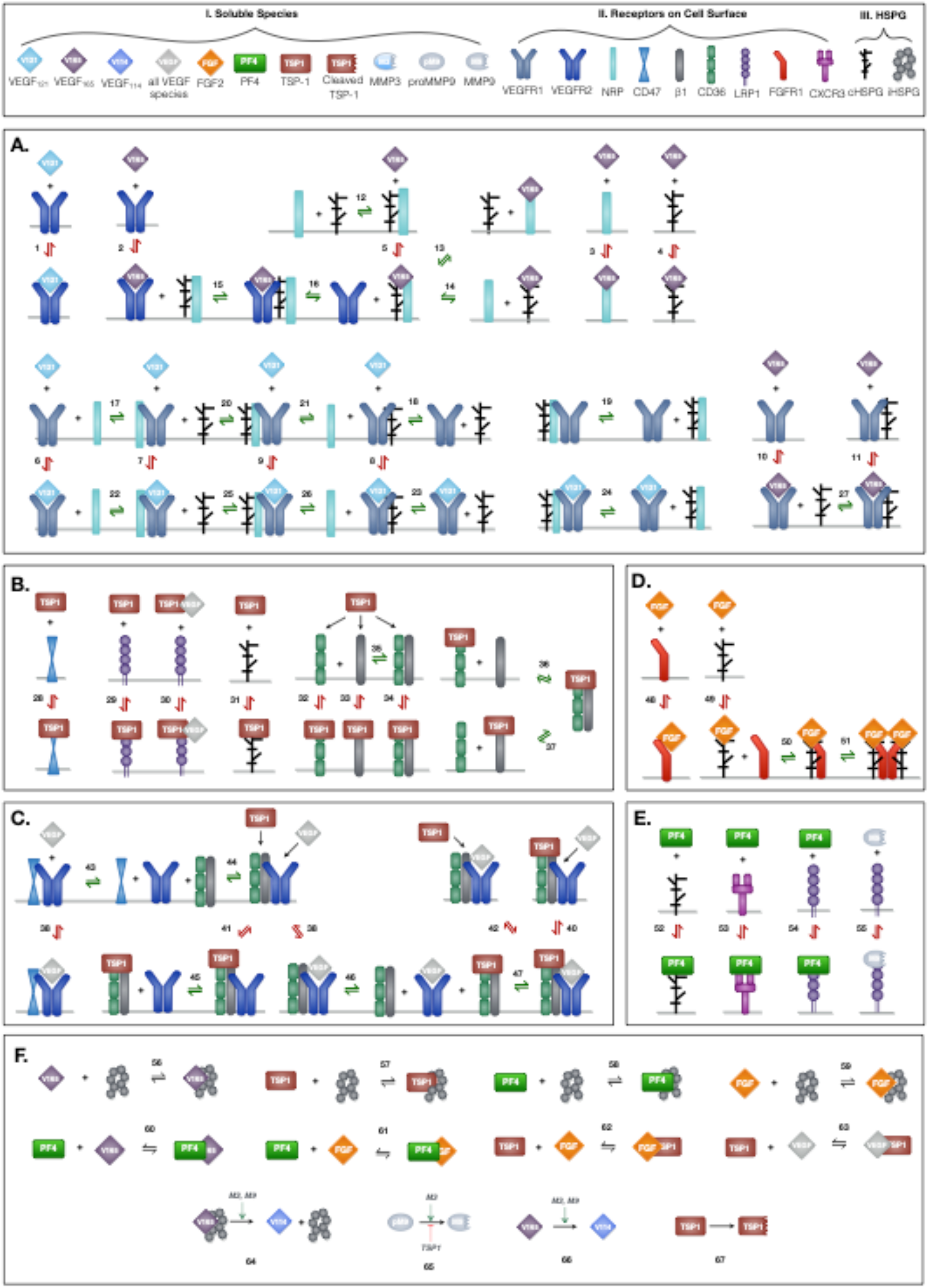
Schematic of the extracellular network of VEGF, FGF2, TSP1 and PF4. (A) Molecular interactions of two active VEGF isoforms (VEGF_165_ and VEGF_121_), receptors (VEGFR1, VEGFR2 and NRP1) and heparan sulfate proteoglycans on the cell surface (cHSPG). (B) Molecular interactions of TSP1 binding to its receptors (CD36, CD47, LRP1 and α_x_β_1_ integrins) and cHSPG. (C) Molecular interactions of the coupling between VEGFR2 and TSP1 receptors. (D) Molecular interactions of FGF2 binding to FGFR1 and cHSPG and the formation of the full signaling complex through dimerization. (E) Molecular interactions of PF4 binding to receptors (CXCR3 and LRP1) and cHSPG, as well as MMP9 binding to LRP1. (F) The molecular interactions of angiogenic factors binding to one another and heparan sulfate proteoglycans in the interstitial space (iHSPG), as well as the proteolysis and degradation of soluble species. Numbers for each reaction correspond to the list of reactions in Supplementary File S1.

### Network of reactions

The principles of mass action kinetics are used to characterize the species’ dynamics. The defined rules that govern the molecular interactions are showed in Fig. 2, and the detailed reactions are given in Supplementary File S1.

#### *VEGF-Receptor axis* (Fig. 2A)

Previous work modeling VEGF ligand-receptor interactions did not explicitly include the surface-level HSPGs (cHSPG)^27^, assuming the presence of abundant HSPGs on the cell surface. To investigate the impact of HSPG on VEGF signaling, we extended previous VEGF-VEGFR modeling to incorporate the cHSPG-facilitated VEGF binding reactions. Previous works have detailed documentation of estimating the kinetic for two VEGF isoforms (VEGF_165_ and VEGF_121_) binding to VEGF receptors^15,27^, and we use those parameter values in our model. Below, we present how we have adapted previous works to include cHSPG regulation.

VEGFR2 and co-receptors (first two rows of Fig. 2A): According its structure, VEGF_165_ binds to VEGFR2 via the exon 4 encoded domain and to NRP1 and HSPG via the exon 7 encoded domain to form a ternary complex^27,28^. It is commonly assumed that VEGFR2 does not directly interact with NRP1, but is bridged by the VEGF_165_^29^. For the HSPG, in a recent study, VEGFR2 was shown not to interact with heparin directly, and that VEGF_165_ also mediates the interactions between VEGR2 and heparin^30^. Therefore, in our model, we assume HSPG does not directly interact with VEGFR2, and the impact of HSPG on VEGFR2 signaling is mediated through supporting the VEGF_165_-mediated bridging of VEGFR2 with NRP1. For the interactions between HSPG and NRP1, it is reported that heparin could bind to the b1b2 domain of NRP1 directly, greatly enhancing the binding of VEGF_165_ to NRP1^31^. This suggests that VEGF_165_ binds to NRP1 in an HSPG-dependent way. To include this knowledge, we allow HSPG to pre-couple with NRP1 before interacting with VEGFR2. The other isoform of VEGF, VEGF_121_, lacks the exon 7 coded region; thus, it does not bind to NRP1 or HSPG^28,30^.

VEGFR1 and co-receptors (second two row of Fig. 2A): Following our previous modeling^15,27^, VEGFR1 can couple with NRP1, while the binding with VEGF_121_ is not affected by the coupling. Since VEGR1 was shown to bind to heparin directly and VEGFR1 does not show a heparin-aided VEGF binding as VEGR2 does^30^, we assume HSPG can couple with VEGR1 and does not affect its binding to VEGF. In addition, we assume VEGFR1 can couple with the NRP1 pre-coupled with HSPG to form a ternary complex and that then binds with VEGF. Following our previous model, VEGF_165_ does not bind to VEGFR1 pre-coupled with NRP1^15,27^. In addition, since it is reported that the presence of heparin does not significantly change the binding of VEGF_165_ to VEGFR1^30^, we assume the pre-coupling of VEGFR1 with HSPG does not affect VEGFR1’s binding with VEGF_165_.

#### *TSP1-Receptor axis* (Fig. 2B-C)

The reaction involving interactions between TSP1 and its receptors are taken from previous works^19^, in which TSP1 regulates angiogenic signaling in different ways. TSP1 binds to its own receptors to induce anti-angiogenic signaling (Fig. 2B). Ligated TSP1 receptors can also couple with VEGFR2 to inhibit the signaling of VEGF (Fig. 2C). These interactions are included in the model.

#### *FGF2-Receptor axis* (Fig. 2D)

The reactions for the FGF2-receptor axis are from the extracellular part of an *in vitro* whole cell FGF2 signaling model^32^, which defines the formation of FGF2 signaling trimeric complexes that then dimerize. FGF2 binds to HSPG to form a complex, which binds to the FGFR1 monomer to form a trimeric complex. Then, dimerization of the trimeric complex leads to the formation of the full FGF2 signaling complex. The choice of this ordering is based on several observations from experimental studies^32,33^: FGF2 shows a lower affinity to FGFR1 than to heparin; the interaction of FGFR1 and heparin has a very weak affinity; and FGF2 dramatically increases the association of FGFR1 with heparin. Alternative orders of the binding reactions are possible; however, they are reported to not conform well with the experimental data^33^.

#### *PF4-Receptor axis* (Fig. 2E)

PF4 regulates angiogenesis through various mechanisms. On the cell surface, PF4 binds to cell surface receptors (CXCR3 and LRP1) to induce anti-angiogenic signaling^34,35^ and binds to cHSPG to control pro-angiogenic signaling^22,26^. We include these interactions in the model.

#### *Interactions between angiogenic factors* (Fig. 2F)

The angiogenic factors also interact either by direct binding or through HSPGs in the interstitial space(iHSPG). TSP1 associates with VEGF and FGF2 to reduce pro-angiogenic signaling^19,36^, and it mediates VEGF cleavage through MMP activity^19^. In addition, TSP1 can compete for the HS binding sites on cHSPG and iHSPG to release HS-bound angiogenic factors. Similarly, FGF2 can be trapped by TSP1, PF4 and iHSPG. Additionally, PF4 directly binds to VEGF_165_ and FGF2, reducing the available pro-angiogenic factors^22,23^. Lastly, PF4 competes for the HS binding sites on the iHSPG.

### Parameterization

The model parameter values are reported in Supplementary File S2 with literature references. Here, we describe the derivation of inherited values and the rationales for the parameterization of newly introduced values.

#### Geometric parameters

The tumor tissue is parameterized as a 33 cm^3^ breast tumor, which is modeled as a spatially averaged compartment in the model (Fig.1). The geometric parameters define the volume of the compartment, the interstitial space volume fraction, and the tissue surface areas of endothelial cells and tumor cells. These geometric parameters enable the conversion of concentration from moles per cm^3^ tissue to standard units (pmol/l), where the derivations are thoroughly documented in previous works^15,16^.

#### Production and degradation of soluble species

The production and degradation rates of VEGF, TSP1, MMP3 and proMMP9 are estimated in our previous work^19^. The baseline production rates of PF4 and FGF2 are set to match an intermediate level within the range of experimental measurements (Table 1). The degradation rates of PF4 and FGF2 are set according to their half-life (*t*_1/2_): the rate of degradation is *In*(2)/*t*_l/2_. Since a wide range of reported values for the FGF2 half-life is found in literature^37,38^, we assume it has a half-life of 60 minutes, similar to VEGF, which is within the reported range. PF4 is reported to be rapidly cleared in human body, where the half-life is assumed to be 5 minutes^39^.

**Table 1.** Comparison of the baseline predictions and the experimental measurements of VEGF, FGF2, TSP1, PF4 and MMPs.

| Species | Range of Experimental Measurements <sup>†</sup> | Predicted Concentration | Source and References |
| --- | --- | --- | --- |
| VEGF unbound | 8.0 - 389 pM | 180.2 pM | Multiple<sup>63</sup> |
| TSP1 total <sup>‡</sup> | 1.0 - 6.2 nM (2.0) | 3.0 nM | Breast cancer patient<sup>64</sup> |
| PF4 unbound | 1.0 - 11.3 nM | 4.7 nM | Multiple<sup>52-55</sup> |
| FGF2 total | 0.2 - 11.1 nM | 3.9 nM | Prostate cancer patient<sup>65</sup> |
| MMP3 total | 1.8 - 65.1 nM (5.1) | 5.0 nM | Oral squamous cell carcinoma patient<sup>66</sup> |
| MMP9 total | 1.0 - 287.8 nM (9.0) | 9.2 nM | Oral squamous cell carcinoma patient<sup>66</sup> |
| MMP9 active | 0 - 22.4 nM (0.8) | 0.2 nM | Oral squamous cell carcinoma patient<sup>66</sup> |

<sup>†</sup> Median value is shown in parentheses, if provided in literature. If the experimental data reflects the total concentration in tissue, we assume 50% of the total protein amount is in the extracellular space.
<sup>‡</sup> TSP1 concentration includes both active TSP1 and cleaved TSP1.

#### Receptor Numbers

The receptor densities for VEGF receptors, TSP1 receptors, FGFR1 and HSPGs are taken from previous modeling works^19,32^. There is a paucity of measurements for the PF4 receptor, CXCR3. Thus, we referred to the qualitative measurements in Human Protein Atlas^40^, assuming “low”, “medium”, and “high” expression levels correspond to 2500, 5000, and 10,000 receptors per cell. CXCR3 has a low expression, which is set to be to be 2,500 receptors per cell accordingly.

#### Kinetic Parameters

For the VEGF axis, the kinetic parameters have been estimated in previous work, based on experimental measurements^27^ and assuming an abundant level of HSPGs. We adopted these values in our current model by incorporating several experimentally observed synergistic interactions in the presence of heparin. Since the previous model is calibrated in condition with abundant HSPGs, we assume the NRP1 in the previous model is already coupled with HSPGs. Therefore, the parameters of VEGF_165_ binding to NRP1 in the previous model is used for VEGF_165_ binding to the NRP1:HSPG complex in our model. Then, we assume the VEGF_165_ binding to NRP1 alone is 20-fold weaker than binding to the NRP1:HSPG complex, according to a study showing that the presence of heparin significantly increases VEGF binding to NRP1^31^. Likewise, the rates for VEGFR2 coupling to VEGF_165_-bound NRP in the previous model are used for the VEGFR2 coupling to the VEGF_165_-bound NRP:HSPG complex in our model, while the previous rates for VEGF_165_-bound VEGFR2 coupling to NRP1 are used for the VEGF_165_-bound VEGFR2 coupling to NRP:HSPG complex in our model. To our knowledge, there are no available measurements to estimate the coupling rates of NRP1 to HSPG. Therefore, we assume the rates of NRP1 coupling to HSPG are the same as rates of VEGFR2 coupling to NRP1, which are taken from previous modeling^15^. Previous experimental study shows a VEGF_165_-mediated synergistic binding between NRP1 and heparin^30^, and we accordingly assume the coupling of VEGF_165_: NRP to HSPG and the coupling of VEGF_165_:HSPG to NRP are an order of magnitude stronger than the coupling between NRP and HSPG. Following previous works^15,18^, the coupling of VEGFR1 to NRP is set to be an order of magnitude weaker than VEGFR2-NRP coupling. According to the measured binding constants^30^, the coupling of VEGFR1 to HSPG is assumed to be 5-fold stronger than NRP-HSPG coupling.

For the TSP1 axis, we followed the values used in our previous works^19^. For the kinetic rates governing the FGF2 axis, we used the values estimated from experimental data in a previous study^32^. For the PF4 axis, the K_d_ values of PF4 binding to CXCR3 and LRP1 are estimated to be 1.85 nmol/l^34^ and 238 nmol/l^41^, respectively. These are used to set the dissociation rate, with the association rate held at 5×10^5^ M^-1^S^-1^, based on molecular dynamics studies of biomolecular reaction kinetics^42,43^. For the binding of angiogenic factors with iHSPG, we use the K_d_ values measured with heparin as estimations. The K_d_ values of heparin binding to VEGF_165_, FGF2, TSP1 and PF4 are 80, 39, 41 and 20 nmol/l^33,44–47^, respectively. The rates for FGF2 binding to HSPG on the cell surface (cHSPG) are estimated in previous work^32^. Based on the binding rates of FGF2 to cHSPG, we derive the rates of VEGF_165_, TSP1 and PF4 binding to cHSPG by scaling the FGF2-HSPG binding parameters according to their relative affinity to heparin, assuming the measured heparin affinity reflects their relative binding affinity to cHSPG. For the associations between pro- and anti-angiogenic factors, PF4 binds to VEGF_165_ and FGF2 with K_d_ values of 5 and 37 nmol/l^22^, respectively. The K_d_ values of TSP1 binding to VEGF and FGF2 are 10 and 10.8 nmol/^26,48^. The parameters for the protease activity are taken from our previous works^18,19^.

### Model implementation and simulation

The model ODEs are generated using BioNetGen^49^, a rule-based modeling framework. BioNetGen produces all possible molecular species and the corresponding ODEs by specifying a set of starting molecular species and defining reaction rules. Given 40 seed species and 127 reaction rules, the model produced by BioNetGen consists of 154 species. The set of 154 ODEs is implemented in MATLAB (The MathWorks, Natick, MA, USA), which we used to generate the dynamic results, as well as steady state predictions (i.e., when the model outputs change less than 0.01%). The MATLAB model file is provided in Supplementary File S3.

## 3 RESULTS

### 3.1 Baseline prediction of the angiogenic factors’ distribution in tumor tissue

The baseline secretion rates of angiogenic factors were tuned in order to obtain concentrations within the range of available experimental measurements. We report the predicted species’ concentrations (for VEGF, TSP1, PF4, FGF2, MMP3 and MMP9) and compare with experimental measurements in the Table 1.

With the baseline secretion rates, the model predicts that the pro-angiogenic factors (VEGF and FGF2) and anti-angiogenic factors (TSP1 and PF4) have significantly different distribution patterns in tumor tissue (Fig. 3). The majority of each pro-angiogenic factor in the tumor (~81% of VEGF and ~50% of FGF2) is bound to the cell surface, while only a small percentage of the anti-angiogenic factors (~16% of PF4 and ~12% of TSP1) exists on the cell surface. The cell surface bound ligands can be further categorized into non-signaling and signaling forms. The non-signaling forms include complexes with cHSPG and non-signaling receptors, and signaling forms include ligated receptors that promote intracellular signaling. Most of the cell-surface bound VEGF is in a signaling form, where VEGFR1-, VEGFR2-and NRP1-bound VEGF comprise 35%, 17% and 29% of the total VEGF in the tumor, respectively. Only 0.4% of total VEGF is bound to cHSPG. The model predicts that 23% of total FGF2 is in a signaling form bound to FGFR1:cHSPG dimers. The balance of the cell-surface FGF2 is non-signaling, bound to either cHSPG or FGFR1 monomers, which comprise 6% and 22% of total FGF2, respectively. In comparison to the distributions of the pro-angiogenic factors, most of the cell-surface bound anti-angiogenic factors are in non-signaling forms. The model predicts that 11% of total TSP1 and 16% of total PF4 are bound to cHSPG. Only 1% of total TSP1 is bound to signaling receptors, including CD47, CD36, LRP1 and β_1_. In the case of PF4, an even smaller fraction (0.4%) is bound to anti-angiogenic receptors, including CXCR3 and LRP1.

**Figure 3.**
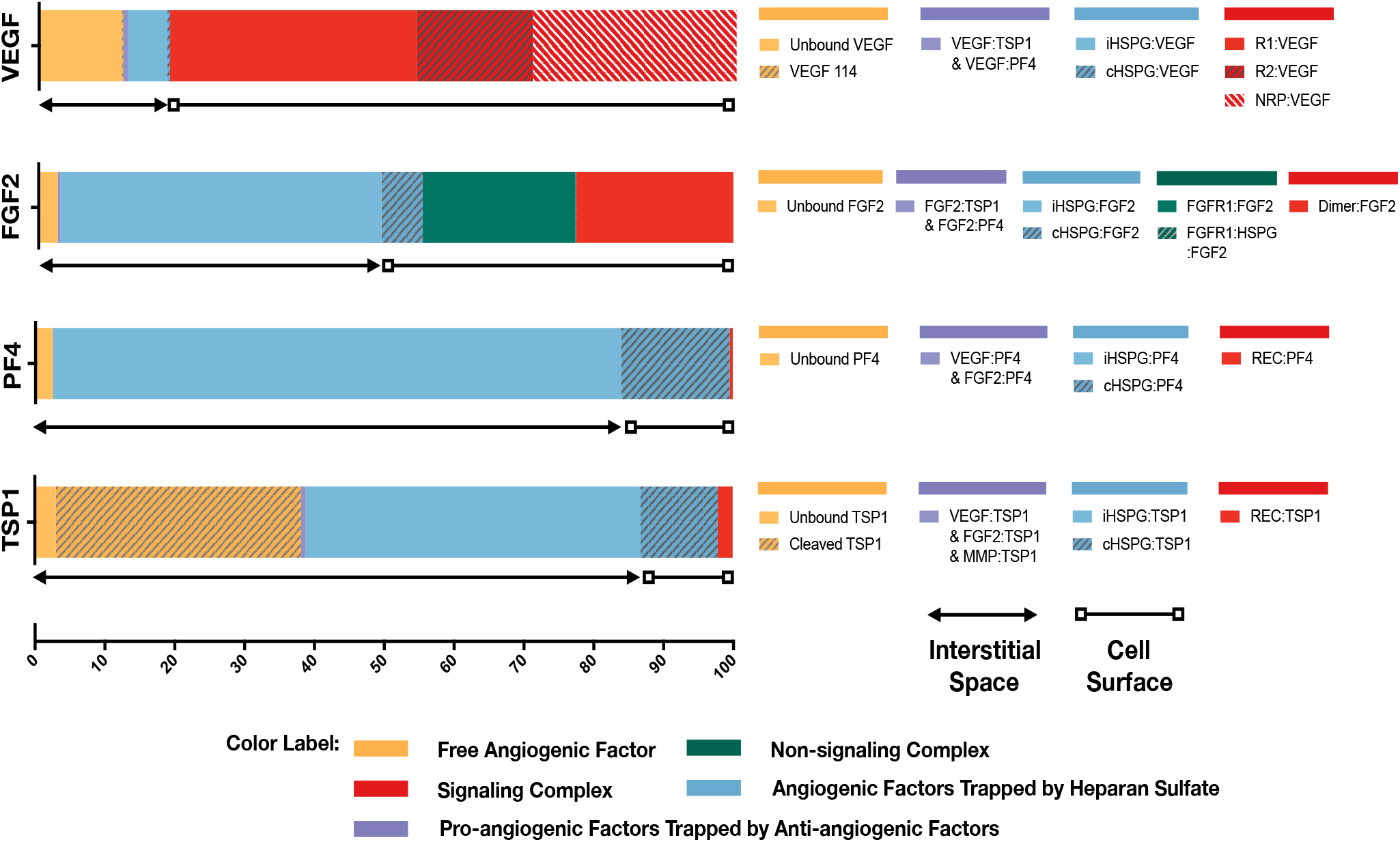
Distribution of VEGF, FGF2, TSP1 and PF4 in tumor tissue at steady state. The percentages of each angiogenic species in its various forms are shown. Species are grouped and labeled with different colors. The sum of the forms bound to the cell surface or in the interstitial space is also indicated.

In the interstitial space, there are three forms of angiogenic factors, including the unbound form, iHSPG-bound form, and the form bound to other angiogenic factors. Approximately 12% of total VEGF is in an unbound active form, including VEGF_121_ and VEGF_165_, and 0.1% of VEGF is present as the inactive isoform VEGF_114_. The percentages of VEGF bound to iHSPG or other angiogenic factors are 6% and 0.7%, respectively. Unlike VEGF, most FGF2 in the interstitial space is trapped by iHSPG. That is, 46% of total FGF2 is bound to iHSPG, while the unbound and angiogenic factor-bound forms only comprise 3% and 0.3% of the total FGF2, respectively. In contrast, the two anti-angiogenic factors, PF4 and TSP1, both have a larger portion in the interstitial space. In the case of PF4, 81% is bound to iHSPG, while the unbound and angiogenic factor-bound forms comprise only 2% and 0.01 % of the total PF4, respectively. Finally, most of TSP1 in the interstitial space is bound to iHSPG (48%) or in the cleaved, inactive form (35%). The balance of TSP1 is unbound or bound to other angiogenic factors, comprising 3% and 0.6% of the total TSP1, respectively

To summarize these results, the model predicts that most of VEGF and FGF2 is bound to the cell surface and in signaling forms, while most of TSP1 and PF4 is in the interstitial space and in non-signaling forms that are trapped by HSPGs or inactive due to proteolysis. It is worth noting that the fraction of the anti-angiogenic factors that is bound to pro-angiogenic factors only comprises a small percentage, which implies that direct binding between pro- and anti-angiogenic factors is not a major mechanism of the extracellular inhibition of pro-angiogenic signaling. Overall, this predicted distribution indicates a tumor state favoring pro-angiogenic signaling and neovascularization. In addition to the prediction under baseline secretion rates, we performed Monte Carlo simulations by sampling the secretion rates of VEGF, TSP1, PF4, FGF2, MMP3 and proMMP9 from a range of 100-fold below and 10-fold above the baseline values. The results (Supplementary Fig. S1) show that, even with potential uncertainty in the secretion rates, the main conclusions of the tumor distribution remain unchanged.

### 3.2 Secretion of anti-angiogenic factors modulates both pro- and anti-angiogenic signaling

To characterize the angiogenic state of the tumor, we defined the *angiogenic ratio*: the ratio of the concentrations of the pro-angiogenic signaling complexes to the anti-angiogenic signaling complexes. This ratio captures the activation level of pro-angiogenic receptors relative to the activation level of anti-angiogenic receptors. We examined how different angiogenic factors shift the angiogenic ratio (Fig. 4, column I) by varying the secretion rates of VEGF, FGF2, TSP1, and PF4 in a range of 100-fold below and 10-fold above the baseline values. We also predict how the concentrations of pro- and anti-angiogenic signaling complexes change in response to varying the secretion rates of the angiogenic factors (Fig. 4, columns I to V). Varying the secretion rates explores how targeting angiogenic factors changes the tumor angiogenic state, assuming 100-fold below represents strong inhibition and 10-fold above represents upregulation. We plot the angiogenic ratio and the concentrations of the angiogenic complexes normalized to the baseline secretion rates.

**Figure 4.**
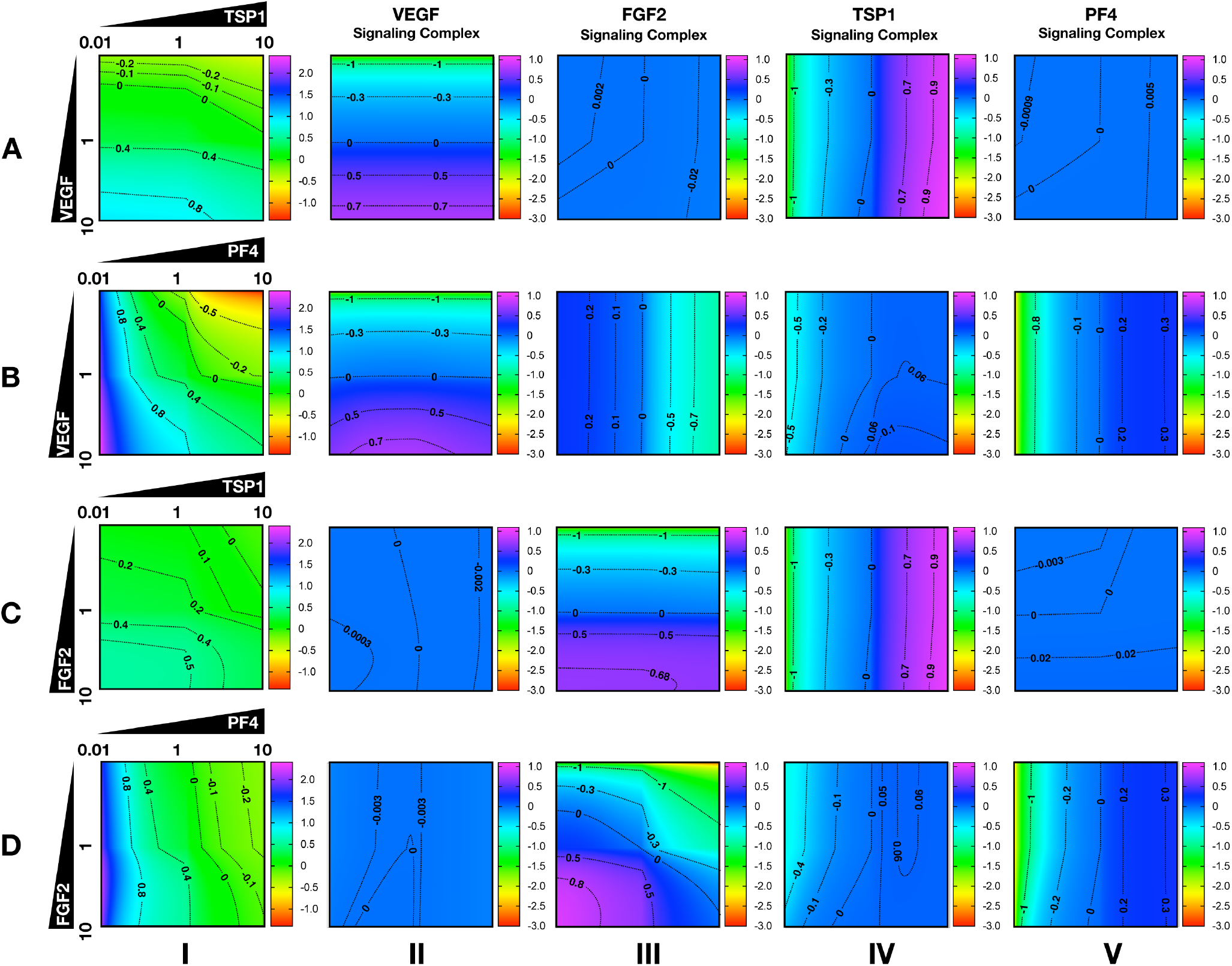
Effects of secretion of angiogenic factors on the angiogenic state of the tumor tissue. Column I shows the angiogenic ratio in the logio scale. Column II to V show the signaling complex levels (normalized to the baseline prediction) in the log_10_ scale. The value range is given by the colorbar. The horizontal and vertical axes of each subplot show the fold-change of the corresponding secretion rates, relative to their baseline values. The different rows show the effects of varying the secretion rates of different angiogenic factors: (A) VEGF and TSP1 secretion rates vary. (B) VEGF and PF4 secretion rates vary. (C) FGF2 and TSP1 secretion rates vary. (D) FGF2 and PF4 secretion rates vary. The predictions shown in the figures are based on the steady state of the system.

Higher secretion of the two pro-angiogenic factors shifts the angiogenic ratio by increasing the level of their corresponding pro-angiogenic complexes (Fig. 4). The gradient along the vertical axis in Fig. 4A and B, column I shows the angiogenic ratio will significantly increase with increasing VEGF secretion, which indicates that the tumor moves to a more pro-angiogenic state. The normalized level of the VEGF signaling complexes (Fig. 4A and B, column II) shows evident color changes along the vertical axis, which indicates the VEGF signaling is strongly enhanced with higher VEGF secretion. Meanwhile, the normalized level of FGF2, TSP1, and PF4 signaling complexes (Fig. 4A and B, columns III to V) shows no pronounced gradient along the vertical axis, implying that these signaling pathways are not affected by changing VEGF secretion.

Similarly, upregulating FGF2 secretion shifts the angiogenic ratio mainly through enhancing the formation of FGF2 pro-angiogenic complexes. The angiogenic ratio change along vertical axis in Fig. 4C, column I implies that the upregulation of FGF2 secretion increases the angiogenic ratio and promotes angiogenesis. The normalized level of the FGF2 complex (Fig. 4C and D, column III) significantly increase with increasing FGF2 secretion, while the concentration of the VEGF signaling complexes (Fig. 4C and D, column II) is highly stable when the FGF2 secretion is changed. The TSP1 and PF4 signaling complexes (Fig. 4D, columns IV to V) slightly change in response to FGF2, where increasing FGF2 secretion to a high level slightly promotes the formation of TSP1 bound and PF4 bound anti-angiogenic complexes.

Increasing the secretion of anti-angiogenic factors, particularly PF4, modulates the angiogenic ratio both by upregulating the levels of anti-angiogenic complexes and downregulating the pro-angiogenic complexes levels (Fig. 4). The gradient along the horizontal axis in Fig. 4A and C, column I indicates that increasing the secretion of TSP1 can decrease the angiogenic ratio. We also examined the change in the normalized levels of the angiogenic complexes. We found only TSP1 signaling complexes (Fig. 4A and C, column IV) show an evident color change along the horizontal axis in response to changing TSP1 secretion rates, which indicates that TSP1 secretion decreases the angiogenic ratio mainly through promoting the formation of TSP1-bound anti-angiogenic complexes. Model predictions show that changing PF4 secretion can strongly shift the angiogenic ratio (Fig. 4B and D, column I). In addition, increasing PF4 secretion promotes the formation of both TSP1- and PF4-bound anti-angiogenic complexes (Fig. 4B and D, columns IV to V). However, there also appears to be a limit to the effect of PF4, where PF4 does not continue to significantly promote the formation of TSP1 anti-angiogenic complexes when its secretion rate is higher than a certain level. Although varying PF4 secretion only slightly affects the formation of VEGF signaling complexes (Fig. 4B, column II), the color change along the horizontal axis in Fig. 4B and D, column III shows that increasing PF4 secretion can strongly inhibit the formation of FGF2 signaling complexes. Furthermore, the secretion of PF4 can nearly neutralize the effect of FGF2 secretion on the formation of FGF2 signaling complex (Fig. 4D, column III).

Overall, the model predicts that VEGF, FGF2 and TSP1 mainly bind to their own receptors to form more anti-angiogenic complexes and shift the angiogenic ratio, while PF4 affects the formation of signaling complexes of various angiogenic factors to change the angiogenic ratio.

### 3.3 Platelet factor 4 secretion can increase the levels of unbound pro-angiogenic factors in tumor

Increased secretion of PF4 is predicted to affect the formation of both anti- and pro-angiogenic signaling complexes. To get detailed insight into how PF4 modulates the distribution of other angiogenic factors, we report the change of specific signaling species upon varying the PF4 secretion rate (Fig. 5 and Supplementary Figure S2). For these simulations, the PF4 secretion rate is again varied in a range of 100-fold below and 10-fold above the baseline value. In the figures, the fold-change of the species on the vertical axis is the species’ concentration normalized to its concentration when PF4 secretion is 100-fold below the baseline value (the lower bound of the range over which the secretion rate was varied). Since PF4 mainly influences the other angiogenic factors by competing for the heparan sulfate binding sites, we investigated how the cHSPG level, a tumor-specific property, also affects the outcome of changing PF4 secretion. When we describe the cHSPG level below, we assume the baseline level as intermediate level. For low cHSPG levels, we ran simulations when cHSPG is 2-, 10- and 100-fold lower than the baseline level. For high cHSPG levels, we considered cHSPH levels 2- and 10-fold higher than the baseline level.

Although anti-angiogenic factors, PF4 and TSP1, can bind to pro-angiogenic factors, VEGF and FGF2, to sequester pro-angiogenic factors, our model predicts that the levels of unbound pro-angiogenic factors do not necessarily decrease in the presence of more anti-angiogenic factors (Fig. 5). Interestingly, varying PF4 secretion can significantly elevate the levels of unbound FGF2 and unbound VEGF in tumor tissue. For low cHSPG levels, increasing PF4 secretion may only slightly affect the levels of unbound FGF2 (Fig. 5A, column I; gray and blue lines), while unbound VEGF levels can decrease with increasing PF4 secretion (Fig. 5B, column I; blue lines). However, the secretion of PF4 strongly increases the level of unbound FGF2 if the tumor has intermediate to high cHSPG level (Fig. 5A, column I; red, orange, and black lines). Similarly, PF4 secretion can increase unbound VEGF when the cHSPG level is high (Fig. 5B, column I; red and orange lines). Examining the levels of specific isoforms of VEGF, we find that both unbound VEGF_165_ and unbound VEGF_121_ are affected. The change of VEGF_165_ is more pronounced (Fig. 5C, column I). Since the majority of unbound VEGF is VEGF_121_, the fold-change of VEGF highly resembles the change of VEGF_121_ (Fig. 5D, column I).

**Figure 5.**
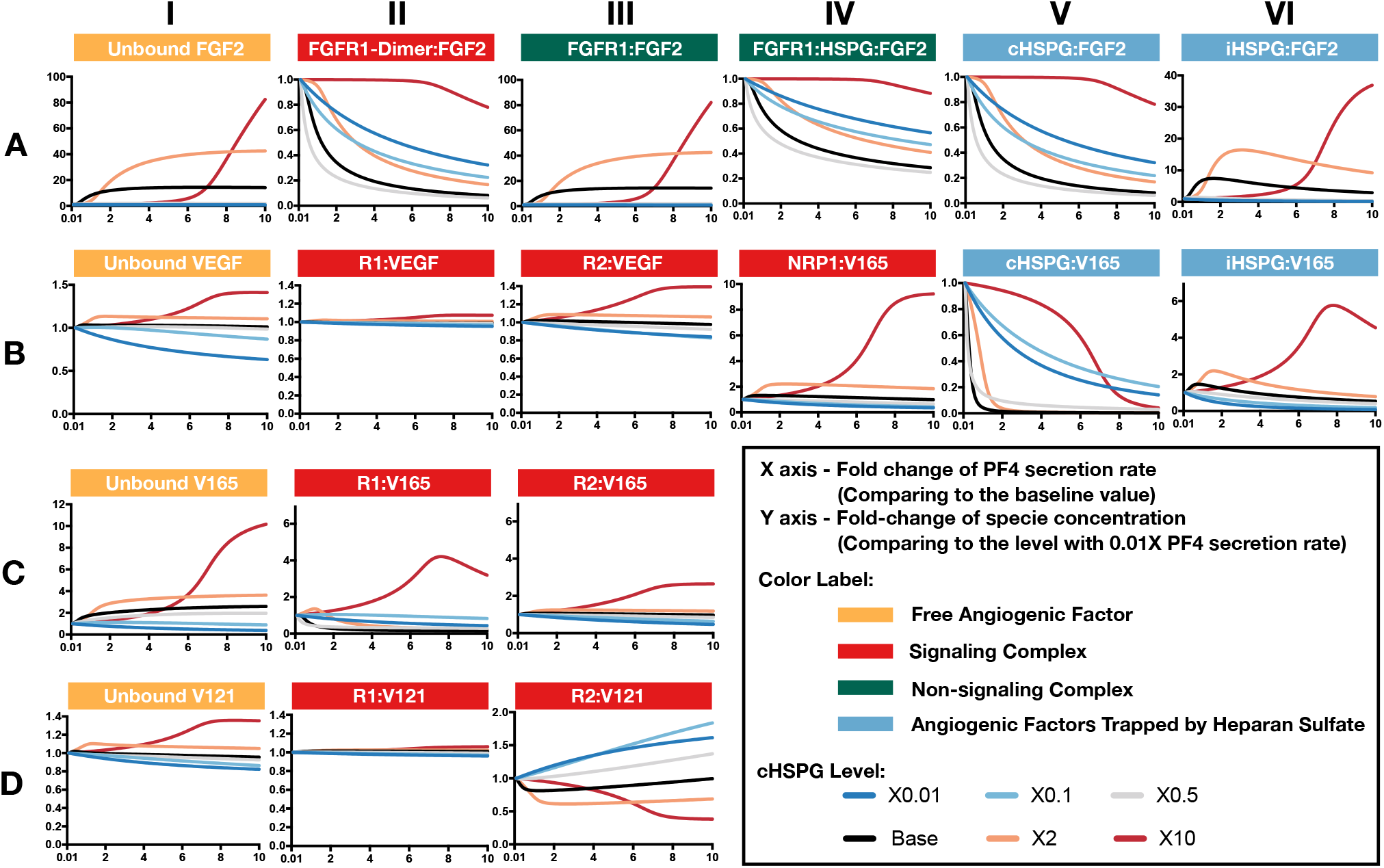
Effects of PF4 secretion on the formation of specific angiogenic complexes. (A) The change of species in the FGF2 axis. (B) The change of species in VEGF axis. (C) Change of VEGF_165_. (D) Change of VEGF_121_. The predictions shown in the figures are based on the steady state of the system.

The counterintuitive increase of unbound the pro-angiogenic factors with increasing PF4 secretion is caused by PF4 displacing pro-angiogenic factors from the cell surface heparan sulfate binding sites. PF4 preferentially competes for the HSPG on the cell surface first, causing cHSPG-bound VEGF and FGF2 to decrease with increasing PF4 secretion (Fig. 5A and B, column V). The decrement of cHSPG:FGF2 leads to a reduction of the trimeric complex FGFR1:HSPG:FGF2 (Fig. 5A, column IV) and the FGF2 signaling dimer (Fig. 5A, column II), which are only formed using cHSPG-bound FGF2. At the same time, the binding of PF4 to cHSPG reduces the availability of HSPG to bind to VEGF and VEGF receptors. Since the cHSPG affects VEGF binding to VEGFR2 and NRP1, the levels of VEGF-bound VEGFR2 and NRP1 change with increasing PF4 secretion. At high cHSPG level (Fig. 5B, columns III to IV; red and orange lines), increasing PF4 secretion promotes the formation of VEGF-bound VEGFR2 and NRP1. At low to intermediate cHSPG levels (Fig. 5B, columns III to IV; black, grey, light blue and dark blue lines), increasing PF4 secretion inhibits the formation of VEGF-bound VEGFR2 and NRP1. This switch is because of the biphasic response to cHSPG level, which will be explored in next section. Since the two isoforms of VEGF have different binding property to receptors, VEGF_165_ and VEGF_121_ bound to VEGFR2 show very different fold-changes in response to increasing PF4 secretion (Fig. 5C and D, column III). However, for both isoforms, varying PF4 secretion has differential effects on the levels of the pro-angiogenic ligated receptor complexes, depending on the cHSPG level.

iHSPG serves as a reservoir of angiogenic factors that can store and release pro-angiogenic factors. With increasing PF4 secretion, the pro-angiogenic factors displaced from cHSPG bind to iHSPG and form more FGF2- and VEGF-bound iHSPG (black, orange, and red lines in Fig. 5A and B, column VI). After the depletion of the available cHSPG, secreted PF4 competes for iHSPG binding sites, and iHSPG-bound PF4 significantly increases as the PF4 secretion rate increases. At high PF4 secretion rates, PF4 is even able to displace FGF2 and VEGF from iHSPG and reduce the iHSPG-bound pro-angiogenic factors (Fig. 5A and B, column VI; black, orange, and red lines).

In summary, the predictions show that the HSPG is an important mediator in how PF4 regulates pro-angiogenic factors. We found that, depending on the HSPG level, the secreted PF4 can displace more pro-angiogenic factors from the HS binding sites than the amount being sequestered, which eventually increases the level of unbound pro-angiogenic factors in the tumor interstitium.

### 3.4 VEGF signaling shows a biphasic response to the HSPG level and PF4 secretion rate

As presented above, the model predicts that the secretion of PF4 can increase the level of pro-angiogenic complexes on the cell surface. In the case of FGF2, the pro-angiogenic signaling complexes involving FGFR1 dimers decrease with increasing PF4 (Fig. 5A, column II) and the non-signaling complexes of ligated FGFR1 monomers increase with increasing PF4 secretion (Fig. 5A, column III). These signaling and nonsignaling forms of cell-surface FGF2 are also affected by cHSPG levels. Interestingly, for the VEGF axis, all three VEGF signaling complexes increase with increasing PF4 secretion, particularly for the high cHSPG condition (Fig. 5B, columns II to IV; red and orange lines). This indicates an activation of VEGF pro-angiogenic signaling caused by the anti-angiogenic factor PF4. To explain these results, we further investigate how cHSPG level and PF4 secretion modulate the signaling complexes for each of the angiogenic factors modeled in the tumor tissue (Fig. 6).

**Figure 6.**
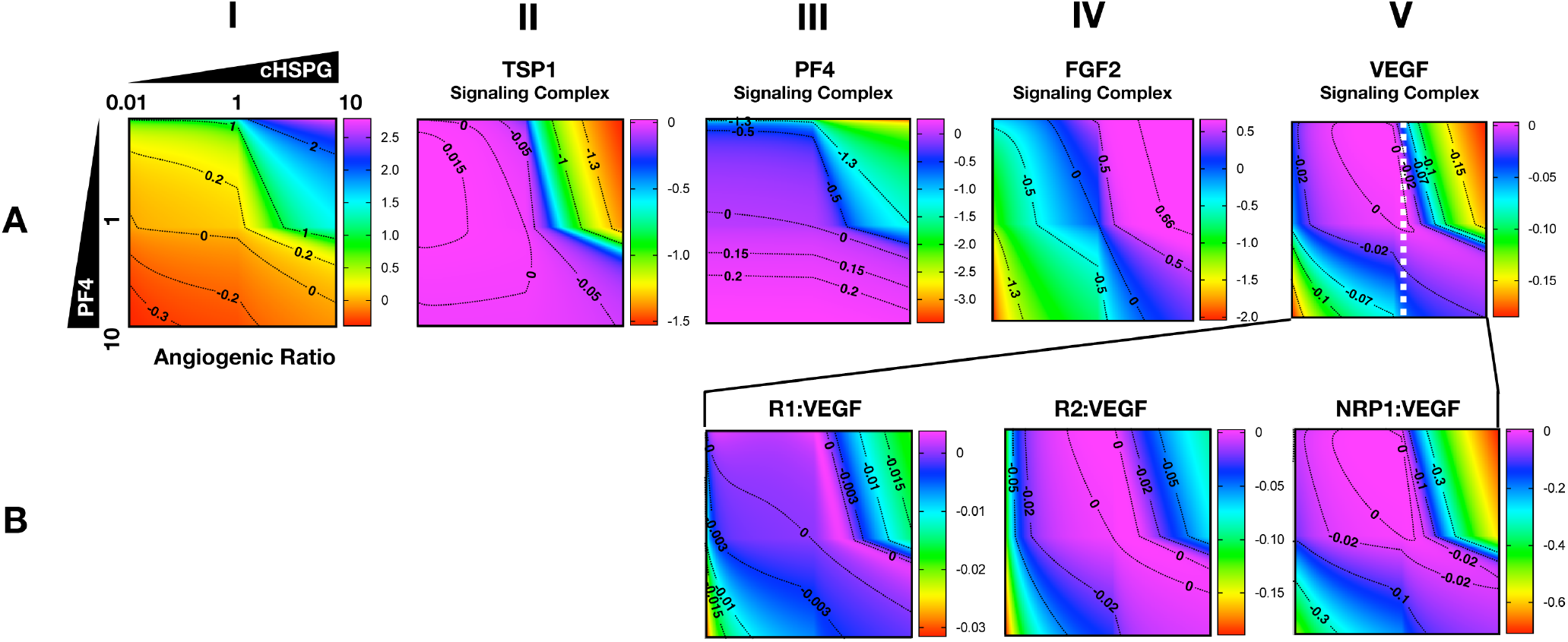
Impact of PF4 and cHSPG on the formation of specific angiogenic complexes. The horizontal axis of each subplot shows the fold-change of cHSPG level, relative the baseline value, and the vertical axes show the fold-change of PF4 secretion rate, relative to its baseline value. The value indicated by the colorbar is in the log_10_ scale. (A) The change of the angiogenic ratio and normalized levels of signaling complexes: Column I shows the change of the angiogenic ratio and column II to V show the changes of the normalized signaling complex levels (the values are normalized to the baseline prediction). (B) The change of the normalized levels of specific VEGF signaling complexes. The predictions shown in the figures are based on the steady state of the system.

As shown in Fig. 6A, column I, the model predicts that increasing cHSPG increases the angiogenic ratio (the tumor tissue is shifting to a more pro-angiogenic state) and increasing PF4 decreases the angiogenic ratio (shifting the tumor tissue to a less pro-angiogenic state). Together, these results indicate that HSPG promotes angiogenesis in tumor tissue, and the secretion of PF4 counteracts the pro-angiogenic effect of HSPG. Given the molecular detail of the model, we can explain these results. HSPG traps the two anti-angiogenic factors TSP1 and PF4. Thus, by increasing the HSPG level, the levels of TSP1 and PF4 signaling complexes are reduced (Fig. 6A, columns II and III). However, HSPG is needed for the formation of the pro-angiogenic FGF2 signaling dimers. Although the HSPG traps FGF2 as well, the predictions show that increasing HSPG increases the FGF2 signaling complexes (Fig. 6A, column IV). Additionally, increasing PF4 decreases FGF2 signaling complexes (Fig. 6A, column IV) by displacing FGF2 from cHSPG, as explained in the previous section.

In contrast, VEGF shows biphasic response to HSPG. The gradient along the horizontal axis in Fig. 6A, column V shows that the concentrations of the VEGF signaling complexes increase and then decrease with increasing HSPG level. Since PF4 competes for HSPG, increasing PF4 secretion decreases the HSPG availability to VEGF. Therefore, VEGF signaling also shows a biphasic response to PF4 secretion. For instance, at a medium HSPG level (dashed red line in Fig. 6A, column V), along the vertical axis, the color changes from blue to purple then back to blue, which means the concentrations of the VEGF signaling complexes go up and then back down with increasing PF4 secretion. The VEGF signaling complexes include of VEGFR1-, VEGFR2-, and NRP1-bound VEGF. In addition, different types of VEGF signaling complexes, including VEGFR1-, VEGFR2- and NRP1-bound VEGF complexes, show a biphasic response to HSPG and PF4 secretion (Fig. 6B, columns III and IV).

In summary, although PF4 secretion increases the unbound FGF2 level, the PF4 secretion strongly inhibits the formation of FGF2 signaling dimers that need HSPG to be formed. However, the VEGF signaling complexes can be formed through HSPG-dependent and HSPG-independent ways. Therefore, the VEGF signaling shows a biphasic response to the HSPG level. A low level HSPG limits the formation of VEGF signaling complexes through HSPG-dependent way. At the intermediate HSPG level, the VEGF signaling complexes reaches a peak level, while HSPG mainly traps VEGF and decreases VEGF signaling when it is present at a high level. Given the fact that PF4 secretion can efficiently limit HSPG availability, VEGF signaling shows a biphasic response to PF4 secretion as well. Therefore, at certain HSPG levels, the secretion of PF4 can enhance the VEGF pro-angiogenic signaling in tumor tissue.

### 3.5 The HSPG level affects the response to platelets activation and exogeneous PF4 therapy

Building on the simulations in which we vary the secretion rate of PF4, we apply the model to predict the effects of a local release of PF4 at the tumor site, mimicking PF4 release following platelet activation (where angiogenic factors are released) or a bolus injection of exogeneous PF4 as an antitumor therapy. The system is first allowed to reach steady state, which occurs in the first day. We then simulate two pulses of 5 mg PF4 per week, injected into the tumor interstitial space. This leads to a peak PF4 concentration of approximately 800 nM. The two pulses of PF4 occur at days 1 and 3.5. The release of PF4 follows an exponential decay with rate constant 2.8 ×10^-5^ s^-1^, assuming the PF4 are encapsulated in a biomaterial delivery vehicle^19^. We also perform the simulation at three different cHSPG levels to represent different tumor microenvironments: low (10-fold below the baseline value), medium (baseline value), and high (10-fold above the baseline). In this way, we examined how the tumor-specific property affects the response. Consistent with the results presented above, the model predictions reveal that depending on the HSPG level, platelet activation and recombinant PF4 can impact the pro-angiogenic signaling pathways in different ways.

The model predicts that cHSPG level significantly changes the response of VEGF signaling to the PF4 release (Fig. 7). In a tumor microenvironment with high HSPG, the release of PF4 in the tumor leads to an activation of VEGF signaling pathway. Specifically, the concentration of unbound VEGF increased from 128 pM to 177 pM (a 1.9-fold increase) after the release of PF4, and it goes back down due to the degradation of PF4 (Fig. 7A, red line). The levels of VEGFR1-, VEGFR2-, and NRP1-bound VEGF increase by 1.1-, 1.5-, and 8.1-fold, respectively (Fig. 7B-D, red line). However, in a microenvironment with medium HSPG level, the release of PF4 inhibits VEGF signaling (Fig. 7, black line). The concentrations of unbound VEGF and ligated VEGFR1 and VEGFR2 slightly decrease following the release of PF4, while NRP1-bound VEGF decreases 1.4-fold. In the tumor with low HSPG level, the release of PF4 shows a stronger inhibition, particularly for unbound VEGF and the VEGFR2 and NRP1 complexes (Fig. 7, blue line). The concentrations of unbound VEGF and VEGFR2-bound VEGF each decreased 1.2-fold, and NRP1-bound VEGF significantly decreased, by 2.3-fold.

**Figure 7.**
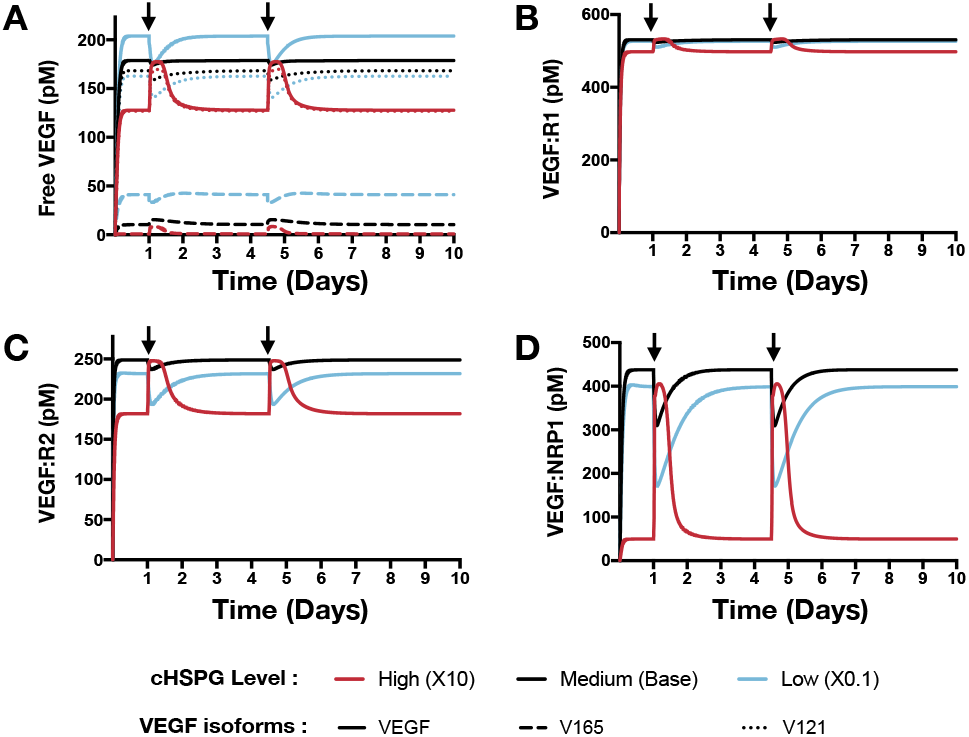
Effect of PF4 pulse release on VEGF signaling pathway. The system is allowed to reach a steady state, followed by two pulses of PF4. The start time of PF4 release is indicated by the arrows: at day 1 and day 3.5. The concentrations of species in the VEGF signaling pathway are predicted: (A) Concentration of free VEGF: Solid line, VEGF; Dashed line, V165; Dotted line, V121). (B) VEGF:VEGFR1 complexes. (C) VEGF:VEGFR2 complexes. (D) VEGF:NRP1 complexes.

In addition to affecting the VEGF signaling complexes, release of PF4 influences the FGF2 signaling complexes to different extents, depending on the tumor microenvironment (Fig. 8). Both unbound FGF2 and FGF2-bound FGFR1 complexes increase upon release of PF4 (Fig. 8A-B). However, the concentration of the trimeric complex HSPG:FGFR1:FGF2 significantly decreased following each PF4 pulse (Fig. 8C), which results in the reduction of FGF2-bound dimers. In a tumor with medium HSPG expression, the concentration of FGF2-bound dimers shows the largest decrease (6.5-fold). For low and high HSPG level, the FGF2-bound dimer concentration exhibits a 3.1- and 1.3-fold reduction, respectively.

**Figure 8.**
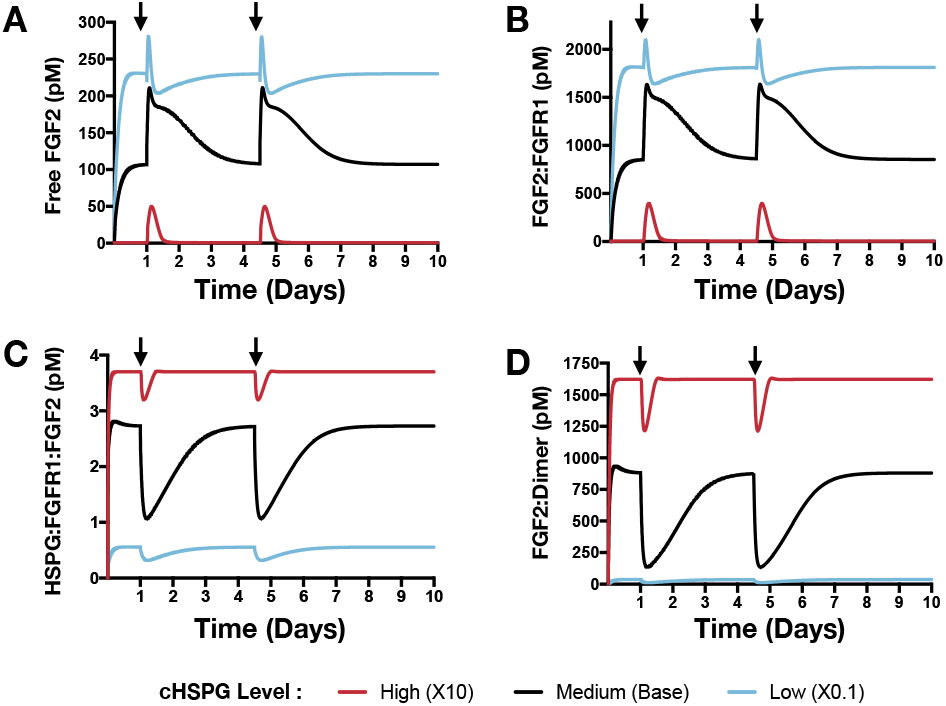
Effect of PF4 pulse release on FGF2 signaling pathway. The system is allowed to reach a steady state, followed by two pulses of PF4. The start time of PF4 release is indicated by the arrows: at day 1 and day 3.5. The concentrations of species in the FGF2 signaling pathway are predicted: (A) Concentration of free FGF2. (B) FGF2:FGFR1 complexes. (C) HSPG:FGFR1:FGF2 complexes. (D) HSPG:FGFR1:FGF2 dimers.

To summarize, we simulated relevant tumor scenarios in which the PF4 concentration would suddenly increase, such as following platelet activation or administration of exogenous PF4 as an anti-angiogenic treatment strategy. The model predicts that PF4 has differential effects on the concentrations of pro-angiogenic signaling complexes involving VEGF and FGF2, depending on the cell-surface level of HSPG. Particularly, at a high cHSPG level, PF4 is shown to have a counterintuitive effect of promoting the formation of pro-angiogenic VEGF complexes. Overall, these simulations demonstrate the utility of the modeling framework in understanding the possible outcomes of events that are physiologically relevant to tumor angiogenesis.

## 4 DISCUSSION

We present a novel systems biology model describing the distribution of two potent pro-angiogenic factors and two important anti-angiogenic factors in tumor tissue. This model significantly expanded previous works to enable a study of four relevant angiogenic factors. Our model considers their interactions with each other in the extracellular space of tumor tissue, which was missing in previous models. In addition, the model expansion allows us to investigate the impact of heparan sulfate proteoglycans (HSPG) on the angiogenic factors’ distribution. HSPG is an important modulator of tumor angiogenesis that is present on the cell surface, in the extracellular matrix, and in the cellular basement membranes. HSPG binds to and stores the angiogenic factors, facilitates the angiogenic factors’ signaling and mediates the extracellular interactions of pro- and anti-angiogenic factors. Thus, HSPGs are a vital part of the extracellular network of angiogenic factors. Although the role of HSPGs in FGF2 signaling has been modeled in several studies^32,33,50^, the impact of HSPGs on VEGF ligand binding has not been modeled explicitly before. We addressed this gap by incorporating knowledge reported in experimental studies of the synergistic binding of VEGF, its receptors and heparin^30^. With the expansions upon previous models, our work reports a new computational framework for a comprehensive study of the angiogenic regulation in the extracellular space of tumor tissue.

Given the molecular detail of the model, we gain mechanistic insight into the extracellular regulation of tumor angiogenic signaling. In the tumor extracellular space, TSP1 and PF4 are thought to regulate the formation of pro-angiogenic signaling complexes involving VEGF and FGF2 through two different mechanisms: sequestration – binding directly to VEGF and FGF2 to prevent binding to their pro-angiogenic receptors, and competition – competing for cell-surface HSPG to inhibit the formation of pro-angiogenic complexes. Our study shows that PF4 significantly inhibits pro-angiogenic signaling, mainly by competing for cell-surface HSPG binding sites, not through direct binding. Our model predicts that the majority of TSP1 is in a cleaved form owing to the action of proteases, and this cleaved form is inactive and unable to compete for cell-surface HSPG. Therefore, our predictions show that TSP1 does not strongly inhibit the formation of VEGF and FGF2 signaling complexes. Moreover, the measured binding affinities between the anti-angiogenic factors (TSP1 and PF4) and the pro-angiogenic factors (VEGF and FGF2) are much weaker than their affinities to the receptors, which explains that the binding between them cannot efficiently sequester the pro-angiogenic ligands.

Our model predicts possible counterintuitive outcomes for the angiogenic state of following the release of anti-angiogenic factors. The secretion of anti-angiogenic factors, PF4 and TSP1, is generally assumed to reduce the concentrations of the free pro-angiogenic factors and inhibit the formation of pro-angiogenic signaling complexes. However, our model predicts that increasing the secretion of PF4 in tumor tissue can lead to two counterintuitive results: an increase in interstitial FGF2 and VEGF levels (Results section 3.3) and greater formation of pro-angiogenic signaling complexes, particularly in the VEGF signaling pathway (Results section 3.4). The reason for the increased VEGF and FGF2 levels in the tumor interstitium following PF4 secretion is that PF4 competes for the HSPG binding sites in the cell surface, basement membrane and extracellular matrix, thereby releasing the pro-angiogenic factors from those sites and increasing level of free pro-angiogenic ligands. When this effect is stronger than the sequestration that occurs when PF4 binds directly to pro-angiogenic factors, the level of unbound VEGF and FGF2 will be higher compared to the tumor microenvironmental condition with lower PF4 secretion (Fig. 5 Column I). The greater formation of VEGF signaling complexes (which presumably will activate intracellular signaling) caused by PF4 is because of the intrinsic biphasic response to HSPG level (Fig. 6A Column V). At low levels, HSPG limits the formation of VEGF signaling complexes. When HSPG is present at an intermediate level, it promotes VEGF signaling by facilitating VEGF binding to receptors. However, when HSPG is at an even higher level, it traps VEGF and reduces the formation of VEGF signaling complexes, which leads to a low VEGF signaling again. Higher secretion of PF4 allows PF4 to more strongly compete for HSPG, which can alleviate the HSPG sequestration of VEGF and promote VEGF signaling in certain conditions.

These predicted counterintuitive results are clinically relevant for understanding the outcome of platelet activation and anti-angiogenic therapy. In the human body, PF4 is stored in platelet α-granules and released upon platelet activation. It is reported that the serum concentrations of PF4 exceeds 8 μg/mL (276 nM) during platelet activation^51–55^. Given the fact that platelets are attracted and accumulated at tumor sites^56^, it is possible that even higher concentrations of PF4 may be present in the local tumor microenvironment when platelet activation occurs. Besides the release of endogenous PF4 from platelets, recombinant PF4 (rPF4) has been studied as an anti-tumor therapeutic to prevent angiogenesis, showing efficacy in both *in vitro* and *in vivo* settings^57,58^. rPF4 was tested in a mouse model to inhibit tumor growth with a dosage at 0.1 μg/μL for 5 μg in total^57^, and rPF4 has been tested in patients with advanced colorectal carcinoma at a dosage of 3 mg/kg in 30-minute infusions^59^. Thus, the administration of rPF4 as a therapeutic agent will greatly increase the total PF4 level in the tumor. To mimic the local increase of PF4 concentration due to either platelet activation or a bolus injection of rPF4, we simulated a situation of controlled release of PF4 in the tumor interstitium. The simulation results highlight the impact of HSPGs level on the outcome of platelet activation and anti-angiogenic therapy (Results 3.5).

In addition, our model also has other practical applications, complementing pre-clinical and clinical studies. The model can be further expanded to a whole body model to study the clinically tested anti-angiogenic therapy and patient response, as we have done in previous work with VEGF and TSP1 modeling^18^. Therefore, our model provides a basis to study the anti-angiogenic therapy targeting multiple angiogenic factors, including VEGF, FGF2, TSP1 and PF4. In addition, the signaling complexes in our model are also the species initiating downstream signaling in previous published downstream signaling models^32,60,61^; therefore, our model can be connected with models of intracellular signaling to characterize the downstream signaling changes.

There are some limitations of our model that can be addressed in future work. Given the scarcity of the quantitative data, we used the measurements from tumor types other than breast cancer to tune the baseline value of the angiogenic factors secretion rates. Since there are no available measurements of the PF4 level directly from tumor tissue sample, we use the measured blood PF4 level in breast cancer patients as an estimation of tumor interstitial PF4 level. Additionally, HSPG includes various types, each with different masses and number and types of heparan sulfate chains^24^. This great complexity is difficult to fully characterize mathematically and warrants its own highly detailed mechanistic model. To make the model more useful, we made a simplification to only explicitly define two generic species of HSPGs that capture the two key HSPG classes with distinct functions, rather than a detailed description of all HSPG species. One of the types of HSPGs in the model is on the cell surface (cHSPG) that can bind to ligand, couple with receptors, and is subject to internalization. The other type is the interstitial HSPG (iHSPG) in the extracellular matrix and basement membranes, which only traps free angiogenic ligands and is not subject to degradation and internalization. We acknowledge that the soluble form of HSPG, such as heparin, is also important to consider in the context of the tumor^62^. However, we do not explicitly model this class of HSPGs, because its binding to ligands and receptors, as well as its degradation, makes it very similar to the cHSPG in the model. If needed, our model can be extended to include more types of HSPGs. Despite these limitations, our model provides relevant mechanistic insight into interactions between angiogenic factors, their receptors, and HSPGs.

In conclusion, in this study, we present a novel model to characterize the extracellular distribution of four important angiogenic factors: VEGF, FGF2, TSP1, and PF4. The model provides mechanistic insights into the regulation of the angiogenic interaction network in the extracellular space of tumor tissue. We expect that the insights generated by our model will enable a better understanding tumor angiogenesis interactions and aid the development of new anti-angiogenic therapy.

## ACKNOWLEDGEMENTS

The authors acknowledge the support of the US National Science Foundation (CAREER Award 1552065 to SDF), the American Cancer Society (130432-RSG-17-133-01-CSM to SDF) and the USC Provost’s PhD Fellowship (DL).

## SUPPORTING INFORMATION

**File S1**. List of reactions in the model

**File S2.** List of parameter values used in the model

**File S3.** MATLAB model file

