## Supplementary File S2 for "Exploring the extracellular regulation of the tumor angiogenic interaction network using a systems biology model"

**Model parameters: numerical values and definitions.**

| **Parameter** | **Description** | **Unit** | **Model Value** | **Reference and Notes** | |
| --- | --- | --- | --- | --- | --- |
| **Geometric Parameters** | | | | | |
| Mac Gabhann et al. constructed the tissue-based model of the VEGF system and described the derivations of following geometric parameters in their work^1^. Here we report the value and the sources of parameters used in the model file provided by us. | | | | | |
| ECM_conc | Binding site density of extracellular matrix | M | 7.5e-7 | ^2^ | * The Heparan Sulfate binding site density are measured with FGF binding. We assume the VEGF, TSP1, and PF4 are binding to these binding sites as well. |
| EBM_conc | Binding site density of the basement membrane surrounding the endothelial cells |  | 1.3e-5 | ^3^ |  |
| PBM_conc | Binding site density of the basement membrane surrounding the parenchymall cells |  | 1.3e-5 |  |  |
| ECM_Vol_tis_dis | Volume of extracellular matrix of which available to soluble species in breast tumor | cm^3^/cm^3^ tissue | 0.51931 |  |  |
| EBM_Vol_tis_dis | Volume of microvessel basement membrane of which available to soluble species in breast tumor |  | 0.00027 |  |  |
| PBM_Vol_tis_dis | Volume of tumor cells basement membrane of which available to soluble species in breast tumor |  | 0.002446 |  |  |
| vol_Tumor | Total volume of tumor tissue | cm^3^ | 33.51032 | ^4^ |  |
| tumorSA_Vol_tis_dis | Total tumor cells surface area in tumor tissue | cm^2^/cm^3^ tissue | 1534 | ^5^ |  |
| VesselSA_Vol_tis_dis | Total microvessels surface area in tumor tissue |  | 105 | ^6^ |  |
| tumorCellSurfArea_tis_dis | Surface area of one cancer cell | cm^2^ | 9.97e-6 | ^5^ |  |
| VesselCellSurfArea_tis_dis | Surface are of the abluminal side of an endotheali cell in tumor tissue |  | 1.00e-5 |  |  |
| **Baseline Secretion Rates** | | | | | |
| The secretion rates of VEGF, TSP1 and MMP are consistent with previous modeling works^1,7^. Two new newly introduced parameters, including the production rates of FGF2 and PF4, are estimated with experimental data (See Method section) in this work. | | | | | |
| qTSP1disEC | TSP1 secretion rate of endothelial cell in tumor tissue (abluminal side) | molecules/cell/s | 6 | ^7^ | * These secretion rates are estimated in our previous modeling work. The ratio of tumor cell secreted TSP1 to endothelial cell secreted TSP1 is set to be 1:10. |
| qTSP1tum | TSP1 secretion rate of tumor cell |  | 0.6 |  |  |
| qMMP3_disEC | MMP3 secretion rate of endothelial cell in tumor tissue |  | 180 |  |  |
| qproMMP9_disEC | proMMP9 secretion rate of endothelial cell in tumor tissue |  | 330 |  |  |
| qV165_tumor | VEGF165 secretion rate of tumor cell |  | 0.387 | ^8^ | * The tumor secretion rates were estimated ^9^ with experimental measurements ^10–13^ . The ratio of tumor secreted VEGF165 to VEGF121 is set to be 1:1.  * The ratio of tumor endothelial cell secreated VEGF165 to VEGF121 is set to 9:1 ^14^. |
| qV121_tumor | VEGF121 secretion rate of tumor cell |  | 0.387 |  |  |
| qV165_disEC | VEGF165 secretion rate of endothelial cell in tumor compartment |  | 0.0324 |  |  |
| qV121_disEC | VEGF121 secretion rate of endothelial cell in tumor compartment |  | 0.0324*10/90 |  |  |
| qFGF2 | FGF2 production rate in tumor | (mol/cm^3^ tissue)^-1^s^-1^ | 3e-16 |  | *The production rates of these two angiogenic factors are tuned to match the experimental data in this work. |
| qPF4 | PF4 production rate in tumor |  | 1e-14 |  |  |
| **Recycling of the receptors** | | | | | |
| The recycling rates of receptors are originally from the first tumor tissue model of VEGF-Receptor System by Gabhann et al^5^. | | | | | |
| sR_receptors | Recycling rate of unbound receptors | s-1 | 0.00028 | ^5^ | *We assume all receptors have same internalization and recycling rates. |
| k_int_receptors | Internalization rate of all ligated and unbound receptors |  | 0.00028 |  |  |
| **Kinetic Parameters** | | | | | |
| Mac Gabhann et al applied following kinetic rates of VEGF system in their work of tumor tissue model^5^ and illustrated the conversion from in vitro parameters to tissue parameters basing on geometric parameters. Following reported values are in vitro parameters, which are converted to tissue parameters used by model with the same strategy. | | | | | |
| kon_FGF2_GAG | FGF2 binding to interstitial HSPG | M^-1^s^-1^ | 8600 |  | *Assumed to have same association rate as VEGF-Heparin binding. |
| kon_FGF2_HSGAG | FGF2 binding to cell surface HSPG |  | 3.66e+6 | ^15^ |  |
| kon_FGF2_FGFR1 | FGF2 binding to FGFR1 monomer |  | 4.91e+5 |  |  |
| kon_FGF2_TSP1 | FGF2 binding to TSP1 |  | 1.89e+4 | ^16^ |  |
| kon_PF4_GAG | PF4 binding to interstitial HSPG |  | 8600 |  | *Assumed to have same association rate as VEGF-Heparin binding. |
| kon_PF4_HSGAG | PF4 binding to cell surface HSPG |  | 3.66e+6 |  | * Assumed to have same association rate as FGF2-cHSPG binding. |
| kon_PF4_CXCR3 | PF4 binding to CXCR3 |  | 5.00e+5 | ^17^ | * Association rates are set to fixed constant (See method section). |
| kon_PF4_LRP1 | PF4 binding to LRP1 |  | 5.00e+5 | ^18^ |  |
| kon_PF4_V165 | PF4 binding to VEGF_165_ |  | 5.00e+5 | ^19^ |  |
| kon_PF4_FGF2 | PF4 binding to FGF2 |  | 5.00e+5 | ^20^ |  |
| kon_TSP1_GAG | TSP1 binding to interstitial HSPG |  | 8600 |  | *Assumed to have same association rate as VEGF-Heparin binding. |
| kon_TSP1_HSGAG | TSP1 binding to cell surface HSPG |  | 3.66e+6 |  | * Assumed to have same association rate as FGF2-cHSPG binding. |
| kon_TSP1_CD36 | TSP1 binding to CD36 |  | 5.00e+5 | ^7^ | *The kinetic paramters of TSP1-receptor system are estimated in our previous tumor tissue model of VEGF and TSP1, which included detailed derivations. |
| kon_TSP1_CD47 | TSP1 binding to CD47 |  | 5.00e+5 |  |  |
| kon_TSP1_LRP1 | TSP1 binding to LRP1 |  | 2.10e+5 |  |  |
| kon_TSP1_B1 | TSP1 binding to β1 integrin |  | 5.00e+5 |  |  |
| kon_TSP1_VEGF | TSP1 binding to VEGF |  | 5.00e+5 |  |  |
| kon_TSP1_MMP3 | TSP1 binding to MMP3 |  | 1.00e+5 |  |  |
| kon_V165_GAG | VEGF_165_ binding to interstitial HSPG |  | 8600 | ^21–23^ |  |
| Kon_V165_HSGAG | VEGF_165_ binding to cell surface HSPG |  | 3.66e+6 |  | * Assumed to have same association rate as FGF2-cHSPG binding. |
| kon_V165_N1 | VEGF_165_ binding to Neuropilin-1 |  | 3.20e+6 | ^24,25^ |  |
| kon_V165_N1H | VEGF_165_ binding to Neuropilin-1 coupled with HSPG |  | 3.20e+6 |  |  |
| kon_V165_R1 | VEGF_165_ binding to VEGFR1 |  | 3.00e+7 | ^26^ |  |
| kon_V165_R2 | VEGF_165_ binding to VEGFR2 |  | 1.00e+7 | ^27,28^ |  |
| kon_V121_R1 | VEGF_121_ binding to VEGFR1 |  | 3.00e+7 | ^8^ | *Set to the same as V165- VEGR receptor |
| kon_V121_R2 | VEGF_121_ binding to VEGFR2 |  | 1.00e+7 |  |  |
| kon_MMP9_LRP1 | MMP9 binding to LRP1 |  | 9245 | ^7^ |  |
| kon_MMP3_proMMP9 | MMP3 binding to proMMP9 |  | 10000 |  |  |
| koff_FGF2_GAG | FGF2 binding to interstitial HSPG | s^-1^ | 6.9e-4/80*39 |  | *Scaled according to VEGF-Heparin binding dissociation rate. |
| koff_FGF2_HSGAG | FGF2 binding to cell surface HSPG |  | 1.11e-3 | ^15^ |  |
| koff_FGF2_FGFR1 | FGF2 binding to FGFR1 monomer |  | 2.36e-4 |  |  |
| koff_FGF2_TSP1 | FGF2 binding to TSP1 |  | 2.05e-4 | ^16^ |  |
| koff_PF4_GAG | PF4 binding to interstitial HSPG |  | 6.9e-4/80*20 |  | *Scaled according to VEGF-Heparin binding dissociation rate. |
| koff_PF4_HSPG | PF4 binding to cell surface HSPG |  | 1.11e-3/39*20 |  | *Scaled according to FGF2-cHSPG binding dissociation rate. |
| koff_PF4_CXCR3 | PF4 binding to CXCR3 |  | 9.25e-4 | ^17^ | * Measured Kd values are used to set dissociation rates accordingly (See method section). |
| koff_PF4_LRP1 | PF4 binding to LRP1 |  | 0.119 | ^18^ |  |
| koff_PF4_V165 | PF4 binding to VEGF_165_ |  | 2.5e-3 | ^19^ |  |
| koff_PF4_FGF2 | PF4 binding to FGF2 |  | 0.016 | ^20^ |  |
| koff_TSP1_GAG | TSP1 binding to interstitial HSPG |  | 6.9e-4/80*41 |  | *Scaled according to VEGF-Heparin binding dissociation rate. |
| koff_TSP1_HSGAG | TSP1 binding to cell surface HSPG |  | 1.11e-3/39*41 |  | *Scaled according to FGF2-cHSPG binding dissociation rate. |
| koff_TSP1_CD36 | TSP1 binding to CD36 |  | 0.115 | ^7,29,30^ |  |
| koff_TSP1_CD47 | TSP1 binding to CD47 |  | 0.005 |  |  |
| koff_TSP1_LRP1 | TSP1 binding to LRP1 |  | 0.0025 |  |  |
| koff_TSP1_B1 | TSP1 binding to β1 integrin |  | 0.05 |  |  |
| koff_TSP1_VEGF | TSP1 binding to VEGF |  | 0.005 |  |  |
| koff_TSP1_MMP3 | TSP1 binding to MMP3 |  | 0.0022303 |  |  |
| koff_V165_GAG | VEGF_165_ binding to interstitial HSPG |  | 6.9e-4 | ^21–23^ |  |
| koff_V165_HSGAG | VEGF_165_ binding to cell surface HSPG |  | 1.11e-3/39*80 |  | *Scaled according to FGF2-cHSPG binding dissociation rate. |
| koff_V165_N1 | VEGF165 binding to Neuropilin-1 |  | 0.020 | ^31^ | *Set to be 20-fold weaker than VEGF_165_ binding to HSPG coupled NRP1 |
| koff_V165_N1H | VEGF165 binding to Neuropilin-1 coupled with HSPG |  | 0.001 | ^24,25^ |  |
| koff_V165_R1 | VEGF165 binding to VEGFR1 |  | 0.001 | ^26^ |  |
| koff_V165_R2 | VEGF165 binding to VEGFR2 |  | 0.001 | ^27,28^ |  |
| koff_V121_R1 | VEGF121 binding to VEGFR1 |  | 0.001 | ^8^ |  |
| koff_V121_R2 | VEGF121 binding to VEGFR2 |  | 0.001 |  |  |
| koff_MMP9_LRP1 | MMP9 binding to LRP1 |  | 0.00049 | ^7^ |  |
| koff_MMP3_proMMP9 | MMP3 binding to proMMP9 |  | 0.001 |  |  |
| kc_N_H | Coupling of Neuropilin and HSPG | (mol/cm^2^)^-1^s^-1^ | 3.10e+13 |  | *Assumed to be same as V165R2-NH coupling. |
| kc_V165N_H | Coupling of VEGF_165_ bound Neuropilin and HSPG |  | 3.10e+13 |  |  |
| kc_V165H_N | Coupling of Neuropilin and VEGF_165_ bound HSPG |  | 3.10e+13 |  |  |
| kc_V165NH_R2 | Coupling of VEGFR2 and Neuropilin |  | 1.00e+14 | ^32,33^ | *Estimated in ^32^ using data from ^33^. |
| kc_V165R2_NH | Coupling of VEGFR2 and Neuropilin |  | 3.10e+13 |  |  |
| kc_R1_N | Coupling of VEGFR1 and Neuropilin |  | 1.00e+14 | ^8^ | *Set to the same as R2-N receptor |
| kc_R1_H | Coupling of VEGFR1 and HSPG |  | 3.10e+13 |  | *Set to the same as V165R2-NH coupling. |
| kc_CD36_R2 | Coupling of CD36 and VEGFR2 |  | 3.10e+11 | ^7^ |  |
| kc_CD36_B1 | Coupling of CD36 and  β1 integrin |  | 3.10e+13 |  |  |
| kc_CD47_R2 | Coupling of CD47 and VEGFR2 |  | 3.10e+11 |  |  |
| kc_H_FGFR1 | Coupling of HSPG and FGFR1 monomer |  | 3.96e+11 | ^15^ |  |
| kc_FGFR1_FGFR1 | Dimerization of HSPG bound FGFR1 |  | 3.17e+17 |  |  |
| kdissoc_N_H | Coupling of Neuropilin and HSPG | s^-1^ | 0.001 |  |  |
| kdissoc_V165N_H | Coupling of VEGF_165_ bound Neuropilin and HSPG |  | 1e-4 |  | *Assumed to be an order tighter than the coupling without the presence of VEGF_165_^34^. |
| kdissoc_V165H_N | Coupling of Neuropilin and VEGF_165_ bound HSPG |  | 1e-4 |  |  |
| kdissoc_R2_NH | Coupling of VEGFR2 and Neuropilin |  | 0.001 | ^32,33^ |  |
| kdissoc_R1_N | Coupling of VEGFR1 and Neuropilin |  | 0.01 | ^8^ | *Assumed to be slower dissociation than R2-N. |
| kdissoc_R1_H | Coupling of VEGFR1 and HSPG |  | 0.001/5 |  | *Set to be 5 folder stronger than HSPG-Neuropilin binding^34^. |
| kdissoc_CD36_B1 | Coupling of CD36 and  β1 integrin |  | 0.001 | ^7^ |  |
| kdissoc_CD36_R2 | Coupling of CD36 and VEGFR2 |  | 0.001 |  |  |
| kdissoc_CD47_R2 | Coupling of CD47 and VEGFR2 |  | 0.001 |  |  |
| kdissoc_H_FGFR1 | Coupling of HSPG and FGFR1 monomer |  | 3.24e-7 | ^15^ |  |
| kdissoc_FGFR1_FGFR1 | Dimerization of HSPG bound FGFR1 |  | 5.83e-4 |  |  |
| **Degradation and Clearance Rates** | | | | | |
| The degradation rates and clearance rates are estimated by conversion from reported half-life time in literatures. The cleavage rate of TSP1 are fitted to match experimental data in our previous work^7^. The catalytic rate of the activation of proMMP9 by MMP3 and the cleavage of VEGF165 by MMP were previously reported in modeling works by Vempati^29,30^. | | | | | |
| kdeg_VEGF | Degradation rate of VEGF | s^-1^ | 1.93e-4 | ^35^ |  |
| kdeg_FGF2 | Degradation rate of FGF2 |  | 1.93e-4 |  | *Set to be same as VEGF. |
| kdeg_PF4 | Degradation rate of PF4 |  | 0.0023 |  | *The clearance of PF4 was reported to be so rapid that no valid estimation of half-life can be made^36^. Here, we use 5 mins as the estimation^37^. |
| kdeg_TSP1 | Degradation rate of TSP1 |  | 3.3e-4 | ^7^ |  |
| kdeg_MMP | Degradation rate of MMP |  | 0.0012 |  |  |
| k_TSP1cleave | The cleavage rate of TSP1 through proteolysis |  | 0.00386 | ^7,29,30^ |  |
| k_act_MMP3_proMMP9 | A Michaelis-Menten Activation constant of the activation of MMP9 by MMP3 |  | 0.0019 |  |  |
| kp_mmp | The proteolysis rate of VEGF by MMPs | (mol/l)^-1^s^-1^ | 631 |  |  |
| **Receptor Numbers** | | | | | |
| The density of VEGF receptors and co-receptors on endothelial and tumor cells are systematically reported in our previous work^9^, which are taken from in vitro and in vivo measurements using quantitative flow cytometry^38^. We used the reported qualitative data in Human Protein Atlas to estimate the values for TSP1 and PF4 receptors as mentioned in Method Section. The numbers of receptors on the endothelial cell are set to be half of that for tumor cells, assuming equal distribution on the luminal and abluminal surfaces. | | | | | |
| CD36_number_tum | CD36 receptor number on tumor cell | receptors/cell | 2500 | ^7^ |  |
| CD47_number_tum | CD47 receptor number on tumor cell |  | 10000 |  |  |
| LRP1_number_tum | LRP1 receptor number on tumor cell |  | 5000 |  |  |
| B1_number_tum | β1 integrin number on tumor cell |  | 10000 |  |  |
| CD36_number_disEC | CD36 receptor number on tumor endothelial cell |  | 1250 |  |  |
| CD47_number_disEC | CD47 receptor number on tumor endothelial cell |  | 5000 |  |  |
| LRP1_number_disEC | LRP1 receptor number on tumor endothelial cell |  | 2500 |  |  |
| B1_number_disEC | β1 integrin receptor number on tumor endothelial cell |  | 5000 |  |  |
| R1_number_tum | VEGFR1 receptor number on tumor cell |  | 1100 | ^9^ | *VEGF receptor density followed our previous studies which uses the in vivo and in vitro measurements using quantitative flow cytometry. |
| R2_number_tum | VEGFR2 receptor number on tumor cell |  | 550 |  |  |
| N1_number_tum | Neuropilin-1 receptor number on tumor cell |  | 39500 |  |  |
| R1_number_disEC | VEGFR1 receptor number on tumor endothelial cell |  | 3750 |  |  |
| R2_number_disEC | VEGFR2 receptor number on tumor endothelial cell |  | 300 |  |  |
| N1_number_disEC | Neuropilin-1 receptor number on tumor endothelial cell |  | 20000 |  |  |
| FGFR1_number_tum | FGFR1 receptor number on tumor cell |  | 20000 | ^39^ |  |
| HSGAG_number_tum | HSPG number on tumor cell surface |  | 100000 |  |  |
| FGFR1_number_disEC | FGFR1 receptor number on tumor endothelial cell |  | 10000 |  |  |
| HSGAG_number_disEC | HSPG number on tumor endothelial cell surface |  | 50000 |  |  |
| CXCR3_number_tum | CXCR3 receptor number on tumor cell |  | 2500 |  | *Estimated according the qualitative data shown in Human Protein Atlas. |
| CXCR3_number_disEC | CXCR3 receptor number on tumor endothelial cell |  | 1250 |  |  |

* Here, M = moles/liter of interstitial fluid available to soluble species.
