## Supplementary File S3 for "Exploring the extracellular regulation of the tumor angiogenic interaction network using a systems biology model": Baseline_distribution_Run.m

```
clear all;
%%%%%%%%%%%%%%%%%%%%%%%%%%%%%%%%%%%%%%%%%%%%%%%%%%%%%%%%%%%%%%%%%%%
% This is the driver file to run the tissue model
% and predict the baseline distribution of VEGF, FGF2, TSP1 and PF4
%%%%%%%%%%%%%%%%%%%%%%%%%%%%%%%%%%%%%%%%%%%%%%%%%%%%%%%%%%%%%%%%%%%


numRuns = 1;

%specify the paramters for the model
for i = 1:numRuns
        modelParams(1,1) = 70;          %BW
        modelParams(2,1) = 6.02E+23;	%Avogadro
        
        modelParams(3,1) = 7.5e-7;      %ECM_conc      
        modelParams(4,1) = 1.3e-5;      %EBM_conc
        modelParams(5,1) = 1.3e-5;      %PBM_conc
        
        modelParams(6,1) = 0.51931;     %ECM_Vol_tis_dis
        modelParams(7,1) = 0.00027;     %EBM_Vol_tis_dis
        modelParams(8,1) = 0.002446;	%PBM_Vol_tis_dis
        
        
        modelParams(9,1) = 33.51032;   %vol_Tumor
        
        modelParams(10,1) = 1534;       %tumorSA_Vol_tis_dis
        modelParams(11,1) = 105;        %VesselSA_Vol_tis_dis
    
        modelParams(12,1) = 9.97E-06;	%tumorCellSurfArea_tis_dis
        modelParams(13,1) = 1.00E-05;	%VesselCellSurfArea_tis_dis
        
  
        modelParams(14,1) = 6;          %qTSP1disEC
        modelParams(15,1) = 0.6;        %qTSP1tum
        modelParams(16,1) = 180;        %qMMP3_disEC
        modelParams(17,1) = 330;        %qproMMP9_disEC
        modelParams(18,1) = 0;          %qMMP3tum
        modelParams(19,1) = 0;          %qproMMP9tum
        modelParams(20,1) = 0.387;      %qV165_tumor
        modelParams(21,1) = 0.387;      %qV121_tumor
        modelParams(22,1) = 0.0324;     %qV165_disEC
        modelParams(23,1) = 0.0324/9;   %qV121_disEC
        modelParams(24,1) = 3e-16;      %qFGF2
        modelParams(25,1) = 1e-14;      %qPF4

       
        modelParams(26,1) = 0.00028;	%sR_receptors
        modelParams(27,1) = 0.00028;	%k_int_receptors
       
        modelParams(28,1) = 3.66E+06;	%kon_FGF2_HSGAG
        modelParams(29,1) = 8600;       %kon_FGF2_GAG
        modelParams(30,1) = 4.91E+05;	%kon_FGF2_FGFR1
        modelParams(31,1) = 1.89E+04;	%kon_FGF2_TSP1
        modelParams(32,1) = modelParams(28,1); %kon_PF4_HSGAG
        modelParams(33,1) = 8600;       %kon_PF4_GAG
        modelParams(34,1) = 5e+05;      %kon_PF4_CXCR3
        modelParams(35,1) = 5e+05;      %kon_PF4_LRP1
        modelParams(36,1) = 5e+05;      %kon_PF4_V165
        modelParams(37,1) = 5e+05;      %kon_PF4_FGF2
        modelParams(38,1) = modelParams(28,1); %kon_TSP1_HSGAG
        modelParams(39,1) = 8600;       %kon_TSP1_GAG
        modelParams(40,1) = 5.00E+05;	%kon_TSP1_CD36
        modelParams(41,1) = 5.00E+05;	%kon_TSP1_CD47
        modelParams(42,1) = 2.10E+05;	%kon_TSP1_LRP1
        modelParams(43,1) = 5.00E+05;	%kon_TSP1_B1
        modelParams(44,1) = 5.00E+05;	%kon_TSP1_VEGF
        modelParams(45,1) = 1.00E+05;	%kon_TSP1_MMP3
        modelParams(46,1) = modelParams(28,1); %kon_V165_HSPG
        modelParams(47,1) = 8600;       %kon_V165_GAG
        modelParams(48,1) = 3.20E+06;	%kon_V165_N1
        modelParams(49,1) = 3.20E+06;	%kon_V165_N1H
        modelParams(50,1) = 3.00E+07;	%kon_V165_R1
        modelParams(51,1) = 1.00E+07;	%kon_V165_R2  
        modelParams(52,1) = 3.00E+07;	%kon_V121_R1
        modelParams(53,1) = 1.00E+07;	%kon_V121_R2
        modelParams(54,1) = 9245;	    %kon_MMP9_LRP1
        modelParams(55,1) = 10000;      %kon_MMP3_proMMP9
       
        modelParams(56,1) = 1.11E-03;   %koff_FGF2_HSGAG
        modelParams(57,1) = 6.9E-04/80*39;    %koff_FGF2_GAG
        modelParams(58,1) = 2.36E-04;   %koff_FGF2_FGFR1
        modelParams(59,1) = 0.000205;   %koff_FGF2_TSP1
        modelParams(60,1) = modelParams(56,1)/39*20; %koff_PF4_HSGAG
        modelParams(61,1) = 6.9E-04/80*20;    %koff_PF4_GAG
        modelParams(62,1) = 9.25e-4;      %koff_PF4_CXCR3
        modelParams(63,1) = 0.119;      %koff_PF4_LRP1
        modelParams(64,1) = 2.5e-3;     %koff_PF4_V165
        modelParams(65,1) = 0.016;      %koff_PF4_FGF2
        modelParams(66,1) = modelParams(56,1)/39*41; %koff_TSP1_HSGAG
        modelParams(67,1) = 6.9E-04/80*41;    %koff_TSP1_GAG
        modelParams(68,1) = 0.115;      %koff_TSP1_CD36
        modelParams(69,1) = 0.005;      %koff_TSP1_CD47
        modelParams(70,1) = 0.0025;     %koff_TSP1_LRP1
        modelParams(71,1) = 0.05;       %koff_TSP1_B1
        modelParams(72,1) = 0.005;      %koff_TSP1_VEGF
        modelParams(73,1) = 0.0022303;	%koff_TSP1_MMP3
        modelParams(74,1) = modelParams(56,1)/39*80;   %koff_V165_HSPG
        modelParams(75,1) = 6.9E-04;    %koff_V165_GAG
        modelParams(76,1) = 0.020;      %koff_V165_N1
        modelParams(77,1) = 0.001;      %koff_V165_N1H
        modelParams(78,1) = 0.001;      %koff_V165_R1
        modelParams(79,1) = 0.001;      %koff_V165_R2   
        modelParams(80,1) = 0.001;      %koff_V121_R1
        modelParams(81,1) = 0.001;      %koff_V121_R2
        modelParams(82,1) = 0.00049;    %koff_MMP9_LRP1   
        modelParams(83,1) = 0.001;      %koff_MMP3_proMMP9         
        
               
        modelParams(84,1) = 3.10E+13;   %kc_N_H
        modelParams(85,1) = 3.10E+13;   %kc_V165N_H
        modelParams(86,1) = 3.10E+13;   %kc_V165H_N
        modelParams(87,1) = 1.00E+14;   %kc_V165NH_R2
        modelParams(88,1) = 3.10E+13;   %kc_V165R2_NH
        modelParams(89,1) = 1.00E+14;   %kc_R1_N
        modelParams(90,1) = 3.10E+13;   %kc_R1_H
        modelParams(91,1) = 3.10E+11;   %kc_CD36_R2
        modelParams(92,1) = 3.10E+13;   %kc_CD36_B1
        modelParams(93,1) = 3.10E+11;   %kc_CD47_R2
        modelParams(94,1) = 3.96E+11;   %kc_HSGAG_FGFR1
        modelParams(95,1) = 3.17E+17;   %kc_FGFR1_FGFR1
        modelParams(96,1) = 0.001;      %kdissoc_N_H
        modelParams(97,1) = 0.0001;     %kdissoc_V165N_H
        modelParams(98,1) = 0.0001;     %kdissoc_V165H_N
        modelParams(99,1) = 0.001;      %kdissoc_R2_NH
        modelParams(100,1) = 0.01;      %kdissoc_R1_N
        modelParams(101,1) = 0.001/5;   %kdissoc_R1_H
        modelParams(102,1) = 0.001;     %kdissoc_CD36_B1
        modelParams(103,1) = 0.001;     %kdissoc_CD36_R2
        modelParams(104,1) = 0.001;     %kdissoc_CD47_R2
        modelParams(105,1) = 3.24E-07;  %kdissoc_HSGAG_FGFR1
        modelParams(106,1) = 5.83E-04;  %kdissoc_FGFR1_FGFR1 
       
        modelParams(107,1) = 0.000193;  %kdeg_VEGF
        modelParams(108,1) = 0.00033;   %kdeg_TSP1
        modelParams(109,1) = 0.000193;  %kdeg_FGF2
        modelParams(110,1) = 0.0023;    %kdeg_PF4
        modelParams(111,1) = 0.0012;    %kdeg_MMP
        modelParams(112,1) = 1.60E-4;   %kdeg_TSP1mim  
        
        modelParams(113,1) = 0.00386;   %k_TSP1cleave
        modelParams(114,1) = 0.0019;    %k_act_MMP3_proMMP9     
        modelParams(115,1) = 631;       %kp_mmp
        
        modelParams(116,1) = 2500;      %CD36_number_tum
        modelParams(117,1) = 10000;     %CD47_number_tum
        modelParams(118,1) = 5000;      %LRP1_number_tum
        modelParams(119,1) = 10000;     %B1_number_tum
       
        modelParams(120,1) = 1250;      %CD36_number_disEC
        modelParams(121,1) = 5000;      %CD47_number_disEC
        modelParams(122,1) = 2500;      %LRP1_number_disEC
        modelParams(123,1) = 5000;      %B1_number_disEC
       
        modelParams(124,1) = 1100;      %R1_number_tum
        modelParams(125,1) = 550;       %R2_number_tum
        modelParams(126,1) = 39500;     %N1_number_tum   
        modelParams(127,1) = 39500;     %N2_number_tum
      
        modelParams(128,1) = 3750;      %R1_number_disEC
        modelParams(129,1) = 300;       %R2_number_disEC
        modelParams(130,1) = 20000;     %N1_number_disEC      
        modelParams(131,1) = 20000;     %N2_number_disEC
        
        modelParams(132,1) = 20000;     %FGFR1_number_tum
        modelParams(133,1) = 100000;    %HSGAG_number_tum
       
        modelParams(134,1) = 10000;     %FGFR1_number_disEC     
        modelParams(135,1) = 50000;     %HSGAG_number_disEC
       
        modelParams(136,1) = 2500;      %CXCR3_number_tum
        modelParams(137,1) = 1250;      %CXCR3_number_disEC
      
end

parameters = modelParams;
% simulate to steady state
stopTime = 24*3600*3; % 1 day
[err, t, results, observables_out] = TissueModel(linspace(0,stopTime,1000)', [], parameters', []);

allObservables = [observables_out];

tsim = [linspace(0,stopTime,1000)];
tsim = tsim/3600;

%%%Paramters to convert mol/cm3 tissue --> pM (pmol/L)
ECM_Vol_tis_dis = 0.51931;
EBM_Vol_tis_dis = 0.00027;
PBM_Vol_tis_dis = 0.002446;
Interstit_Vol_tis_dis = ECM_Vol_tis_dis+EBM_Vol_tis_dis+PBM_Vol_tis_dis;


%%% VEGF distribution
VEGF_tot = allObservables(:,1)/Interstit_Vol_tis_dis*1e15;
VEGF_free = allObservables(:,2)/Interstit_Vol_tis_dis*1e15;
VEGF_TSP1 = allObservables(:,3)/Interstit_Vol_tis_dis*1e15;
VEGF_PF4 = allObservables(:,4)/Interstit_Vol_tis_dis*1e15;
VEGF_V114 = allObservables(:,5)/Interstit_Vol_tis_dis*1e15;
VEGF_R1 = allObservables(:,6)/Interstit_Vol_tis_dis*1e15;
VEGF_R2 = allObservables(:,7)/Interstit_Vol_tis_dis*1e15;
VEGF_N = allObservables(:,8)/Interstit_Vol_tis_dis*1e15;
VEGF_HSPG = allObservables(:,9)/Interstit_Vol_tis_dis*1e15;
VEGF_ECM = allObservables(:,10)/Interstit_Vol_tis_dis*1e15;

VEGF_percent = 100*[VEGF_free(end),VEGF_V114(end),VEGF_TSP1(end)+ VEGF_PF4(end),VEGF_ECM(end),VEGF_HSPG(end),VEGF_R1(end),VEGF_R2(end),VEGF_N(end)]./(VEGF_tot(end));

%%% FGF distribution
FGF_tot = allObservables(:,11)/Interstit_Vol_tis_dis*1e15;
FGF_free = allObservables(:,12)/Interstit_Vol_tis_dis*1e15;
FGF_TSP1 = allObservables(:,13)/Interstit_Vol_tis_dis*1e15;
FGF_PF4 = allObservables(:,14)/Interstit_Vol_tis_dis*1e15;
FGF_FGFR1 = allObservables(:,15)/Interstit_Vol_tis_dis*1e15;
FGF_HSPG = allObservables(:,16)/Interstit_Vol_tis_dis*1e15;
FGF_FGFR1_HSPG = allObservables(:,17)/Interstit_Vol_tis_dis*1e15;
FGF_dimer = allObservables(:,18)/Interstit_Vol_tis_dis*1e15;
FGF_ECM = allObservables(:,19)/Interstit_Vol_tis_dis*1e15;

FGF_percent = 100*[FGF_free(end), FGF_TSP1(end)+ FGF_PF4(end), FGF_ECM(end), FGF_HSPG(end), FGF_FGFR1(end),  FGF_FGFR1_HSPG(end), FGF_dimer(end)]./FGF_tot(end);

%%% TSP1 distribution
TSP1_tot = allObservables(:,20)/Interstit_Vol_tis_dis*1e15;
TSP1_cleaved = allObservables(:,21)/Interstit_Vol_tis_dis*1e15;
TSP1_free = allObservables(:,22)/Interstit_Vol_tis_dis*1e15;
TSP1_VEGF = allObservables(:,23)/Interstit_Vol_tis_dis*1e15;
TSP1_FGF = allObservables(:,24)/Interstit_Vol_tis_dis*1e15;
TSP1_MMP = allObservables(:,25)/Interstit_Vol_tis_dis*1e15;
TSP1_HSPG = allObservables(:,26)/Interstit_Vol_tis_dis*1e15;
TSP1_REC = allObservables(:,27)/Interstit_Vol_tis_dis*1e15;
TSP1_ECM = allObservables(:,28)/Interstit_Vol_tis_dis*1e15;

TSP1_percent = 100*[TSP1_free(end),TSP1_cleaved(end), TSP1_VEGF(end)+TSP1_FGF(end)+TSP1_MMP(end),  TSP1_ECM(end),TSP1_HSPG(end), TSP1_REC(end),]./TSP1_tot(end);

%%% PF4 distribution
PF4_tot = allObservables(:,29)/Interstit_Vol_tis_dis*1e15;
PF4_free = allObservables(:,30)/Interstit_Vol_tis_dis*1e15;
PF4_VEGF = allObservables(:,31)/Interstit_Vol_tis_dis*1e15;
PF4_FGF = allObservables(:,32)/Interstit_Vol_tis_dis*1e15;
PF4_HSPG = allObservables(:,33)/Interstit_Vol_tis_dis*1e15;
PF4_REC = allObservables(:,34)/Interstit_Vol_tis_dis*1e15;
PF4_ECM = allObservables(:,35)/Interstit_Vol_tis_dis*1e15;

PF4_percent = 100*[PF4_free(end), PF4_VEGF(end)+PF4_FGF(end), PF4_ECM(end),PF4_HSPG(end),  PF4_REC(end)]./PF4_tot(end);

MMP3_tot = allObservables(:,36)/Interstit_Vol_tis_dis*1e15;
MMP9_tot = allObservables(:,37)/Interstit_Vol_tis_dis*1e15;
MMP9 = allObservables(:,38)/Interstit_Vol_tis_dis*1e15;

Angiogenic_Ratio = (allObservables(end,39) + allObservables(end,40))/(allObservables(end,41) + allObservables(end,42));


disp('VEGF distribution in Percentage');
disp(VEGF_percent);
disp('FGF2 distribution in Percentage');
disp(FGF_percent);
disp('TSP1 distribution in Percentage');
disp(TSP1_percent);
disp('PF4 distribution in Percentage');
disp(PF4_percent);

disp('Angiogenic ratio');
disp(Angiogenic_Ratio);
```
