## Supplementary File S3 for "Exploring the extracellular regulation of the tumor angiogenic interaction network using a systems biology model": TissueModel.m

```
function [err, timepoints, species_out, observables_out] = TissueModel( timepoints, species_init, parameters, suppress_plot )
%TISSUEMODEL Integrate reaction network and plot observables.
%   Integrates the reaction network corresponding to the BioNetGen model
%   'TissueModel_HSPG' and then (optionally) plots the observable trajectories,
%   or species trajectories if no observables are defined. Trajectories are
%   generated using either default or user-defined parameters and initial
%   species values. Integration is performed by the MATLAB stiff solver
%   'ode15s'. TISSUEMODEL_HSPG returns an error value, a vector of timepoints,
%   species trajectories, and observable trajectories.
%   
%   [err, timepoints, species_out, observables_out]
%        = TissueModel_HSPG( timepoints, species_init, parameters, suppress_plot )
%
%   INPUTS:
%   -------
%   species_init    : row vector of 168 initial species populations.
%   timepoints      : column vector of time points returned by integrator.
%   parameters      : row vector of 137 model parameters.
%   suppress_plot   : 0 if a plot is desired (default), 1 if plot is suppressed.
%
%   Note: to specify default value for an input argument, pass the empty array.
%
%   OUTPUTS:
%   --------
%   err             : 0 if the integrator exits without error, non-zero otherwise.
%   timepoints      : a row vector of timepoints returned by the integrator.
%   species_out     : array of species population trajectories
%                        (columns correspond to species, rows correspond to time).
%   observables_out : array of observable trajectories
%                        (columns correspond to observables, rows correspond to time).
%
%   QUESTIONS about the BNG Mfile generator? 


%% Process input arguments

% define any missing arguments
if ( nargin < 1 )
    timepoints = [];
end

if ( nargin < 2 )
    species_init = [];
end

if ( nargin < 3 )
    parameters = [];
end

if ( nargin < 4 )
    suppress_plot = 0;
end


% initialize outputs (to avoid error msgs if script terminates early
err = 0;
species_out     = [];
observables_out = [];


% setup default parameters, if necessary
if ( isempty(parameters) )
   parameters = [ 70, 6.02E+23, 7.5e-7, 1.3e-5, 1.3e-5, 0.51931, 0.00027, 0.002446, 33.51032, 1534, 105, 9.97E-06, 1.00E-05, 6, 0.6, 180, 330, 0, 0, 0.387, 0.387, 0.0324, 0.0036, 3e-16, 1e-14, 0.00028, 0.00028, 6.26E+06, 8600, 5.71E+05, 1.89E+04, 2.00E+04, 8600, 5.4054e+05, 4.20e+03, 2E+05, 31250, 5E+03, 8600, 5.00E+05, 5.00E+05, 2.10E+05, 5.00E+05, 5.00E+05, 1.00E+05, 8600, 8600, 3.20E+06, 3.20E+06, 3.00E+07, 1.00E+07, 3.00E+07, 1.00E+07, 9245, 10000, 1.95E-03, 0.000336375, 2.75E-04, 0.000205, 0.001, 0.0001725, 0.001, 0.001, 0.001, 0.001, 0.001, 0.000353625, 0.115, 0.005, 0.0025, 0.05, 0.005, 0.0022303, 0.00069, 0.00069, 0.020, 0.001, 0.001, 0.001, 0.001, 0.001, 0.00049, 0.001, 3.10E+13, 3.10E+13, 3.10E+13, 1.00E+14, 3.10E+13, 1.00E+14, 3.10E+13, 3.10E+11, 3.10E+13, 3.10E+11, 1.87E+13, 2.33E+13, 0.001, 0.0001, 0.0001, 0.001, 0.01, 0.0002, 0.001, 0.001, 0.001, 2.14E-01, 1.35E-02, 0.000193, 0.00033, 0.0000253, 0.0023, 0.0012, 1.60E-4, 0.00386, 0.0019, 631, 2500, 10000, 5000, 10000, 1250, 5000, 2500, 5000, 1100, 550, 39500, 39500, 3750, 300, 20000, 20000, 20000, 100000, 10000, 50000, 2500, 1250 ];
end
% check that parameters has proper dimensions
if (  size(parameters,1) ~= 1  ||  size(parameters,2) ~= 137  )
    fprintf( 1, 'Error: size of parameter argument is invalid! Correct size = [1 137].\n' );
    err = 1;
    return;
end

% setup default initial values, if necessary
if ( isempty(species_init) )
   species_init = initialize_species( parameters );
end
% check that species_init has proper dimensions
if (  size(species_init,1) ~= 1  ||  size(species_init,2) ~= 168  )
    fprintf( 1, 'Error: size of species_init argument is invalid! Correct size = [1 168].\n' );
    err = 1;
    return;
end

% setup default timepoints, if necessary
if ( isempty(timepoints) )
   timepoints = linspace(0,10*3600,600)';
end
% check that timepoints has proper dimensions
if (  size(timepoints,1) < 2  ||  size(timepoints,2) ~= 1  )
    fprintf( 1, 'Error: size of timepoints argument is invalid! Correct size = [t 1], t>1.\n' );
    err = 1;
    return;
end

% setup default suppress_plot, if necessary
if ( isempty(suppress_plot) )
   suppress_plot = 0;
end
% check that suppress_plot has proper dimensions
if ( size(suppress_plot,1) ~= 1  ||  size(suppress_plot,2) ~= 1 )
    fprintf( 1, 'Error: suppress_plots argument should be a scalar!\n' );
    err = 1;
    return;
end

% define parameter labels (this is for the user's reference!)
param_labels = { 'BW', 'Avogadro', 'ECM_conc', 'EBM_conc', 'PBM_conc', 'ECM_Vol_tis_dis', 'EBM_Vol_tis_dis', 'PBM_Vol_tis_dis', 'vol_Tumor', 'tumorSA_Vol_tis_dis', 'VesselSA_Vol_tis_dis', 'tumorCellSurfArea_tis_dis', 'VesselCellSurfArea_tis_dis', 'qTSP1disEC', 'qTSP1tum', 'qMMP3_disEC', 'qproMMP9_disEC', 'qMMP3tum', 'qproMMP9tum', 'qV165_tumor', 'qV121_tumor', 'qV165_disEC', 'qV121_disEC', 'qFGF2', 'qPF4', 'sR_receptors', 'k_int_receptors', 'kon_FGF2_HSPG', 'kon_FGF2_GAG', 'kon_FGF2_FGFR1', 'kon_FGF2_TSP1', 'kon_PF4_HSPG', 'kon_PF4_GAG', 'kon_PF4_CXCR3', 'kon_PF4_LRP1', 'kon_PF4_V165', 'kon_PF4_FGF2', 'kon_TSP1_HSPG', 'kon_TSP1_GAG', 'kon_TSP1_CD36', 'kon_TSP1_CD47', 'kon_TSP1_LRP1', 'kon_TSP1_B1', 'kon_TSP1_VEGF', 'kon_TSP1_MMP3', 'kon_V165_HSPG', 'kon_V165_GAG', 'kon_V165_N1', 'kon_V165_N1H', 'kon_V165_R1', 'kon_V165_R2', 'kon_V121_R1', 'kon_V121_R2', 'kon_MMP9_LRP1', 'kon_MMP3_proMMP9', 'koff_FGF2_HSPG', 'koff_FGF2_GAG', 'koff_FGF2_FGFR1', 'koff_FGF2_TSP1', 'koff_PF4_HSPG', 'koff_PF4_GAG', 'koff_PF4_CXCR3', 'koff_PF4_LRP1', 'koff_PF4_V165', 'koff_PF4_FGF2', 'koff_TSP1_HSPG', 'koff_TSP1_GAG', 'koff_TSP1_CD36', 'koff_TSP1_CD47', 'koff_TSP1_LRP1', 'koff_TSP1_B1', 'koff_TSP1_VEGF', 'koff_TSP1_MMP3', 'koff_V165_HSPG', 'koff_V165_GAG', 'koff_V165_N1', 'koff_V165_N1H', 'koff_V165_R1', 'koff_V165_R2', 'koff_V121_R1', 'koff_V121_R2', 'koff_MMP9_LRP1', 'koff_MMP3_proMMP9', 'kc_N_H', 'kc_V165N_H', 'kc_V165H_N', 'kc_V165NH_R2', 'kc_V165R2_NH', 'kc_R1_N', 'kc_R1_H', 'kc_CD36_R2', 'kc_CD36_B1', 'kc_CD47_R2', 'kc_HSPG_FGFR1', 'kc_FGFR1_FGFR1', 'kdissoc_N_H', 'kdissoc_V165N_H', 'kdissoc_V165H_N', 'kdissoc_R2_NH', 'kdissoc_R1_N', 'kdissoc_R1_H', 'kdissoc_CD36_B1', 'kdissoc_CD36_R2', 'kdissoc_CD47_R2', 'kdissoc_HSPG_FGFR1', 'kdissoc_FGFR1_FGFR1', 'kdeg_VEGF', 'kdeg_TSP1', 'kdeg_FGF2', 'kdeg_PF4', 'kdeg_MMP', 'kdeg_TSP1mim', 'k_TSP1cleave', 'k_act_MMP3_proMMP9', 'kp_mmp', 'CD36_number_tum', 'CD47_number_tum', 'LRP1_number_tum', 'B1_number_tum', 'CD36_number_disEC', 'CD47_number_disEC', 'LRP1_number_disEC', 'B1_number_disEC', 'R1_number_tum', 'R2_number_tum', 'N1_number_tum', 'N2_number_tum', 'R1_number_disEC', 'R2_number_disEC', 'N1_number_disEC', 'N2_number_disEC', 'FGFR1_number_tum', 'HSPG_number_tum', 'FGFR1_number_disEC', 'HSPG_number_disEC', 'CXCR3_number_tum', 'CXCR3_number_disEC' };


%% Integrate Network Model
 
% calculate expressions
[expressions] = calc_expressions( parameters );

% set ODE integrator options
opts = odeset( 'RelTol',   1e-12,   ...
               'AbsTol',   1e-18,   ...
               'Stats',    'off',  ...
               'BDF',      'off',    ...
               'MaxOrder', 5   );


% define derivative function
rhs_fcn = @(t,y)( calc_species_deriv( t, y, expressions ) );

% simulate model system (stiff integrator)
try 
    [~, species_out] = ode15s( rhs_fcn, timepoints, species_init', opts );
    if(length(timepoints) ~= size(species_out,1))
        exception = MException('ODE15sError:MissingOutput','Not all timepoints output\n');
        throw(exception);
    end
catch
    err = 1;
    fprintf( 1, 'Error: some problem encountered while integrating ODE network!\n' );
    return;
end

% calculate observables
observables_out = zeros( length(timepoints), 64 );
for t = 1 : length(timepoints)
    observables_out(t,:) = calc_observables( species_out(t,:), expressions );
end


%% Plot Output, if desired

if ( ~suppress_plot )
    
    % define plot labels
    observable_labels = { 'VEGF_tot', 'VEGF_free', 'VEGF_TSP1', 'VEGF_PF4', 'VEGF_free114', 'VEGF_R1', 'VEGF_R2', 'VEGF_N', 'VEGF_HSPG', 'VEGF_ECM', 'FGF_tot', 'FGF_free', 'FGF_TSP1', 'FGF_PF4', 'FGF_FGFR1', 'FGF_HSPG', 'FGF_FGFR1_HSPG', 'FGF_dimer', 'FGF_ECM', 'TSP1_tot', 'TSP1_cleaved', 'TSP1_free', 'TSP1_VEGF', 'TSP1_FGF', 'TSP1_MMP', 'TSP1_HSPG', 'TSP1_REC', 'TSP1_ECM', 'PF4_tot', 'PF4_free', 'PF4_VEGF', 'PF4_FGF', 'PF4_HSPG', 'PF4_REC', 'PF4_ECM', 'MMP3_tot', 'MMP9_tot', 'MMP9_active', 'VEGF_pro', 'FGF_pro', 'TSP1_anti', 'PF4_anti', 'VEGF_165', 'VEGF_121', 'V121_tot', 'V121_R2', 'V121_R1', 'V121_TSP1', 'V165_tot', 'V165_free', 'V165_HSPG', 'V165_GAG', 'V165_R1', 'V165_R2', 'V165_N', 'V121tot', 'V121_free', 'V121R1', 'V121R2', 'TSP1_V165', 'TSP1_V121', 'TSP1_V114', 'R1free', 'R1N' };

    % construct figure
    plot(timepoints,observables_out);
    title('TissueModel_HSPG observables','fontSize',14,'Interpreter','none');
    axis([0 timepoints(end) 0 inf]);
    legend(observable_labels,'fontSize',10,'Interpreter','none');
    xlabel('time','fontSize',12,'Interpreter','none');
    ylabel('number or concentration','fontSize',12,'Interpreter','none');

end


%~~~~~~~~~~~~~~~~~~~~~%
% END of main script! %
%~~~~~~~~~~~~~~~~~~~~~%


% initialize species function
function [species_init] = initialize_species( params )

    species_init = zeros(1,168);
    species_init(1) = 0;
    species_init(2) = 1;
    species_init(3) = ((params(116)/params(12))/params(2))*params(10);
    species_init(4) = ((params(119)/params(12))/params(2))*params(10);
    species_init(5) = ((params(117)/params(12))/params(2))*params(10);
    species_init(6) = ((params(118)/params(12))/params(2))*params(10);
    species_init(7) = ((params(120)/params(13))/params(2))*params(11);
    species_init(8) = ((params(123)/params(13))/params(2))*params(11);
    species_init(9) = ((params(121)/params(13))/params(2))*params(11);
    species_init(10) = ((params(122)/params(13))/params(2))*params(11);
    species_init(11) = ((params(124)/params(12))/params(2))*params(10);
    species_init(12) = ((params(125)/params(12))/params(2))*params(10);
    species_init(13) = ((params(126)/params(12))/params(2))*params(10);
    species_init(14) = ((params(127)/params(12))/params(2))*params(10);
    species_init(15) = ((params(128)/params(13))/params(2))*params(11);
    species_init(16) = ((params(129)/params(13))/params(2))*params(11);
    species_init(17) = ((params(130)/params(13))/params(2))*params(11);
    species_init(18) = ((params(131)/params(13))/params(2))*params(11);
    species_init(19) = ((params(133)/params(12))/params(2))*params(10);
    species_init(20) = ((params(135)/params(13))/params(2))*params(11);
    species_init(21) = 0;
    species_init(22) = ((params(132)/params(12))/params(2))*params(10);
    species_init(23) = ((params(134)/params(13))/params(2))*params(11);
    species_init(24) = 0;
    species_init(25) = 0;
    species_init(26) = ((params(136)/params(12))/params(2))*params(10);
    species_init(27) = ((params(137)/params(13))/params(2))*params(11);
    species_init(28) = (params(3)*params(6))/1000;
    species_init(29) = (params(4)*params(7))/1000;
    species_init(30) = (params(5)*params(8))/1000;
    species_init(31) = 0;
    species_init(32) = 0;
    species_init(33) = 0;
    species_init(34) = 0;
    species_init(35) = 0;
    species_init(36) = 0;
    species_init(37) = 0;
    species_init(38) = 0;
    species_init(39) = 0;
    species_init(40) = 0;
    species_init(41) = 0;
    species_init(42) = 0;
    species_init(43) = 0;
    species_init(44) = 0;
    species_init(45) = 0;
    species_init(46) = 0;
    species_init(47) = 0;
    species_init(48) = 0;
    species_init(49) = 0;
    species_init(50) = 0;
    species_init(51) = 0;
    species_init(52) = 0;
    species_init(53) = 0;
    species_init(54) = 0;
    species_init(55) = 0;
    species_init(56) = 0;
    species_init(57) = 0;
    species_init(58) = 0;
    species_init(59) = 0;
    species_init(60) = 0;
    species_init(61) = 0;
    species_init(62) = 0;
    species_init(63) = 0;
    species_init(64) = 0;
    species_init(65) = 0;
    species_init(66) = 0;
    species_init(67) = 0;
    species_init(68) = 0;
    species_init(69) = 0;
    species_init(70) = 0;
    species_init(71) = 0;
    species_init(72) = 0;
    species_init(73) = 0;
    species_init(74) = 0;
    species_init(75) = 0;
    species_init(76) = 0;
    species_init(77) = 0;
    species_init(78) = 0;
    species_init(79) = 0;
    species_init(80) = 0;
    species_init(81) = 0;
    species_init(82) = 0;
    species_init(83) = 0;
    species_init(84) = 0;
    species_init(85) = 0;
    species_init(86) = 0;
    species_init(87) = 0;
    species_init(88) = 0;
    species_init(89) = 0;
    species_init(90) = 0;
    species_init(91) = 0;
    species_init(92) = 0;
    species_init(93) = 0;
    species_init(94) = 0;
    species_init(95) = 0;
    species_init(96) = 0;
    species_init(97) = 0;
    species_init(98) = 0;
    species_init(99) = 0;
    species_init(100) = 0;
    species_init(101) = 0;
    species_init(102) = 0;
    species_init(103) = 0;
    species_init(104) = 0;
    species_init(105) = 0;
    species_init(106) = 0;
    species_init(107) = 0;
    species_init(108) = 0;
    species_init(109) = 0;
    species_init(110) = 0;
    species_init(111) = 0;
    species_init(112) = 0;
    species_init(113) = 0;
    species_init(114) = 0;
    species_init(115) = 0;
    species_init(116) = 0;
    species_init(117) = 0;
    species_init(118) = 0;
    species_init(119) = 0;
    species_init(120) = 0;
    species_init(121) = 0;
    species_init(122) = 0;
    species_init(123) = 0;
    species_init(124) = 0;
    species_init(125) = 0;
    species_init(126) = 0;
    species_init(127) = 0;
    species_init(128) = 0;
    species_init(129) = 0;
    species_init(130) = 0;
    species_init(131) = 0;
    species_init(132) = 0;
    species_init(133) = 0;
    species_init(134) = 0;
    species_init(135) = 0;
    species_init(136) = 0;
    species_init(137) = 0;
    species_init(138) = 0;
    species_init(139) = 0;
    species_init(140) = 0;
    species_init(141) = 0;
    species_init(142) = 0;
    species_init(143) = 0;
    species_init(144) = 0;
    species_init(145) = 0;
    species_init(146) = 0;
    species_init(147) = 0;
    species_init(148) = 0;
    species_init(149) = 0;
    species_init(150) = 0;
    species_init(151) = 0;
    species_init(152) = 0;
    species_init(153) = 0;
    species_init(154) = 0;
    species_init(155) = 0;
    species_init(156) = 0;
    species_init(157) = 0;
    species_init(158) = 0;
    species_init(159) = 0;
    species_init(160) = 0;
    species_init(161) = 0;
    species_init(162) = 0;
    species_init(163) = 0;
    species_init(164) = 0;
    species_init(165) = 0;
    species_init(166) = 0;
    species_init(167) = 0;
    species_init(168) = 0;

end


% user-defined functions


% Calculate expressions
function [ expressions ] = calc_expressions ( parameters )

    expressions = zeros(1,253);
    expressions(1) = parameters(1);
    expressions(2) = parameters(2);
    expressions(3) = parameters(3);
    expressions(4) = parameters(4);
    expressions(5) = parameters(5);
    expressions(6) = parameters(6);
    expressions(7) = parameters(7);
    expressions(8) = parameters(8);
    expressions(9) = parameters(9);
    expressions(10) = parameters(10);
    expressions(11) = parameters(11);
    expressions(12) = parameters(12);
    expressions(13) = parameters(13);
    expressions(14) = parameters(14);
    expressions(15) = parameters(15);
    expressions(16) = parameters(16);
    expressions(17) = parameters(17);
    expressions(18) = parameters(18);
    expressions(19) = parameters(19);
    expressions(20) = parameters(20);
    expressions(21) = parameters(21);
    expressions(22) = parameters(22);
    expressions(23) = parameters(23);
    expressions(24) = parameters(24);
    expressions(25) = parameters(25);
    expressions(26) = parameters(26);
    expressions(27) = parameters(27);
    expressions(28) = parameters(28);
    expressions(29) = parameters(29);
    expressions(30) = parameters(30);
    expressions(31) = parameters(31);
    expressions(32) = parameters(32);
    expressions(33) = parameters(33);
    expressions(34) = parameters(34);
    expressions(35) = parameters(35);
    expressions(36) = parameters(36);
    expressions(37) = parameters(37);
    expressions(38) = parameters(38);
    expressions(39) = parameters(39);
    expressions(40) = parameters(40);
    expressions(41) = parameters(41);
    expressions(42) = parameters(42);
    expressions(43) = parameters(43);
    expressions(44) = parameters(44);
    expressions(45) = parameters(45);
    expressions(46) = parameters(46);
    expressions(47) = parameters(47);
    expressions(48) = parameters(48);
    expressions(49) = parameters(49);
    expressions(50) = parameters(50);
    expressions(51) = parameters(51);
    expressions(52) = parameters(52);
    expressions(53) = parameters(53);
    expressions(54) = parameters(54);
    expressions(55) = parameters(55);
    expressions(56) = parameters(56);
    expressions(57) = parameters(57);
    expressions(58) = parameters(58);
    expressions(59) = parameters(59);
    expressions(60) = parameters(60);
    expressions(61) = parameters(61);
    expressions(62) = parameters(62);
    expressions(63) = parameters(63);
    expressions(64) = parameters(64);
    expressions(65) = parameters(65);
    expressions(66) = parameters(66);
    expressions(67) = parameters(67);
    expressions(68) = parameters(68);
    expressions(69) = parameters(69);
    expressions(70) = parameters(70);
    expressions(71) = parameters(71);
    expressions(72) = parameters(72);
    expressions(73) = parameters(73);
    expressions(74) = parameters(74);
    expressions(75) = parameters(75);
    expressions(76) = parameters(76);
    expressions(77) = parameters(77);
    expressions(78) = parameters(78);
    expressions(79) = parameters(79);
    expressions(80) = parameters(80);
    expressions(81) = parameters(81);
    expressions(82) = parameters(82);
    expressions(83) = parameters(83);
    expressions(84) = parameters(84);
    expressions(85) = parameters(85);
    expressions(86) = parameters(86);
    expressions(87) = parameters(87);
    expressions(88) = parameters(88);
    expressions(89) = parameters(89);
    expressions(90) = parameters(90);
    expressions(91) = parameters(91);
    expressions(92) = parameters(92);
    expressions(93) = parameters(93);
    expressions(94) = parameters(94);
    expressions(95) = parameters(95);
    expressions(96) = parameters(96);
    expressions(97) = parameters(97);
    expressions(98) = parameters(98);
    expressions(99) = parameters(99);
    expressions(100) = parameters(100);
    expressions(101) = parameters(101);
    expressions(102) = parameters(102);
    expressions(103) = parameters(103);
    expressions(104) = parameters(104);
    expressions(105) = parameters(105);
    expressions(106) = parameters(106);
    expressions(107) = parameters(107);
    expressions(108) = parameters(108);
    expressions(109) = parameters(109);
    expressions(110) = parameters(110);
    expressions(111) = parameters(111);
    expressions(112) = parameters(112);
    expressions(113) = parameters(113);
    expressions(114) = parameters(114);
    expressions(115) = parameters(115);
    expressions(116) = parameters(116);
    expressions(117) = parameters(117);
    expressions(118) = parameters(118);
    expressions(119) = parameters(119);
    expressions(120) = parameters(120);
    expressions(121) = parameters(121);
    expressions(122) = parameters(122);
    expressions(123) = parameters(123);
    expressions(124) = parameters(124);
    expressions(125) = parameters(125);
    expressions(126) = parameters(126);
    expressions(127) = parameters(127);
    expressions(128) = parameters(128);
    expressions(129) = parameters(129);
    expressions(130) = parameters(130);
    expressions(131) = parameters(131);
    expressions(132) = parameters(132);
    expressions(133) = parameters(133);
    expressions(134) = parameters(134);
    expressions(135) = parameters(135);
    expressions(136) = parameters(136);
    expressions(137) = parameters(137);
    expressions(138) = ((expressions(6)+expressions(7))+expressions(8));
    expressions(139) = ((((expressions(26)*expressions(116))/expressions(12))/expressions(2))*expressions(10));
    expressions(140) = ((((expressions(26)*expressions(119))/expressions(12))/expressions(2))*expressions(10));
    expressions(141) = ((((expressions(26)*expressions(117))/expressions(12))/expressions(2))*expressions(10));
    expressions(142) = ((((expressions(26)*expressions(118))/expressions(12))/expressions(2))*expressions(10));
    expressions(143) = ((((expressions(26)*expressions(124))/expressions(12))/expressions(2))*expressions(10));
    expressions(144) = ((((expressions(26)*expressions(125))/expressions(12))/expressions(2))*expressions(10));
    expressions(145) = ((((expressions(26)*expressions(126))/expressions(12))/expressions(2))*expressions(10));
    expressions(146) = ((((expressions(26)*expressions(127))/expressions(12))/expressions(2))*expressions(10));
    expressions(147) = ((((expressions(26)*expressions(132))/expressions(12))/expressions(2))*expressions(10));
    expressions(148) = ((((expressions(26)*expressions(133))/expressions(12))/expressions(2))*expressions(10));
    expressions(149) = ((((expressions(26)*expressions(136))/expressions(12))/expressions(2))*expressions(10));
    expressions(150) = ((((expressions(26)*expressions(120))/expressions(13))/expressions(2))*expressions(11));
    expressions(151) = ((((expressions(26)*expressions(123))/expressions(13))/expressions(2))*expressions(11));
    expressions(152) = ((((expressions(26)*expressions(121))/expressions(13))/expressions(2))*expressions(11));
    expressions(153) = ((((expressions(26)*expressions(122))/expressions(13))/expressions(2))*expressions(11));
    expressions(154) = ((((expressions(26)*expressions(128))/expressions(13))/expressions(2))*expressions(11));
    expressions(155) = ((((expressions(26)*expressions(129))/expressions(13))/expressions(2))*expressions(11));
    expressions(156) = ((((expressions(26)*expressions(130))/expressions(13))/expressions(2))*expressions(11));
    expressions(157) = ((((expressions(26)*expressions(131))/expressions(13))/expressions(2))*expressions(11));
    expressions(158) = ((((expressions(26)*expressions(134))/expressions(13))/expressions(2))*expressions(11));
    expressions(159) = ((((expressions(26)*expressions(135))/expressions(13))/expressions(2))*expressions(11));
    expressions(160) = ((((expressions(26)*expressions(137))/expressions(13))/expressions(2))*expressions(11));
    expressions(161) = ((expressions(38)*1000)/expressions(138));
    expressions(162) = ((expressions(39)*1000)/expressions(138));
    expressions(163) = ((expressions(40)*1000)/expressions(138));
    expressions(164) = ((expressions(41)*1000)/expressions(138));
    expressions(165) = ((expressions(42)*1000)/expressions(138));
    expressions(166) = ((expressions(43)*1000)/expressions(138));
    expressions(167) = ((expressions(44)*1000)/expressions(138));
    expressions(168) = ((expressions(45)*1000)/expressions(138));
    expressions(169) = ((expressions(48)*1000)/expressions(138));
    expressions(170) = ((expressions(49)*1000)/expressions(138));
    expressions(171) = ((expressions(50)*1000)/expressions(138));
    expressions(172) = ((expressions(51)*1000)/expressions(138));
    expressions(173) = ((expressions(46)*1000)/expressions(138));
    expressions(174) = ((expressions(47)*1000)/expressions(138));
    expressions(175) = ((expressions(52)*1000)/expressions(138));
    expressions(176) = ((expressions(53)*1000)/expressions(138));
    expressions(177) = ((expressions(54)*1000)/expressions(138));
    expressions(178) = ((expressions(55)*1000)/expressions(138));
    expressions(179) = ((expressions(28)*1000)/expressions(138));
    expressions(180) = ((expressions(30)*1000)/expressions(138));
    expressions(181) = ((expressions(31)*1000)/expressions(138));
    expressions(182) = ((expressions(29)*1000)/expressions(138));
    expressions(183) = ((expressions(32)*1000)/expressions(138));
    expressions(184) = ((expressions(33)*1000)/expressions(138));
    expressions(185) = ((expressions(34)*1000)/expressions(138));
    expressions(186) = ((expressions(35)*1000)/expressions(138));
    expressions(187) = ((expressions(36)*1000)/expressions(138));
    expressions(188) = ((expressions(37)*1000)/expressions(138));
    expressions(189) = (expressions(84)/expressions(10));
    expressions(190) = (expressions(85)/expressions(10));
    expressions(191) = (expressions(86)/expressions(10));
    expressions(192) = (expressions(87)/expressions(10));
    expressions(193) = (expressions(88)/expressions(10));
    expressions(194) = (expressions(89)/expressions(10));
    expressions(195) = (expressions(90)/expressions(10));
    expressions(196) = (expressions(91)/expressions(10));
    expressions(197) = (expressions(92)/expressions(10));
    expressions(198) = (expressions(93)/expressions(10));
    expressions(199) = (expressions(94)/expressions(10));
    expressions(200) = (expressions(95)/expressions(10));
    expressions(201) = (expressions(84)/expressions(11));
    expressions(202) = (expressions(85)/expressions(11));
    expressions(203) = (expressions(86)/expressions(11));
    expressions(204) = (expressions(87)/expressions(11));
    expressions(205) = (expressions(88)/expressions(11));
    expressions(206) = (expressions(89)/expressions(11));
    expressions(207) = (expressions(90)/expressions(11));
    expressions(208) = (expressions(91)/expressions(11));
    expressions(209) = (expressions(92)/expressions(11));
    expressions(210) = (expressions(93)/expressions(11));
    expressions(211) = (expressions(94)/expressions(11));
    expressions(212) = (expressions(95)/expressions(11));
    expressions(213) = (((expressions(15)/expressions(2))/expressions(12))*expressions(10));
    expressions(214) = (((expressions(14)/expressions(2))/expressions(13))*expressions(11));
    expressions(215) = (((expressions(18)/expressions(2))/expressions(12))*expressions(10));
    expressions(216) = (((expressions(19)/expressions(2))/expressions(12))*expressions(10));
    expressions(217) = (((expressions(16)/expressions(2))/expressions(13))*expressions(11));
    expressions(218) = (((expressions(17)/expressions(2))/expressions(13))*expressions(11));
    expressions(219) = (((expressions(21)/expressions(2))/expressions(12))*expressions(10));
    expressions(220) = (((expressions(20)/expressions(2))/expressions(12))*expressions(10));
    expressions(221) = (((expressions(23)/expressions(2))/expressions(13))*expressions(11));
    expressions(222) = (((expressions(22)/expressions(2))/expressions(13))*expressions(11));
    expressions(223) = ((expressions(115)*1000)/expressions(138));
    expressions(224) = ((expressions(3)*expressions(6))/1000);
    expressions(225) = ((expressions(4)*expressions(7))/1000);
    expressions(226) = ((expressions(5)*expressions(8))/1000);
    expressions(227) = (((expressions(124)/expressions(12))/expressions(2))*expressions(10));
    expressions(228) = (((expressions(125)/expressions(12))/expressions(2))*expressions(10));
    expressions(229) = (((expressions(126)/expressions(12))/expressions(2))*expressions(10));
    expressions(230) = (((expressions(127)/expressions(12))/expressions(2))*expressions(10));
    expressions(231) = (((expressions(116)/expressions(12))/expressions(2))*expressions(10));
    expressions(232) = (((expressions(119)/expressions(12))/expressions(2))*expressions(10));
    expressions(233) = (((expressions(117)/expressions(12))/expressions(2))*expressions(10));
    expressions(234) = (((expressions(118)/expressions(12))/expressions(2))*expressions(10));
    expressions(235) = (((expressions(132)/expressions(12))/expressions(2))*expressions(10));
    expressions(236) = (((expressions(133)/expressions(12))/expressions(2))*expressions(10));
    expressions(237) = (((expressions(136)/expressions(12))/expressions(2))*expressions(10));
    expressions(238) = (((expressions(128)/expressions(13))/expressions(2))*expressions(11));
    expressions(239) = (((expressions(129)/expressions(13))/expressions(2))*expressions(11));
    expressions(240) = (((expressions(130)/expressions(13))/expressions(2))*expressions(11));
    expressions(241) = (((expressions(131)/expressions(13))/expressions(2))*expressions(11));
    expressions(242) = (((expressions(120)/expressions(13))/expressions(2))*expressions(11));
    expressions(243) = (((expressions(123)/expressions(13))/expressions(2))*expressions(11));
    expressions(244) = (((expressions(121)/expressions(13))/expressions(2))*expressions(11));
    expressions(245) = (((expressions(122)/expressions(13))/expressions(2))*expressions(11));
    expressions(246) = (((expressions(134)/expressions(13))/expressions(2))*expressions(11));
    expressions(247) = (((expressions(135)/expressions(13))/expressions(2))*expressions(11));
    expressions(248) = (((expressions(137)/expressions(13))/expressions(2))*expressions(11));
    expressions(249) = (expressions(213)+expressions(214));
    expressions(250) = (expressions(220)+expressions(222));
    expressions(251) = (expressions(219)+expressions(221));
    expressions(252) = (expressions(218)+expressions(216));
    expressions(253) = (expressions(217)+expressions(215));
   
end


% Calculate observables
function [ observables ] = calc_observables ( species, expressions )

    observables = zeros(1,64);
    observables(1) = species(32) +species(33) +species(34) +species(43) +species(44) +species(45) +species(46) +species(49) +species(50) +species(51) +species(52) +species(53) +species(54) +species(55) +species(56) +species(59) +species(60) +species(61) +species(85) +species(86) +species(87) +species(109) +species(111) +species(112) +species(114) +species(115) +species(117) +species(118) +species(119) +species(120) +species(121) +species(122) +species(123) +species(124) +species(125) +species(126) +species(127) +species(128) +species(129) +species(130) +species(141) +species(142) +species(143) +species(144) +species(145) +species(146) +species(147) +species(148) +species(149) +species(150) +species(151) +species(152) +species(153) +species(154) +species(155) +species(156) +species(157) +species(158) +species(159) +species(160) +species(161) +species(162) +species(165) +species(166) +species(167) +species(168);
    observables(2) = species(32) +species(33);
    observables(3) = species(59) +species(60) +species(149) +species(150) +species(151) +species(152);
    observables(4) = species(109);
    observables(5) = species(34) +species(61) +species(153) +species(154);
    observables(6) = species(43) +species(44) +species(45) +species(46) +species(111) +species(112) +species(114) +species(115) +species(117) +species(118) +species(155) +species(156);
    observables(7) = species(49) +species(50) +species(51) +species(52) +species(119) +species(120) +species(121) +species(122) +species(123) +species(124) +species(125) +species(126) +species(129) +species(130) +species(141) +species(142) +species(143) +species(144) +species(145) +species(146) +species(147) +species(148) +species(157) +species(158) +species(159) +species(160) +species(161) +species(162) +species(165) +species(166) +species(167) +species(168);
    observables(8) = species(55) +species(56) +species(127) +species(128);
    observables(9) = species(53) +species(54);
    observables(10) = species(85) +species(86) +species(87);
    observables(11) = 2*species(24) +2*species(25) +species(39) +species(88) +species(89) +species(90) +species(94) +species(95) +species(96) +species(97) +species(98) +species(99) +species(100) +species(101) +species(110);
    observables(12) = species(39);
    observables(13) = species(99);
    observables(14) = species(110);
    observables(15) = species(97) +species(98);
    observables(16) = species(94) +species(95);
    observables(17) = species(100) +species(101);
    observables(18) = 2*species(24) +2*species(25);
    observables(19) = species(88) +species(89) +species(90);
    observables(20) = species(31) +species(38) +species(59) +species(60) +species(61) +species(62) +species(63) +species(66) +species(67) +species(68) +species(69) +species(74) +species(75) +species(76) +species(77) +species(79) +species(82) +species(83) +species(84) +species(99) +species(131) +species(132) +species(133) +species(134) +species(135) +species(136) +species(138) +species(140) +species(141) +species(143) +species(145) +species(147) +species(149) +species(150) +species(151) +species(152) +species(153) +species(154) +species(157) +species(158) +species(159) +species(160) +species(163) +species(164) +species(165) +species(166) +species(167) +species(168);
    observables(21) = species(38);
    observables(22) = species(31);
    observables(23) = species(59) +species(60) +species(61);
    observables(24) = species(99);
    observables(25) = species(79);
    observables(26) = species(76) +species(77);
    observables(27) = species(62) +species(63) +species(66) +species(67) +species(68) +species(69) +species(74) +species(75) +species(131) +species(132) +species(133) +species(134) +species(135) +species(136) +species(138) +species(140) +species(141) +species(143) +species(145) +species(147) +species(149) +species(150) +species(151) +species(152) +species(153) +species(154) +species(157) +species(158) +species(159) +species(160) +species(163) +species(164) +species(165) +species(166) +species(167) +species(168);
    observables(28) = species(82) +species(83) +species(84);
    observables(29) = species(40) +species(91) +species(92) +species(93) +species(102) +species(103) +species(104) +species(105) +species(106) +species(107) +species(108) +species(109) +species(110);
    observables(30) = species(40);
    observables(31) = species(109);
    observables(32) = species(110);
    observables(33) = species(102) +species(103);
    observables(34) = species(105) +species(106) +species(107) +species(108);
    observables(35) = species(91) +species(92) +species(93);
    observables(36) = species(37) +species(78) +species(79);
    observables(37) = species(35) +species(36) +species(78) +species(80) +species(81);
    observables(38) = species(36) +species(80) +species(81);
    observables(39) = species(43) +species(44) +species(45) +species(46) +species(49) +species(50) +species(51) +species(52) +species(55) +species(56) +species(111) +species(112) +species(114) +species(115) +species(117) +species(118) +species(119) +species(120) +species(121) +species(122) +species(123) +species(124) +species(125) +species(126) +species(127) +species(128) +species(129) +species(130) +species(142) +species(144) +species(146) +species(148) +species(155) +species(156) +species(161) +species(162);
    observables(40) = species(24) +species(25);
    observables(41) = species(62) +species(63) +species(66) +species(67) +species(68) +species(69) +species(74) +species(75) +species(131) +species(132) +species(133) +species(134) +species(135) +species(136) +species(138) +species(140) +species(141) +species(143) +species(145) +species(147) +species(149) +species(150) +species(151) +species(152) +species(153) +species(154) +species(157) +species(158) +species(159) +species(160) +species(163) +species(164) +species(165) +species(166) +species(167) +species(168);
    observables(42) = species(105) +species(106) +species(107) +species(108);
    observables(43) = species(32);
    observables(44) = species(33);
    observables(45) = species(33) +species(45) +species(46) +species(49) +species(50) +species(60) +species(112) +species(115) +species(117) +species(118) +species(119) +species(120) +species(121) +species(122) +species(141) +species(142) +species(145) +species(146) +species(151) +species(152) +species(155) +species(156) +species(157) +species(158) +species(165) +species(167);
    observables(46) = species(49) +species(50) +species(119) +species(120) +species(121) +species(122) +species(141) +species(142) +species(145) +species(146) +species(157) +species(158) +species(165) +species(167);
    observables(47) = species(45) +species(46) +species(112) +species(115) +species(117) +species(118) +species(155) +species(156);
    observables(48) = species(60) +species(151) +species(152);
    observables(49) = species(32) +species(43) +species(44) +species(51) +species(52) +species(53) +species(54) +species(55) +species(56) +species(59) +species(85) +species(86) +species(87) +species(109) +species(111) +species(114) +species(123) +species(124) +species(125) +species(126) +species(127) +species(128) +species(129) +species(130) +species(143) +species(144) +species(147) +species(148) +species(149) +species(150) +species(159) +species(160) +species(161) +species(162) +species(166) +species(168);
    observables(50) = species(32);
    observables(51) = species(53) +species(54);
    observables(52) = species(85) +species(86) +species(87);
    observables(53) = species(43) +species(44) +species(111) +species(114);
    observables(54) = species(51) +species(52) +species(123) +species(124) +species(125) +species(126) +species(129) +species(130) +species(143) +species(144) +species(147) +species(148) +species(159) +species(160) +species(161) +species(162) +species(166) +species(168);
    observables(55) = species(55) +species(56) +species(127) +species(128);
    observables(56) = species(33) +species(45) +species(46) +species(49) +species(50) +species(60) +species(112) +species(115) +species(117) +species(118) +species(119) +species(120) +species(121) +species(122) +species(141) +species(142) +species(145) +species(146) +species(151) +species(152) +species(155) +species(156) +species(157) +species(158) +species(165) +species(167);
    observables(57) = species(33);
    observables(58) = species(45) +species(46) +species(112) +species(115) +species(117) +species(118) +species(155) +species(156);
    observables(59) = species(49) +species(50) +species(119) +species(120) +species(121) +species(122) +species(141) +species(142) +species(145) +species(146) +species(157) +species(158) +species(165) +species(167);
    observables(60) = species(59) +species(149) +species(150);
    observables(61) = species(60) +species(151) +species(152);
    observables(62) = species(61) +species(153) +species(154);
    observables(63) = species(11) +species(15) +species(41) +species(42);
    observables(64) = species(47) +species(48) +species(113) +species(116);

end


% Calculate ratelaws
function [ ratelaws ] = calc_ratelaws ( species, expressions, observables )

    ratelaws = zeros(1,64);
    ratelaws(1) = ((((expressions(26)*expressions(116))/expressions(12))/expressions(2))*expressions(10))*species(2);
    ratelaws(2) = ((((expressions(26)*expressions(119))/expressions(12))/expressions(2))*expressions(10))*species(2);
    ratelaws(3) = ((((expressions(26)*expressions(117))/expressions(12))/expressions(2))*expressions(10))*species(2);
    ratelaws(4) = ((((expressions(26)*expressions(118))/expressions(12))/expressions(2))*expressions(10))*species(2);
    ratelaws(5) = ((((expressions(26)*expressions(120))/expressions(13))/expressions(2))*expressions(11))*species(2);
    ratelaws(6) = ((((expressions(26)*expressions(123))/expressions(13))/expressions(2))*expressions(11))*species(2);
    ratelaws(7) = ((((expressions(26)*expressions(121))/expressions(13))/expressions(2))*expressions(11))*species(2);
    ratelaws(8) = ((((expressions(26)*expressions(122))/expressions(13))/expressions(2))*expressions(11))*species(2);
    ratelaws(9) = ((((expressions(26)*expressions(124))/expressions(12))/expressions(2))*expressions(10))*species(2);
    ratelaws(10) = ((((expressions(26)*expressions(125))/expressions(12))/expressions(2))*expressions(10))*species(2);
    ratelaws(11) = ((((expressions(26)*expressions(126))/expressions(12))/expressions(2))*expressions(10))*species(2);
    ratelaws(12) = ((((expressions(26)*expressions(127))/expressions(12))/expressions(2))*expressions(10))*species(2);
    ratelaws(13) = ((((expressions(26)*expressions(128))/expressions(13))/expressions(2))*expressions(11))*species(2);
    ratelaws(14) = ((((expressions(26)*expressions(129))/expressions(13))/expressions(2))*expressions(11))*species(2);
    ratelaws(15) = ((((expressions(26)*expressions(130))/expressions(13))/expressions(2))*expressions(11))*species(2);
    ratelaws(16) = ((((expressions(26)*expressions(131))/expressions(13))/expressions(2))*expressions(11))*species(2);
    ratelaws(17) = ((((expressions(26)*expressions(133))/expressions(12))/expressions(2))*expressions(10))*species(2);
    ratelaws(18) = ((((expressions(26)*expressions(135))/expressions(13))/expressions(2))*expressions(11))*species(2);
    ratelaws(19) = ((((expressions(26)*expressions(132))/expressions(12))/expressions(2))*expressions(10))*species(2);
    ratelaws(20) = ((((expressions(26)*expressions(134))/expressions(13))/expressions(2))*expressions(11))*species(2);
    ratelaws(21) = ((((expressions(26)*expressions(136))/expressions(12))/expressions(2))*expressions(10))*species(2);
    ratelaws(22) = ((((expressions(26)*expressions(137))/expressions(13))/expressions(2))*expressions(11))*species(2);
    ratelaws(23) = expressions(27)*species(19);
    ratelaws(24) = expressions(27)*species(20);
    ratelaws(25) = expressions(27)*species(4);
    ratelaws(26) = expressions(27)*species(8);
    ratelaws(27) = expressions(27)*species(3);
    ratelaws(28) = expressions(27)*species(7);
    ratelaws(29) = expressions(27)*species(5);
    ratelaws(30) = expressions(27)*species(9);
    ratelaws(31) = expressions(27)*species(6);
    ratelaws(32) = expressions(27)*species(10);
    ratelaws(33) = expressions(27)*species(11);
    ratelaws(34) = expressions(27)*species(15);
    ratelaws(35) = expressions(27)*species(12);
    ratelaws(36) = expressions(27)*species(16);
    ratelaws(37) = expressions(27)*species(14);
    ratelaws(38) = expressions(27)*species(18);
    ratelaws(39) = expressions(27)*species(13);
    ratelaws(40) = expressions(27)*species(17);
    ratelaws(41) = expressions(27)*species(22);
    ratelaws(42) = expressions(27)*species(23);
    ratelaws(43) = expressions(27)*species(24);
    ratelaws(44) = expressions(27)*species(25);
    ratelaws(45) = expressions(27)*species(26);
    ratelaws(46) = expressions(27)*species(27);
    ratelaws(47) = (expressions(213)+expressions(214))*species(2);
    ratelaws(48) = expressions(113)*species(31);
    ratelaws(49) = expressions(108)*species(31);
    ratelaws(50) = expressions(108)*species(38);
    ratelaws(51) = expressions(24)*species(2);
    ratelaws(52) = expressions(109)*species(39);
    ratelaws(53) = expressions(25)*species(2);
    ratelaws(54) = expressions(110)*species(40);
    ratelaws(55) = (expressions(220)+expressions(222))*species(2);
    ratelaws(56) = (expressions(219)+expressions(221))*species(2);
    ratelaws(57) = expressions(107)*species(32);
    ratelaws(58) = expressions(107)*species(33);
    ratelaws(59) = expressions(107)*species(34);
    ratelaws(60) = (expressions(218)+expressions(216))*species(2);
    ratelaws(61) = (expressions(217)+expressions(215))*species(2);
    ratelaws(62) = expressions(111)*species(37);
    ratelaws(63) = expressions(111)*species(36);
    ratelaws(64) = expressions(111)*species(35);
    ratelaws(65) = (expressions(90)/expressions(10))*species(11)*species(19);
    ratelaws(66) = (expressions(90)/expressions(11))*species(15)*species(20);
    ratelaws(67) = ((expressions(50)*1000)/expressions(138))*species(32)*species(11);
    ratelaws(68) = ((expressions(50)*1000)/expressions(138))*species(32)*species(15);
    ratelaws(69) = ((expressions(52)*1000)/expressions(138))*species(33)*species(11);
    ratelaws(70) = ((expressions(52)*1000)/expressions(138))*species(33)*species(15);
    ratelaws(71) = (expressions(89)/expressions(10))*species(11)*species(13);
    ratelaws(72) = (expressions(89)/expressions(11))*species(15)*species(17);
    ratelaws(73) = ((expressions(53)*1000)/expressions(138))*species(33)*species(12);
    ratelaws(74) = ((expressions(53)*1000)/expressions(138))*species(33)*species(16);
    ratelaws(75) = ((expressions(51)*1000)/expressions(138))*species(32)*species(12);
    ratelaws(76) = ((expressions(51)*1000)/expressions(138))*species(32)*species(16);
    ratelaws(77) = ((expressions(46)*1000)/expressions(138))*species(32)*species(19);
    ratelaws(78) = ((expressions(46)*1000)/expressions(138))*species(32)*species(20);
    ratelaws(79) = ((expressions(48)*1000)/expressions(138))*species(32)*species(13);
    ratelaws(80) = ((expressions(48)*1000)/expressions(138))*species(32)*species(17);
    ratelaws(81) = (expressions(84)/expressions(10))*species(19)*species(13);
    ratelaws(82) = (expressions(84)/expressions(11))*species(20)*species(17);
    ratelaws(83) = ((expressions(44)*1000)/expressions(138))*species(31)*species(32);
    ratelaws(84) = ((expressions(44)*1000)/expressions(138))*species(31)*species(33);
    ratelaws(85) = ((expressions(44)*1000)/expressions(138))*species(31)*species(34);
    ratelaws(86) = ((expressions(41)*1000)/expressions(138))*species(31)*species(5);
    ratelaws(87) = ((expressions(41)*1000)/expressions(138))*species(31)*species(9);
    ratelaws(88) = (expressions(93)/expressions(10))*species(5)*species(12);
    ratelaws(89) = (expressions(93)/expressions(11))*species(9)*species(16);
    ratelaws(90) = ((expressions(40)*1000)/expressions(138))*species(31)*species(3);
    ratelaws(91) = ((expressions(40)*1000)/expressions(138))*species(31)*species(7);
    ratelaws(92) = ((expressions(43)*1000)/expressions(138))*species(31)*species(4);
    ratelaws(93) = ((expressions(43)*1000)/expressions(138))*species(31)*species(8);
    ratelaws(94) = (expressions(92)/expressions(10))*species(4)*species(3);
    ratelaws(95) = (expressions(92)/expressions(11))*species(8)*species(7);
    ratelaws(96) = (expressions(91)/expressions(10))*species(12)*species(3);
    ratelaws(97) = (expressions(91)/expressions(11))*species(16)*species(7);
    ratelaws(98) = ((expressions(42)*1000)/expressions(138))*species(31)*species(6);
    ratelaws(99) = ((expressions(42)*1000)/expressions(138))*species(31)*species(10);
    ratelaws(100) = ((expressions(38)*1000)/expressions(138))*species(31)*species(19);
    ratelaws(101) = ((expressions(38)*1000)/expressions(138))*species(31)*species(20);
    ratelaws(102) = ((expressions(55)*1000)/expressions(138))*species(35)*species(37);
    ratelaws(103) = ((expressions(45)*1000)/expressions(138))*species(37)*species(31);
    ratelaws(104) = ((expressions(115)*1000)/expressions(138))*species(37)*species(32);
    ratelaws(105) = ((expressions(115)*1000)/expressions(138))*species(36)*species(32);
    ratelaws(106) = ((expressions(54)*1000)/expressions(138))*species(36)*species(6);
    ratelaws(107) = ((expressions(54)*1000)/expressions(138))*species(36)*species(10);
    ratelaws(108) = ((expressions(39)*1000)/expressions(138))*species(31)*species(28);
    ratelaws(109) = ((expressions(39)*1000)/expressions(138))*species(31)*species(29);
    ratelaws(110) = ((expressions(39)*1000)/expressions(138))*species(31)*species(30);
    ratelaws(111) = ((expressions(47)*1000)/expressions(138))*species(32)*species(28);
    ratelaws(112) = ((expressions(47)*1000)/expressions(138))*species(32)*species(29);
    ratelaws(113) = ((expressions(47)*1000)/expressions(138))*species(32)*species(30);
    ratelaws(114) = ((expressions(29)*1000)/expressions(138))*species(39)*species(28);
    ratelaws(115) = ((expressions(29)*1000)/expressions(138))*species(39)*species(29);
    ratelaws(116) = ((expressions(29)*1000)/expressions(138))*species(39)*species(30);
    ratelaws(117) = ((expressions(33)*1000)/expressions(138))*species(40)*species(28);
    ratelaws(118) = ((expressions(33)*1000)/expressions(138))*species(40)*species(29);
    ratelaws(119) = ((expressions(33)*1000)/expressions(138))*species(40)*species(30);
    ratelaws(120) = ((expressions(28)*1000)/expressions(138))*species(39)*species(19);
    ratelaws(121) = ((expressions(28)*1000)/expressions(138))*species(39)*species(20);
    ratelaws(122) = ((expressions(28)*1000)/expressions(138))*species(39)*species(21);
    ratelaws(123) = ((expressions(30)*1000)/expressions(138))*species(39)*species(22);
    ratelaws(124) = ((expressions(30)*1000)/expressions(138))*species(39)*species(23);
    ratelaws(125) = ((expressions(31)*1000)/expressions(138))*species(39)*species(31);
    ratelaws(126) = expressions(106)*species(24);
    ratelaws(127) = expressions(106)*species(25);
    ratelaws(128) = ((expressions(32)*1000)/expressions(138))*species(40)*species(19);
    ratelaws(129) = ((expressions(32)*1000)/expressions(138))*species(40)*species(20);
    ratelaws(130) = ((expressions(32)*1000)/expressions(138))*species(40)*species(21);
    ratelaws(131) = ((expressions(34)*1000)/expressions(138))*species(40)*species(26);
    ratelaws(132) = ((expressions(34)*1000)/expressions(138))*species(40)*species(27);
    ratelaws(133) = ((expressions(35)*1000)/expressions(138))*species(40)*species(6);
    ratelaws(134) = ((expressions(35)*1000)/expressions(138))*species(40)*species(10);
    ratelaws(135) = ((expressions(36)*1000)/expressions(138))*species(40)*species(32);
    ratelaws(136) = ((expressions(37)*1000)/expressions(138))*species(40)*species(39);
    ratelaws(137) = expressions(27)*species(41);
    ratelaws(138) = expressions(27)*species(53);
    ratelaws(139) = expressions(27)*species(57);
    ratelaws(140) = expressions(27)*species(76);
    ratelaws(141) = expressions(27)*species(94);
    ratelaws(142) = expressions(27)*species(100);
    ratelaws(143) = expressions(27)*species(102);
    ratelaws(144) = expressions(27)*species(42);
    ratelaws(145) = expressions(27)*species(54);
    ratelaws(146) = expressions(27)*species(58);
    ratelaws(147) = expressions(27)*species(77);
    ratelaws(148) = expressions(27)*species(95);
    ratelaws(149) = expressions(27)*species(101);
    ratelaws(150) = expressions(27)*species(103);
    ratelaws(151) = expressions(27)*species(68);
    ratelaws(152) = expressions(27)*species(69);
    ratelaws(153) = expressions(27)*species(70);
    ratelaws(154) = expressions(27)*species(71);
    ratelaws(155) = expressions(27)*species(66);
    ratelaws(156) = expressions(27)*species(67);
    ratelaws(157) = expressions(27)*species(72);
    ratelaws(158) = expressions(27)*species(73);
    ratelaws(159) = expressions(27)*species(62);
    ratelaws(160) = expressions(27)*species(63);
    ratelaws(161) = expressions(27)*species(64);
    ratelaws(162) = expressions(27)*species(65);
    ratelaws(163) = expressions(27)*species(74);
    ratelaws(164) = expressions(27)*species(75);
    ratelaws(165) = expressions(27)*species(80);
    ratelaws(166) = expressions(27)*species(81);
    ratelaws(167) = expressions(27)*species(107);
    ratelaws(168) = expressions(27)*species(108);
    ratelaws(169) = expressions(27)*species(43);
    ratelaws(170) = expressions(27)*species(44);
    ratelaws(171) = expressions(27)*species(45);
    ratelaws(172) = expressions(27)*species(46);
    ratelaws(173) = expressions(27)*species(47);
    ratelaws(174) = expressions(27)*species(48);
    ratelaws(175) = expressions(27)*species(49);
    ratelaws(176) = expressions(27)*species(50);
    ratelaws(177) = expressions(27)*species(51);
    ratelaws(178) = expressions(27)*species(52);
    ratelaws(179) = expressions(27)*species(55);
    ratelaws(180) = expressions(27)*species(56);
    ratelaws(181) = expressions(27)*species(97);
    ratelaws(182) = expressions(27)*species(98);
    ratelaws(183) = expressions(27)*species(105);
    ratelaws(184) = expressions(27)*species(106);
    ratelaws(185) = expressions(111)*species(78);
    ratelaws(186) = (expressions(90)/expressions(10))*species(43)*species(19);
    ratelaws(187) = (expressions(90)/expressions(10))*species(45)*species(19);
    ratelaws(188) = (expressions(90)/expressions(10))*species(47)*species(19);
    ratelaws(189) = expressions(101)*species(41);
    ratelaws(190) = (expressions(90)/expressions(11))*species(44)*species(20);
    ratelaws(191) = (expressions(90)/expressions(11))*species(46)*species(20);
    ratelaws(192) = (expressions(90)/expressions(11))*species(48)*species(20);
    ratelaws(193) = expressions(101)*species(42);
    ratelaws(194) = ((expressions(50)*1000)/expressions(138))*species(32)*species(41);
    ratelaws(195) = ((expressions(50)*1000)/expressions(138))*species(32)*species(42);
    ratelaws(196) = expressions(78)*species(43);
    ratelaws(197) = expressions(78)*species(44);
    ratelaws(198) = ((expressions(52)*1000)/expressions(138))*species(33)*species(41);
    ratelaws(199) = ((expressions(52)*1000)/expressions(138))*species(33)*species(42);
    ratelaws(200) = ((expressions(52)*1000)/expressions(138))*species(33)*species(47);
    ratelaws(201) = ((expressions(52)*1000)/expressions(138))*species(33)*species(48);
    ratelaws(202) = expressions(80)*species(45);
    ratelaws(203) = expressions(80)*species(46);
    ratelaws(204) = (expressions(89)/expressions(10))*species(41)*species(13);
    ratelaws(205) = expressions(100)*species(47);
    ratelaws(206) = (expressions(89)/expressions(11))*species(42)*species(17);
    ratelaws(207) = expressions(100)*species(48);
    ratelaws(208) = (expressions(89)/expressions(10))*species(45)*species(13);
    ratelaws(209) = (expressions(89)/expressions(11))*species(46)*species(17);
    ratelaws(210) = ((expressions(53)*1000)/expressions(138))*species(33)*species(64);
    ratelaws(211) = ((expressions(53)*1000)/expressions(138))*species(33)*species(65);
    ratelaws(212) = ((expressions(53)*1000)/expressions(138))*species(33)*species(72);
    ratelaws(213) = ((expressions(53)*1000)/expressions(138))*species(33)*species(73);
    ratelaws(214) = expressions(81)*species(49);
    ratelaws(215) = expressions(81)*species(50);
    ratelaws(216) = ((expressions(51)*1000)/expressions(138))*species(32)*species(64);
    ratelaws(217) = ((expressions(51)*1000)/expressions(138))*species(32)*species(65);
    ratelaws(218) = ((expressions(51)*1000)/expressions(138))*species(32)*species(72);
    ratelaws(219) = ((expressions(51)*1000)/expressions(138))*species(32)*species(73);
    ratelaws(220) = expressions(79)*species(51);
    ratelaws(221) = expressions(79)*species(52);
    ratelaws(222) = expressions(74)*species(53);
    ratelaws(223) = expressions(74)*species(54);
    ratelaws(224) = expressions(76)*species(55);
    ratelaws(225) = expressions(76)*species(56);
    ratelaws(226) = ((expressions(49)*1000)/expressions(138))*species(32)*species(57);
    ratelaws(227) = ((expressions(49)*1000)/expressions(138))*species(32)*species(58);
    ratelaws(228) = expressions(96)*species(57);
    ratelaws(229) = expressions(96)*species(58);
    ratelaws(230) = (expressions(86)/expressions(10))*species(53)*species(13);
    ratelaws(231) = (expressions(86)/expressions(11))*species(54)*species(17);
    ratelaws(232) = (expressions(85)/expressions(10))*species(55)*species(19);
    ratelaws(233) = (expressions(85)/expressions(11))*species(56)*species(20);
    ratelaws(234) = (expressions(88)/expressions(10))*species(51)*species(57);
    ratelaws(235) = (expressions(88)/expressions(11))*species(52)*species(58);
    ratelaws(236) = expressions(72)*species(59);
    ratelaws(237) = expressions(72)*species(60);
    ratelaws(238) = expressions(72)*species(61);
    ratelaws(239) = expressions(107)*species(59);
    ratelaws(240) = expressions(107)*species(60);
    ratelaws(241) = expressions(107)*species(61);
    ratelaws(242) = expressions(69)*species(62);
    ratelaws(243) = expressions(69)*species(63);
    ratelaws(244) = expressions(104)*species(64);
    ratelaws(245) = expressions(104)*species(65);
    ratelaws(246) = ((expressions(40)*1000)/expressions(138))*species(31)*species(72);
    ratelaws(247) = ((expressions(40)*1000)/expressions(138))*species(31)*species(73);
    ratelaws(248) = expressions(68)*species(66);
    ratelaws(249) = expressions(68)*species(67);
    ratelaws(250) = expressions(71)*species(68);
    ratelaws(251) = expressions(71)*species(69);
    ratelaws(252) = ((expressions(40)*1000)/expressions(138))*species(31)*species(70);
    ratelaws(253) = ((expressions(40)*1000)/expressions(138))*species(31)*species(71);
    ratelaws(254) = ((expressions(43)*1000)/expressions(138))*species(31)*species(70);
    ratelaws(255) = ((expressions(43)*1000)/expressions(138))*species(31)*species(71);
    ratelaws(256) = (expressions(92)/expressions(10))*species(4)*species(66);
    ratelaws(257) = (expressions(92)/expressions(10))*species(4)*species(72);
    ratelaws(258) = expressions(102)*species(70);
    ratelaws(259) = (expressions(92)/expressions(10))*species(68)*species(3);
    ratelaws(260) = (expressions(92)/expressions(10))*species(68)*species(72);
    ratelaws(261) = (expressions(92)/expressions(11))*species(8)*species(67);
    ratelaws(262) = (expressions(92)/expressions(11))*species(8)*species(73);
    ratelaws(263) = expressions(102)*species(71);
    ratelaws(264) = (expressions(92)/expressions(11))*species(69)*species(7);
    ratelaws(265) = (expressions(92)/expressions(11))*species(69)*species(73);
    ratelaws(266) = (expressions(91)/expressions(10))*species(12)*species(66);
    ratelaws(267) = (expressions(91)/expressions(10))*species(12)*species(70);
    ratelaws(268) = (expressions(91)/expressions(10))*species(49)*species(3);
    ratelaws(269) = (expressions(91)/expressions(10))*species(49)*species(66);
    ratelaws(270) = (expressions(91)/expressions(10))*species(49)*species(70);
    ratelaws(271) = (expressions(91)/expressions(10))*species(51)*species(3);
    ratelaws(272) = (expressions(91)/expressions(10))*species(51)*species(66);
    ratelaws(273) = (expressions(91)/expressions(10))*species(51)*species(70);
    ratelaws(274) = expressions(103)*species(72);
    ratelaws(275) = (expressions(91)/expressions(11))*species(16)*species(67);
    ratelaws(276) = (expressions(91)/expressions(11))*species(16)*species(71);
    ratelaws(277) = (expressions(91)/expressions(11))*species(50)*species(7);
    ratelaws(278) = (expressions(91)/expressions(11))*species(50)*species(67);
    ratelaws(279) = (expressions(91)/expressions(11))*species(50)*species(71);
    ratelaws(280) = (expressions(91)/expressions(11))*species(52)*species(7);
    ratelaws(281) = (expressions(91)/expressions(11))*species(52)*species(67);
    ratelaws(282) = (expressions(91)/expressions(11))*species(52)*species(71);
    ratelaws(283) = expressions(103)*species(73);
    ratelaws(284) = ((expressions(42)*1000)/expressions(138))*species(59)*species(6);
    ratelaws(285) = ((expressions(42)*1000)/expressions(138))*species(59)*species(10);
    ratelaws(286) = ((expressions(42)*1000)/expressions(138))*species(60)*species(6);
    ratelaws(287) = ((expressions(42)*1000)/expressions(138))*species(60)*species(10);
    ratelaws(288) = ((expressions(42)*1000)/expressions(138))*species(61)*species(6);
    ratelaws(289) = ((expressions(42)*1000)/expressions(138))*species(61)*species(10);
    ratelaws(290) = expressions(70)*species(74);
    ratelaws(291) = expressions(70)*species(75);
    ratelaws(292) = ((expressions(44)*1000)/expressions(138))*species(32)*species(74);
    ratelaws(293) = ((expressions(44)*1000)/expressions(138))*species(32)*species(75);
    ratelaws(294) = ((expressions(44)*1000)/expressions(138))*species(33)*species(74);
    ratelaws(295) = ((expressions(44)*1000)/expressions(138))*species(33)*species(75);
    ratelaws(296) = expressions(66)*species(76);
    ratelaws(297) = expressions(66)*species(77);
    ratelaws(298) = expressions(83)*species(78);
    ratelaws(299) = expressions(114)*species(78);
    ratelaws(300) = expressions(73)*species(79);
    ratelaws(301) = expressions(82)*species(80);
    ratelaws(302) = expressions(82)*species(81);
    ratelaws(303) = expressions(108)*species(79);
    ratelaws(304) = expressions(67)*species(82);
    ratelaws(305) = expressions(67)*species(83);
    ratelaws(306) = expressions(67)*species(84);
    ratelaws(307) = expressions(75)*species(85);
    ratelaws(308) = expressions(75)*species(86);
    ratelaws(309) = expressions(75)*species(87);
    ratelaws(310) = expressions(57)*species(88);
    ratelaws(311) = expressions(57)*species(89);
    ratelaws(312) = expressions(57)*species(90);
    ratelaws(313) = expressions(61)*species(91);
    ratelaws(314) = expressions(61)*species(92);
    ratelaws(315) = expressions(61)*species(93);
    ratelaws(316) = ((expressions(115)*1000)/expressions(138))*species(85)*species(37);
    ratelaws(317) = ((expressions(115)*1000)/expressions(138))*species(85)*species(36);
    ratelaws(318) = expressions(56)*species(94);
    ratelaws(319) = expressions(56)*species(95);
    ratelaws(320) = expressions(56)*species(96);
    ratelaws(321) = expressions(58)*species(97);
    ratelaws(322) = expressions(58)*species(98);
    ratelaws(323) = expressions(59)*species(99);
    ratelaws(324) = expressions(108)*species(99);
    ratelaws(325) = (expressions(94)/expressions(10))*species(94)*species(22);
    ratelaws(326) = expressions(105)*species(100);
    ratelaws(327) = (expressions(94)/expressions(11))*species(95)*species(23);
    ratelaws(328) = expressions(105)*species(101);
    ratelaws(329) = 0.5*(expressions(95)/expressions(10))*species(100)*species(100);
    ratelaws(330) = 0.5*(expressions(95)/expressions(11))*species(101)*species(101);
    ratelaws(331) = expressions(60)*species(102);
    ratelaws(332) = expressions(60)*species(103);
    ratelaws(333) = expressions(60)*species(104);
    ratelaws(334) = expressions(62)*species(105);
    ratelaws(335) = expressions(62)*species(106);
    ratelaws(336) = expressions(63)*species(107);
    ratelaws(337) = expressions(63)*species(108);
    ratelaws(338) = expressions(64)*species(109);
    ratelaws(339) = expressions(65)*species(110);
    ratelaws(340) = expressions(107)*species(109);
    ratelaws(341) = expressions(109)*species(110);
    ratelaws(342) = expressions(27)*species(111);
    ratelaws(343) = expressions(27)*species(112);
    ratelaws(344) = expressions(27)*species(113);
    ratelaws(345) = expressions(27)*species(127);
    ratelaws(346) = 2*expressions(27)*species(129);
    ratelaws(347) = expressions(27)*species(114);
    ratelaws(348) = expressions(27)*species(115);
    ratelaws(349) = expressions(27)*species(116);
    ratelaws(350) = expressions(27)*species(128);
    ratelaws(351) = 2*expressions(27)*species(130);
    ratelaws(352) = expressions(27)*species(133);
    ratelaws(353) = expressions(27)*species(134);
    ratelaws(354) = expressions(27)*species(135);
    ratelaws(355) = expressions(27)*species(136);
    ratelaws(356) = expressions(27)*species(137);
    ratelaws(357) = expressions(27)*species(138);
    ratelaws(358) = expressions(27)*species(139);
    ratelaws(359) = expressions(27)*species(140);
    ratelaws(360) = expressions(27)*species(142);
    ratelaws(361) = expressions(27)*species(144);
    ratelaws(362) = expressions(27)*species(146);
    ratelaws(363) = expressions(27)*species(148);
    ratelaws(364) = expressions(27)*species(121);
    ratelaws(365) = expressions(27)*species(122);
    ratelaws(366) = expressions(27)*species(125);
    ratelaws(367) = expressions(27)*species(126);
    ratelaws(368) = expressions(27)*species(131);
    ratelaws(369) = expressions(27)*species(132);
    ratelaws(370) = expressions(27)*species(141);
    ratelaws(371) = expressions(27)*species(143);
    ratelaws(372) = expressions(27)*species(145);
    ratelaws(373) = expressions(27)*species(147);
    ratelaws(374) = expressions(27)*species(119);
    ratelaws(375) = expressions(27)*species(120);
    ratelaws(376) = expressions(27)*species(123);
    ratelaws(377) = expressions(27)*species(124);
    ratelaws(378) = expressions(27)*species(149);
    ratelaws(379) = expressions(27)*species(150);
    ratelaws(380) = expressions(27)*species(151);
    ratelaws(381) = expressions(27)*species(152);
    ratelaws(382) = expressions(27)*species(153);
    ratelaws(383) = expressions(27)*species(154);
    ratelaws(384) = expressions(27)*species(117);
    ratelaws(385) = expressions(27)*species(118);
    ratelaws(386) = (expressions(90)/expressions(10))*species(117)*species(19);
    ratelaws(387) = expressions(101)*species(111);
    ratelaws(388) = expressions(101)*species(112);
    ratelaws(389) = expressions(101)*species(113);
    ratelaws(390) = (expressions(90)/expressions(11))*species(118)*species(20);
    ratelaws(391) = expressions(101)*species(114);
    ratelaws(392) = expressions(101)*species(115);
    ratelaws(393) = expressions(101)*species(116);
    ratelaws(394) = expressions(78)*species(111);
    ratelaws(395) = expressions(78)*species(114);
    ratelaws(396) = ((expressions(52)*1000)/expressions(138))*species(33)*species(113);
    ratelaws(397) = ((expressions(52)*1000)/expressions(138))*species(33)*species(116);
    ratelaws(398) = expressions(80)*species(112);
    ratelaws(399) = expressions(80)*species(115);
    ratelaws(400) = expressions(80)*species(117);
    ratelaws(401) = expressions(80)*species(118);
    ratelaws(402) = expressions(100)*species(113);
    ratelaws(403) = expressions(100)*species(116);
    ratelaws(404) = (expressions(89)/expressions(10))*species(112)*species(13);
    ratelaws(405) = expressions(100)*species(117);
    ratelaws(406) = (expressions(89)/expressions(11))*species(115)*species(17);
    ratelaws(407) = expressions(100)*species(118);
    ratelaws(408) = ((expressions(53)*1000)/expressions(138))*species(33)*species(131);
    ratelaws(409) = ((expressions(53)*1000)/expressions(138))*species(33)*species(132);
    ratelaws(410) = ((expressions(53)*1000)/expressions(138))*species(33)*species(137);
    ratelaws(411) = ((expressions(53)*1000)/expressions(138))*species(33)*species(138);
    ratelaws(412) = ((expressions(53)*1000)/expressions(138))*species(33)*species(139);
    ratelaws(413) = ((expressions(53)*1000)/expressions(138))*species(33)*species(140);
    ratelaws(414) = expressions(81)*species(119);
    ratelaws(415) = expressions(81)*species(120);
    ratelaws(416) = expressions(81)*species(121);
    ratelaws(417) = expressions(81)*species(122);
    ratelaws(418) = expressions(81)*species(141);
    ratelaws(419) = expressions(81)*species(142);
    ratelaws(420) = expressions(81)*species(145);
    ratelaws(421) = expressions(81)*species(146);
    ratelaws(422) = ((expressions(51)*1000)/expressions(138))*species(32)*species(131);
    ratelaws(423) = ((expressions(51)*1000)/expressions(138))*species(32)*species(132);
    ratelaws(424) = ((expressions(51)*1000)/expressions(138))*species(32)*species(137);
    ratelaws(425) = ((expressions(51)*1000)/expressions(138))*species(32)*species(138);
    ratelaws(426) = ((expressions(51)*1000)/expressions(138))*species(32)*species(139);
    ratelaws(427) = ((expressions(51)*1000)/expressions(138))*species(32)*species(140);
    ratelaws(428) = expressions(79)*species(123);
    ratelaws(429) = expressions(79)*species(124);
    ratelaws(430) = expressions(79)*species(125);
    ratelaws(431) = expressions(79)*species(126);
    ratelaws(432) = expressions(79)*species(143);
    ratelaws(433) = expressions(79)*species(144);
    ratelaws(434) = expressions(79)*species(147);
    ratelaws(435) = expressions(79)*species(148);
    ratelaws(436) = expressions(77)*species(127);
    ratelaws(437) = expressions(77)*species(128);
    ratelaws(438) = expressions(98)*species(127);
    ratelaws(439) = expressions(98)*species(128);
    ratelaws(440) = expressions(97)*species(127);
    ratelaws(441) = expressions(97)*species(128);
    ratelaws(442) = expressions(99)*species(129);
    ratelaws(443) = expressions(99)*species(130);
    ratelaws(444) = (expressions(87)/expressions(10))*species(12)*species(127);
    ratelaws(445) = (expressions(87)/expressions(11))*species(16)*species(128);
    ratelaws(446) = ((expressions(40)*1000)/expressions(138))*species(31)*species(121);
    ratelaws(447) = ((expressions(40)*1000)/expressions(138))*species(31)*species(122);
    ratelaws(448) = ((expressions(40)*1000)/expressions(138))*species(31)*species(125);
    ratelaws(449) = ((expressions(40)*1000)/expressions(138))*species(31)*species(126);
    ratelaws(450) = expressions(68)*species(131);
    ratelaws(451) = expressions(68)*species(132);
    ratelaws(452) = expressions(68)*species(141);
    ratelaws(453) = expressions(68)*species(143);
    ratelaws(454) = expressions(68)*species(145);
    ratelaws(455) = expressions(68)*species(147);
    ratelaws(456) = ((expressions(40)*1000)/expressions(138))*species(31)*species(137);
    ratelaws(457) = ((expressions(40)*1000)/expressions(138))*species(31)*species(139);
    ratelaws(458) = ((expressions(40)*1000)/expressions(138))*species(31)*species(142);
    ratelaws(459) = ((expressions(40)*1000)/expressions(138))*species(31)*species(144);
    ratelaws(460) = ((expressions(40)*1000)/expressions(138))*species(31)*species(146);
    ratelaws(461) = ((expressions(40)*1000)/expressions(138))*species(31)*species(148);
    ratelaws(462) = expressions(68)*species(133);
    ratelaws(463) = expressions(68)*species(134);
    ratelaws(464) = ((expressions(43)*1000)/expressions(138))*species(31)*species(137);
    ratelaws(465) = ((expressions(43)*1000)/expressions(138))*species(31)*species(139);
    ratelaws(466) = ((expressions(43)*1000)/expressions(138))*species(31)*species(142);
    ratelaws(467) = ((expressions(43)*1000)/expressions(138))*species(31)*species(144);
    ratelaws(468) = ((expressions(43)*1000)/expressions(138))*species(31)*species(146);
    ratelaws(469) = ((expressions(43)*1000)/expressions(138))*species(31)*species(148);
    ratelaws(470) = expressions(71)*species(135);
    ratelaws(471) = expressions(71)*species(136);
    ratelaws(472) = expressions(71)*species(138);
    ratelaws(473) = expressions(71)*species(140);
    ratelaws(474) = (expressions(92)/expressions(10))*species(4)*species(121);
    ratelaws(475) = (expressions(92)/expressions(10))*species(4)*species(125);
    ratelaws(476) = (expressions(92)/expressions(10))*species(4)*species(131);
    ratelaws(477) = (expressions(92)/expressions(10))*species(4)*species(141);
    ratelaws(478) = (expressions(92)/expressions(10))*species(4)*species(143);
    ratelaws(479) = expressions(102)*species(133);
    ratelaws(480) = expressions(102)*species(137);
    ratelaws(481) = expressions(102)*species(142);
    ratelaws(482) = expressions(102)*species(144);
    ratelaws(483) = (expressions(92)/expressions(10))*species(68)*species(121);
    ratelaws(484) = (expressions(92)/expressions(10))*species(68)*species(125);
    ratelaws(485) = expressions(102)*species(135);
    ratelaws(486) = expressions(102)*species(138);
    ratelaws(487) = (expressions(92)/expressions(11))*species(8)*species(122);
    ratelaws(488) = (expressions(92)/expressions(11))*species(8)*species(126);
    ratelaws(489) = (expressions(92)/expressions(11))*species(8)*species(132);
    ratelaws(490) = (expressions(92)/expressions(11))*species(8)*species(145);
    ratelaws(491) = (expressions(92)/expressions(11))*species(8)*species(147);
    ratelaws(492) = expressions(102)*species(134);
    ratelaws(493) = expressions(102)*species(139);
    ratelaws(494) = expressions(102)*species(146);
    ratelaws(495) = expressions(102)*species(148);
    ratelaws(496) = (expressions(92)/expressions(11))*species(69)*species(122);
    ratelaws(497) = (expressions(92)/expressions(11))*species(69)*species(126);
    ratelaws(498) = expressions(102)*species(136);
    ratelaws(499) = expressions(102)*species(140);
    ratelaws(500) = (expressions(91)/expressions(10))*species(12)*species(133);
    ratelaws(501) = (expressions(91)/expressions(10))*species(12)*species(135);
    ratelaws(502) = (expressions(91)/expressions(10))*species(49)*species(133);
    ratelaws(503) = (expressions(91)/expressions(10))*species(49)*species(135);
    ratelaws(504) = (expressions(91)/expressions(10))*species(51)*species(133);
    ratelaws(505) = (expressions(91)/expressions(10))*species(51)*species(135);
    ratelaws(506) = expressions(103)*species(121);
    ratelaws(507) = expressions(103)*species(125);
    ratelaws(508) = expressions(103)*species(131);
    ratelaws(509) = expressions(103)*species(137);
    ratelaws(510) = expressions(103)*species(138);
    ratelaws(511) = expressions(103)*species(141);
    ratelaws(512) = expressions(103)*species(142);
    ratelaws(513) = expressions(103)*species(143);
    ratelaws(514) = expressions(103)*species(144);
    ratelaws(515) = (expressions(91)/expressions(11))*species(16)*species(134);
    ratelaws(516) = (expressions(91)/expressions(11))*species(16)*species(136);
    ratelaws(517) = (expressions(91)/expressions(11))*species(50)*species(134);
    ratelaws(518) = (expressions(91)/expressions(11))*species(50)*species(136);
    ratelaws(519) = (expressions(91)/expressions(11))*species(52)*species(134);
    ratelaws(520) = (expressions(91)/expressions(11))*species(52)*species(136);
    ratelaws(521) = expressions(103)*species(122);
    ratelaws(522) = expressions(103)*species(126);
    ratelaws(523) = expressions(103)*species(132);
    ratelaws(524) = expressions(103)*species(139);
    ratelaws(525) = expressions(103)*species(140);
    ratelaws(526) = expressions(103)*species(145);
    ratelaws(527) = expressions(103)*species(146);
    ratelaws(528) = expressions(103)*species(147);
    ratelaws(529) = expressions(103)*species(148);
    ratelaws(530) = expressions(70)*species(149);
    ratelaws(531) = expressions(70)*species(150);
    ratelaws(532) = expressions(70)*species(151);
    ratelaws(533) = expressions(70)*species(152);
    ratelaws(534) = expressions(70)*species(153);
    ratelaws(535) = expressions(70)*species(154);
    ratelaws(536) = expressions(72)*species(149);
    ratelaws(537) = expressions(72)*species(150);
    ratelaws(538) = expressions(72)*species(151);
    ratelaws(539) = expressions(72)*species(152);
    ratelaws(540) = expressions(27)*species(155);
    ratelaws(541) = expressions(27)*species(161);
    ratelaws(542) = expressions(27)*species(156);
    ratelaws(543) = expressions(27)*species(162);
    ratelaws(544) = expressions(27)*species(157);
    ratelaws(545) = expressions(27)*species(158);
    ratelaws(546) = expressions(27)*species(159);
    ratelaws(547) = expressions(27)*species(160);
    ratelaws(548) = expressions(27)*species(163);
    ratelaws(549) = expressions(27)*species(164);
    ratelaws(550) = expressions(27)*species(165);
    ratelaws(551) = expressions(27)*species(166);
    ratelaws(552) = expressions(27)*species(167);
    ratelaws(553) = expressions(27)*species(168);
    ratelaws(554) = expressions(101)*species(155);
    ratelaws(555) = expressions(101)*species(156);
    ratelaws(556) = expressions(80)*species(155);
    ratelaws(557) = expressions(80)*species(156);
    ratelaws(558) = expressions(100)*species(155);
    ratelaws(559) = expressions(100)*species(156);
    ratelaws(560) = ((expressions(53)*1000)/expressions(138))*species(33)*species(163);
    ratelaws(561) = ((expressions(53)*1000)/expressions(138))*species(33)*species(164);
    ratelaws(562) = expressions(81)*species(157);
    ratelaws(563) = expressions(81)*species(158);
    ratelaws(564) = expressions(81)*species(165);
    ratelaws(565) = expressions(81)*species(167);
    ratelaws(566) = ((expressions(51)*1000)/expressions(138))*species(32)*species(163);
    ratelaws(567) = ((expressions(51)*1000)/expressions(138))*species(32)*species(164);
    ratelaws(568) = expressions(79)*species(159);
    ratelaws(569) = expressions(79)*species(160);
    ratelaws(570) = expressions(79)*species(166);
    ratelaws(571) = expressions(79)*species(168);
    ratelaws(572) = expressions(99)*species(161);
    ratelaws(573) = expressions(99)*species(162);
    ratelaws(574) = expressions(68)*species(163);
    ratelaws(575) = expressions(68)*species(164);
    ratelaws(576) = expressions(68)*species(165);
    ratelaws(577) = expressions(68)*species(166);
    ratelaws(578) = expressions(68)*species(167);
    ratelaws(579) = expressions(68)*species(168);
    ratelaws(580) = expressions(71)*species(157);
    ratelaws(581) = expressions(71)*species(158);
    ratelaws(582) = expressions(71)*species(159);
    ratelaws(583) = expressions(71)*species(160);
    ratelaws(584) = expressions(102)*species(163);
    ratelaws(585) = expressions(102)*species(165);
    ratelaws(586) = expressions(102)*species(166);
    ratelaws(587) = expressions(102)*species(157);
    ratelaws(588) = expressions(102)*species(159);
    ratelaws(589) = expressions(102)*species(164);
    ratelaws(590) = expressions(102)*species(167);
    ratelaws(591) = expressions(102)*species(168);
    ratelaws(592) = expressions(102)*species(158);
    ratelaws(593) = expressions(102)*species(160);
    ratelaws(594) = expressions(103)*species(157);
    ratelaws(595) = expressions(103)*species(159);
    ratelaws(596) = expressions(103)*species(163);
    ratelaws(597) = expressions(103)*species(165);
    ratelaws(598) = expressions(103)*species(166);
    ratelaws(599) = expressions(103)*species(158);
    ratelaws(600) = expressions(103)*species(160);
    ratelaws(601) = expressions(103)*species(164);
    ratelaws(602) = expressions(103)*species(167);
    ratelaws(603) = expressions(103)*species(168);

end

% Calculate species derivatives
function [ Dspecies ] = calc_species_deriv ( time, species, expressions )
    
    % initialize derivative vector
    Dspecies = zeros(168,1);
    
    % update observables
    [ observables ] = calc_observables( species, expressions );
    
    % update ratelaws
    [ ratelaws ] = calc_ratelaws( species, expressions, observables );
                        
    % calculate derivatives
    Dspecies(1) = ratelaws(23) +ratelaws(24) +ratelaws(25) +ratelaws(26) +ratelaws(27) +ratelaws(28) +ratelaws(29) +ratelaws(30) +ratelaws(31) +ratelaws(32) +ratelaws(33) +ratelaws(34) +ratelaws(35) +ratelaws(36) +ratelaws(37) +ratelaws(38) +ratelaws(39) +ratelaws(40) +ratelaws(41) +ratelaws(42) +ratelaws(43) +ratelaws(44) +ratelaws(45) +ratelaws(46) +ratelaws(49) +ratelaws(50) +ratelaws(52) +ratelaws(54) +ratelaws(57) +ratelaws(58) +ratelaws(59) +ratelaws(62) +ratelaws(63) +ratelaws(64) +ratelaws(137) +ratelaws(138) +ratelaws(139) +ratelaws(140) +ratelaws(141) +ratelaws(142) +ratelaws(143) +ratelaws(144) +ratelaws(145) +ratelaws(146) +ratelaws(147) +ratelaws(148) +ratelaws(149) +ratelaws(150) +ratelaws(151) +ratelaws(152) +ratelaws(153) +ratelaws(154) +ratelaws(155) +ratelaws(156) +ratelaws(157) +ratelaws(158) +ratelaws(159) +ratelaws(160) +ratelaws(161) +ratelaws(162) +ratelaws(163) +ratelaws(164) +ratelaws(165) +ratelaws(166) +ratelaws(167) +ratelaws(168) +ratelaws(169) +ratelaws(170) +ratelaws(171) +ratelaws(172) +ratelaws(173) +ratelaws(174) +ratelaws(175) +ratelaws(176) +ratelaws(177) +ratelaws(178) +ratelaws(179) +ratelaws(180) +ratelaws(181) +ratelaws(182) +ratelaws(183) +ratelaws(184) +ratelaws(185) +ratelaws(239) +ratelaws(240) +ratelaws(241) +ratelaws(303) +ratelaws(324) +ratelaws(340) +ratelaws(341) +ratelaws(342) +ratelaws(343) +ratelaws(344) +ratelaws(345) +ratelaws(346) +ratelaws(347) +ratelaws(348) +ratelaws(349) +ratelaws(350) +ratelaws(351) +ratelaws(352) +ratelaws(353) +ratelaws(354) +ratelaws(355) +ratelaws(356) +ratelaws(357) +ratelaws(358) +ratelaws(359) +ratelaws(360) +ratelaws(361) +ratelaws(362) +ratelaws(363) +ratelaws(364) +ratelaws(365) +ratelaws(366) +ratelaws(367) +ratelaws(368) +ratelaws(369) +ratelaws(370) +ratelaws(371) +ratelaws(372) +ratelaws(373) +ratelaws(374) +ratelaws(375) +ratelaws(376) +ratelaws(377) +ratelaws(378) +ratelaws(379) +ratelaws(380) +ratelaws(381) +ratelaws(382) +ratelaws(383) +ratelaws(384) +ratelaws(385) +ratelaws(540) +ratelaws(541) +ratelaws(542) +ratelaws(543) +ratelaws(544) +ratelaws(545) +ratelaws(546) +ratelaws(547) +ratelaws(548) +ratelaws(549) +ratelaws(550) +ratelaws(551) +ratelaws(552) +ratelaws(553);
    Dspecies(2) = 0.0;
    Dspecies(3) = ratelaws(1) -ratelaws(27) -ratelaws(90) -ratelaws(94) -ratelaws(96) +ratelaws(248) +ratelaws(258) -ratelaws(259) -ratelaws(268) -ratelaws(271) +ratelaws(274) +ratelaws(485) +ratelaws(506) +ratelaws(507);
    Dspecies(4) = ratelaws(2) -ratelaws(25) -ratelaws(92) -ratelaws(94) +ratelaws(250) -ratelaws(256) -ratelaws(257) +ratelaws(258) -ratelaws(474) -ratelaws(475) -ratelaws(476) -ratelaws(477) -ratelaws(478) +ratelaws(479) +ratelaws(480) +ratelaws(481) +ratelaws(482) +ratelaws(584) +ratelaws(585) +ratelaws(586);
    Dspecies(5) = ratelaws(3) -ratelaws(29) -ratelaws(86) -ratelaws(88) +ratelaws(242) +ratelaws(244);
    Dspecies(6) = ratelaws(4) -ratelaws(31) -ratelaws(98) -ratelaws(106) -ratelaws(133) -ratelaws(284) -ratelaws(286) -ratelaws(288) +ratelaws(290) +ratelaws(301) +ratelaws(336) +ratelaws(530) +ratelaws(532) +ratelaws(534);
    Dspecies(7) = ratelaws(5) -ratelaws(28) -ratelaws(91) -ratelaws(95) -ratelaws(97) +ratelaws(249) +ratelaws(263) -ratelaws(264) -ratelaws(277) -ratelaws(280) +ratelaws(283) +ratelaws(498) +ratelaws(521) +ratelaws(522);
    Dspecies(8) = ratelaws(6) -ratelaws(26) -ratelaws(93) -ratelaws(95) +ratelaws(251) -ratelaws(261) -ratelaws(262) +ratelaws(263) -ratelaws(487) -ratelaws(488) -ratelaws(489) -ratelaws(490) -ratelaws(491) +ratelaws(492) +ratelaws(493) +ratelaws(494) +ratelaws(495) +ratelaws(589) +ratelaws(590) +ratelaws(591);
    Dspecies(9) = ratelaws(7) -ratelaws(30) -ratelaws(87) -ratelaws(89) +ratelaws(243) +ratelaws(245);
    Dspecies(10) = ratelaws(8) -ratelaws(32) -ratelaws(99) -ratelaws(107) -ratelaws(134) -ratelaws(285) -ratelaws(287) -ratelaws(289) +ratelaws(291) +ratelaws(302) +ratelaws(337) +ratelaws(531) +ratelaws(533) +ratelaws(535);
    Dspecies(11) = ratelaws(9) -ratelaws(33) -ratelaws(65) -ratelaws(67) -ratelaws(69) -ratelaws(71) +ratelaws(189) +ratelaws(196) +ratelaws(202) +ratelaws(205);
    Dspecies(12) = ratelaws(10) -ratelaws(35) -ratelaws(73) -ratelaws(75) -ratelaws(88) -ratelaws(96) +ratelaws(214) +ratelaws(220) +ratelaws(244) -ratelaws(266) -ratelaws(267) +ratelaws(274) -ratelaws(444) -ratelaws(500) -ratelaws(501) +ratelaws(508) +ratelaws(509) +ratelaws(510) +ratelaws(572) +ratelaws(596);
    Dspecies(13) = ratelaws(11) -ratelaws(39) -ratelaws(71) -ratelaws(79) -ratelaws(81) -ratelaws(204) +ratelaws(205) -ratelaws(208) +ratelaws(224) +ratelaws(228) -ratelaws(230) +ratelaws(402) -ratelaws(404) +ratelaws(405) +ratelaws(438) +ratelaws(558);
    Dspecies(14) = ratelaws(12) -ratelaws(37);
    Dspecies(15) = ratelaws(13) -ratelaws(34) -ratelaws(66) -ratelaws(68) -ratelaws(70) -ratelaws(72) +ratelaws(193) +ratelaws(197) +ratelaws(203) +ratelaws(207);
    Dspecies(16) = ratelaws(14) -ratelaws(36) -ratelaws(74) -ratelaws(76) -ratelaws(89) -ratelaws(97) +ratelaws(215) +ratelaws(221) +ratelaws(245) -ratelaws(275) -ratelaws(276) +ratelaws(283) -ratelaws(445) -ratelaws(515) -ratelaws(516) +ratelaws(523) +ratelaws(524) +ratelaws(525) +ratelaws(573) +ratelaws(601);
    Dspecies(17) = ratelaws(15) -ratelaws(40) -ratelaws(72) -ratelaws(80) -ratelaws(82) -ratelaws(206) +ratelaws(207) -ratelaws(209) +ratelaws(225) +ratelaws(229) -ratelaws(231) +ratelaws(403) -ratelaws(406) +ratelaws(407) +ratelaws(439) +ratelaws(559);
    Dspecies(18) = ratelaws(16) -ratelaws(38);
    Dspecies(19) = ratelaws(17) -ratelaws(23) -ratelaws(65) -ratelaws(77) -ratelaws(81) -ratelaws(100) -ratelaws(120) -ratelaws(128) -ratelaws(186) -ratelaws(187) -ratelaws(188) +ratelaws(189) +ratelaws(222) +ratelaws(228) -ratelaws(232) +ratelaws(296) +ratelaws(318) +ratelaws(331) -ratelaws(386) +ratelaws(387) +ratelaws(388) +ratelaws(389) +ratelaws(440) +ratelaws(554);
    Dspecies(20) = ratelaws(18) -ratelaws(24) -ratelaws(66) -ratelaws(78) -ratelaws(82) -ratelaws(101) -ratelaws(121) -ratelaws(129) -ratelaws(190) -ratelaws(191) -ratelaws(192) +ratelaws(193) +ratelaws(223) +ratelaws(229) -ratelaws(233) +ratelaws(297) +ratelaws(319) +ratelaws(332) -ratelaws(390) +ratelaws(391) +ratelaws(392) +ratelaws(393) +ratelaws(441) +ratelaws(555);
    Dspecies(21) = -ratelaws(122) -ratelaws(130) +ratelaws(320) +ratelaws(333);
    Dspecies(22) = ratelaws(19) -ratelaws(41) -ratelaws(123) +ratelaws(321) -ratelaws(325) +ratelaws(326);
    Dspecies(23) = ratelaws(20) -ratelaws(42) -ratelaws(124) +ratelaws(322) -ratelaws(327) +ratelaws(328);
    Dspecies(24) = -ratelaws(43) -ratelaws(126) +ratelaws(329);
    Dspecies(25) = -ratelaws(44) -ratelaws(127) +ratelaws(330);
    Dspecies(26) = ratelaws(21) -ratelaws(45) -ratelaws(131) +ratelaws(334);
    Dspecies(27) = ratelaws(22) -ratelaws(46) -ratelaws(132) +ratelaws(335);
    Dspecies(28) = -ratelaws(108) -ratelaws(111) -ratelaws(114) -ratelaws(117) +ratelaws(304) +ratelaws(307) +ratelaws(310) +ratelaws(313) +ratelaws(316) +ratelaws(317);
    Dspecies(29) = -ratelaws(109) -ratelaws(112) -ratelaws(115) -ratelaws(118) +ratelaws(305) +ratelaws(308) +ratelaws(311) +ratelaws(314);
    Dspecies(30) = -ratelaws(110) -ratelaws(113) -ratelaws(116) -ratelaws(119) +ratelaws(306) +ratelaws(309) +ratelaws(312) +ratelaws(315);
    Dspecies(31) = ratelaws(47) -ratelaws(48) -ratelaws(49) -ratelaws(83) -ratelaws(84) -ratelaws(85) -ratelaws(86) -ratelaws(87) -ratelaws(90) -ratelaws(91) -ratelaws(92) -ratelaws(93) -ratelaws(98) -ratelaws(99) -ratelaws(100) -ratelaws(101) -ratelaws(103) -ratelaws(108) -ratelaws(109) -ratelaws(110) -ratelaws(125) +ratelaws(236) +ratelaws(237) +ratelaws(238) +ratelaws(242) +ratelaws(243) -ratelaws(246) -ratelaws(247) +ratelaws(248) +ratelaws(249) +ratelaws(250) +ratelaws(251) -ratelaws(252) -ratelaws(253) -ratelaws(254) -ratelaws(255) +ratelaws(290) +ratelaws(291) +ratelaws(296) +ratelaws(297) +ratelaws(300) +ratelaws(304) +ratelaws(305) +ratelaws(306) +ratelaws(323) -ratelaws(446) -ratelaws(447) -ratelaws(448) -ratelaws(449) +ratelaws(450) +ratelaws(451) +ratelaws(452) +ratelaws(453) +ratelaws(454) +ratelaws(455) -ratelaws(456) -ratelaws(457) -ratelaws(458) -ratelaws(459) -ratelaws(460) -ratelaws(461) +ratelaws(462) +ratelaws(463) -ratelaws(464) -ratelaws(465) -ratelaws(466) -ratelaws(467) -ratelaws(468) -ratelaws(469) +ratelaws(470) +ratelaws(471) +ratelaws(472) +ratelaws(473) +ratelaws(574) +ratelaws(575) +ratelaws(576) +ratelaws(577) +ratelaws(578) +ratelaws(579) +ratelaws(580) +ratelaws(581) +ratelaws(582) +ratelaws(583);
    Dspecies(32) = ratelaws(55) -ratelaws(57) -ratelaws(67) -ratelaws(68) -ratelaws(75) -ratelaws(76) -ratelaws(77) -ratelaws(78) -ratelaws(79) -ratelaws(80) -ratelaws(83) -ratelaws(104) -ratelaws(105) -ratelaws(111) -ratelaws(112) -ratelaws(113) -ratelaws(135) -ratelaws(194) -ratelaws(195) +ratelaws(196) +ratelaws(197) -ratelaws(216) -ratelaws(217) -ratelaws(218) -ratelaws(219) +ratelaws(220) +ratelaws(221) +ratelaws(222) +ratelaws(223) +ratelaws(224) +ratelaws(225) -ratelaws(226) -ratelaws(227) +ratelaws(236) -ratelaws(292) -ratelaws(293) +ratelaws(307) +ratelaws(308) +ratelaws(309) +ratelaws(338) +ratelaws(394) +ratelaws(395) -ratelaws(422) -ratelaws(423) -ratelaws(424) -ratelaws(425) -ratelaws(426) -ratelaws(427) +ratelaws(428) +ratelaws(429) +ratelaws(430) +ratelaws(431) +ratelaws(432) +ratelaws(433) +ratelaws(434) +ratelaws(435) +ratelaws(436) +ratelaws(437) +ratelaws(536) +ratelaws(537) -ratelaws(566) -ratelaws(567) +ratelaws(568) +ratelaws(569) +ratelaws(570) +ratelaws(571);
    Dspecies(33) = ratelaws(56) -ratelaws(58) -ratelaws(69) -ratelaws(70) -ratelaws(73) -ratelaws(74) -ratelaws(84) -ratelaws(198) -ratelaws(199) -ratelaws(200) -ratelaws(201) +ratelaws(202) +ratelaws(203) -ratelaws(210) -ratelaws(211) -ratelaws(212) -ratelaws(213) +ratelaws(214) +ratelaws(215) +ratelaws(237) -ratelaws(294) -ratelaws(295) -ratelaws(396) -ratelaws(397) +ratelaws(398) +ratelaws(399) +ratelaws(400) +ratelaws(401) -ratelaws(408) -ratelaws(409) -ratelaws(410) -ratelaws(411) -ratelaws(412) -ratelaws(413) +ratelaws(414) +ratelaws(415) +ratelaws(416) +ratelaws(417) +ratelaws(418) +ratelaws(419) +ratelaws(420) +ratelaws(421) +ratelaws(538) +ratelaws(539) +ratelaws(556) +ratelaws(557) -ratelaws(560) -ratelaws(561) +ratelaws(562) +ratelaws(563) +ratelaws(564) +ratelaws(565);
    Dspecies(34) = -ratelaws(59) -ratelaws(85) +ratelaws(104) +ratelaws(105) +ratelaws(238) +ratelaws(316) +ratelaws(317);
    Dspecies(35) = ratelaws(60) -ratelaws(64) -ratelaws(102) +ratelaws(298);
    Dspecies(36) = -ratelaws(63) -ratelaws(106) -ratelaws(107) +ratelaws(299) +ratelaws(301) +ratelaws(302);
    Dspecies(37) = ratelaws(61) -ratelaws(62) -ratelaws(102) -ratelaws(103) +ratelaws(298) +ratelaws(299) +ratelaws(300);
    Dspecies(38) = ratelaws(48) -ratelaws(50);
    Dspecies(39) = ratelaws(51) -ratelaws(52) -ratelaws(114) -ratelaws(115) -ratelaws(116) -ratelaws(120) -ratelaws(121) -ratelaws(122) -ratelaws(123) -ratelaws(124) -ratelaws(125) -ratelaws(136) +ratelaws(310) +ratelaws(311) +ratelaws(312) +ratelaws(318) +ratelaws(319) +ratelaws(320) +ratelaws(321) +ratelaws(322) +ratelaws(323) +ratelaws(339);
    Dspecies(40) = ratelaws(53) -ratelaws(54) -ratelaws(117) -ratelaws(118) -ratelaws(119) -ratelaws(128) -ratelaws(129) -ratelaws(130) -ratelaws(131) -ratelaws(132) -ratelaws(133) -ratelaws(134) -ratelaws(135) -ratelaws(136) +ratelaws(313) +ratelaws(314) +ratelaws(315) +ratelaws(331) +ratelaws(332) +ratelaws(333) +ratelaws(334) +ratelaws(335) +ratelaws(336) +ratelaws(337) +ratelaws(338) +ratelaws(339);
    Dspecies(41) = ratelaws(65) -ratelaws(137) -ratelaws(189) -ratelaws(194) -ratelaws(198) -ratelaws(204) +ratelaws(394) +ratelaws(398) +ratelaws(402);
    Dspecies(42) = ratelaws(66) -ratelaws(144) -ratelaws(193) -ratelaws(195) -ratelaws(199) -ratelaws(206) +ratelaws(395) +ratelaws(399) +ratelaws(403);
    Dspecies(43) = ratelaws(67) -ratelaws(169) -ratelaws(186) -ratelaws(196) +ratelaws(387);
    Dspecies(44) = ratelaws(68) -ratelaws(170) -ratelaws(190) -ratelaws(197) +ratelaws(391);
    Dspecies(45) = ratelaws(69) -ratelaws(171) -ratelaws(187) -ratelaws(202) -ratelaws(208) +ratelaws(388) +ratelaws(405);
    Dspecies(46) = ratelaws(70) -ratelaws(172) -ratelaws(191) -ratelaws(203) -ratelaws(209) +ratelaws(392) +ratelaws(407);
    Dspecies(47) = ratelaws(71) -ratelaws(173) -ratelaws(188) -ratelaws(200) -ratelaws(205) +ratelaws(389) +ratelaws(400);
    Dspecies(48) = ratelaws(72) -ratelaws(174) -ratelaws(192) -ratelaws(201) -ratelaws(207) +ratelaws(393) +ratelaws(401);
    Dspecies(49) = ratelaws(73) -ratelaws(175) -ratelaws(214) -ratelaws(268) -ratelaws(269) -ratelaws(270) -ratelaws(502) -ratelaws(503) +ratelaws(506) +ratelaws(511) +ratelaws(512) +ratelaws(594) +ratelaws(597);
    Dspecies(50) = ratelaws(74) -ratelaws(176) -ratelaws(215) -ratelaws(277) -ratelaws(278) -ratelaws(279) -ratelaws(517) -ratelaws(518) +ratelaws(521) +ratelaws(526) +ratelaws(527) +ratelaws(599) +ratelaws(602);
    Dspecies(51) = ratelaws(75) -ratelaws(177) -ratelaws(220) -ratelaws(234) -ratelaws(271) -ratelaws(272) -ratelaws(273) +ratelaws(442) -ratelaws(504) -ratelaws(505) +ratelaws(507) +ratelaws(513) +ratelaws(514) +ratelaws(595) +ratelaws(598);
    Dspecies(52) = ratelaws(76) -ratelaws(178) -ratelaws(221) -ratelaws(235) -ratelaws(280) -ratelaws(281) -ratelaws(282) +ratelaws(443) -ratelaws(519) -ratelaws(520) +ratelaws(522) +ratelaws(528) +ratelaws(529) +ratelaws(600) +ratelaws(603);
    Dspecies(53) = ratelaws(77) -ratelaws(138) -ratelaws(222) -ratelaws(230) +ratelaws(438);
    Dspecies(54) = ratelaws(78) -ratelaws(145) -ratelaws(223) -ratelaws(231) +ratelaws(439);
    Dspecies(55) = ratelaws(79) -ratelaws(179) -ratelaws(224) -ratelaws(232) +ratelaws(440);
    Dspecies(56) = ratelaws(80) -ratelaws(180) -ratelaws(225) -ratelaws(233) +ratelaws(441);
    Dspecies(57) = ratelaws(81) -ratelaws(139) -ratelaws(226) -ratelaws(228) -ratelaws(234) +ratelaws(436) +ratelaws(442);
    Dspecies(58) = ratelaws(82) -ratelaws(146) -ratelaws(227) -ratelaws(229) -ratelaws(235) +ratelaws(437) +ratelaws(443);
    Dspecies(59) = ratelaws(83) -ratelaws(236) -ratelaws(239) -ratelaws(284) -ratelaws(285) +ratelaws(530) +ratelaws(531);
    Dspecies(60) = ratelaws(84) -ratelaws(237) -ratelaws(240) -ratelaws(286) -ratelaws(287) +ratelaws(532) +ratelaws(533);
    Dspecies(61) = ratelaws(85) -ratelaws(238) -ratelaws(241) -ratelaws(288) -ratelaws(289) +ratelaws(534) +ratelaws(535);
    Dspecies(62) = ratelaws(86) -ratelaws(159) -ratelaws(242);
    Dspecies(63) = ratelaws(87) -ratelaws(160) -ratelaws(243);
    Dspecies(64) = ratelaws(88) -ratelaws(161) -ratelaws(210) -ratelaws(216) -ratelaws(244) +ratelaws(414) +ratelaws(428);
    Dspecies(65) = ratelaws(89) -ratelaws(162) -ratelaws(211) -ratelaws(217) -ratelaws(245) +ratelaws(415) +ratelaws(429);
    Dspecies(66) = ratelaws(90) -ratelaws(155) -ratelaws(248) -ratelaws(256) -ratelaws(266) -ratelaws(269) -ratelaws(272) +ratelaws(479) +ratelaws(508) +ratelaws(511) +ratelaws(513);
    Dspecies(67) = ratelaws(91) -ratelaws(156) -ratelaws(249) -ratelaws(261) -ratelaws(275) -ratelaws(278) -ratelaws(281) +ratelaws(492) +ratelaws(523) +ratelaws(526) +ratelaws(528);
    Dspecies(68) = ratelaws(92) -ratelaws(151) -ratelaws(250) -ratelaws(259) -ratelaws(260) -ratelaws(483) -ratelaws(484) +ratelaws(485) +ratelaws(486) +ratelaws(587) +ratelaws(588);
    Dspecies(69) = ratelaws(93) -ratelaws(152) -ratelaws(251) -ratelaws(264) -ratelaws(265) -ratelaws(496) -ratelaws(497) +ratelaws(498) +ratelaws(499) +ratelaws(592) +ratelaws(593);
    Dspecies(70) = ratelaws(94) -ratelaws(153) -ratelaws(252) -ratelaws(254) -ratelaws(258) -ratelaws(267) -ratelaws(270) -ratelaws(273) +ratelaws(462) +ratelaws(470) +ratelaws(509) +ratelaws(512) +ratelaws(514);
    Dspecies(71) = ratelaws(95) -ratelaws(154) -ratelaws(253) -ratelaws(255) -ratelaws(263) -ratelaws(276) -ratelaws(279) -ratelaws(282) +ratelaws(463) +ratelaws(471) +ratelaws(524) +ratelaws(527) +ratelaws(529);
    Dspecies(72) = ratelaws(96) -ratelaws(157) -ratelaws(212) -ratelaws(218) -ratelaws(246) -ratelaws(257) -ratelaws(260) -ratelaws(274) +ratelaws(416) +ratelaws(430) +ratelaws(450) +ratelaws(480) +ratelaws(486);
    Dspecies(73) = ratelaws(97) -ratelaws(158) -ratelaws(213) -ratelaws(219) -ratelaws(247) -ratelaws(262) -ratelaws(265) -ratelaws(283) +ratelaws(417) +ratelaws(431) +ratelaws(451) +ratelaws(493) +ratelaws(499);
    Dspecies(74) = ratelaws(98) -ratelaws(163) -ratelaws(290) -ratelaws(292) -ratelaws(294) +ratelaws(536) +ratelaws(538);
    Dspecies(75) = ratelaws(99) -ratelaws(164) -ratelaws(291) -ratelaws(293) -ratelaws(295) +ratelaws(537) +ratelaws(539);
    Dspecies(76) = ratelaws(100) -ratelaws(140) -ratelaws(296);
    Dspecies(77) = ratelaws(101) -ratelaws(147) -ratelaws(297);
    Dspecies(78) = ratelaws(102) -ratelaws(185) -ratelaws(298) -ratelaws(299);
    Dspecies(79) = ratelaws(103) -ratelaws(300) -ratelaws(303);
    Dspecies(80) = ratelaws(106) -ratelaws(165) -ratelaws(301);
    Dspecies(81) = ratelaws(107) -ratelaws(166) -ratelaws(302);
    Dspecies(82) = ratelaws(108) -ratelaws(304);
    Dspecies(83) = ratelaws(109) -ratelaws(305);
    Dspecies(84) = ratelaws(110) -ratelaws(306);
    Dspecies(85) = ratelaws(111) -ratelaws(307) -ratelaws(316) -ratelaws(317);
    Dspecies(86) = ratelaws(112) -ratelaws(308);
    Dspecies(87) = ratelaws(113) -ratelaws(309);
    Dspecies(88) = ratelaws(114) -ratelaws(310);
    Dspecies(89) = ratelaws(115) -ratelaws(311);
    Dspecies(90) = ratelaws(116) -ratelaws(312);
    Dspecies(91) = ratelaws(117) -ratelaws(313);
    Dspecies(92) = ratelaws(118) -ratelaws(314);
    Dspecies(93) = ratelaws(119) -ratelaws(315);
    Dspecies(94) = ratelaws(120) -ratelaws(141) -ratelaws(318) -ratelaws(325) +ratelaws(326);
    Dspecies(95) = ratelaws(121) -ratelaws(148) -ratelaws(319) -ratelaws(327) +ratelaws(328);
    Dspecies(96) = ratelaws(122) -ratelaws(320);
    Dspecies(97) = ratelaws(123) -ratelaws(181) -ratelaws(321);
    Dspecies(98) = ratelaws(124) -ratelaws(182) -ratelaws(322);
    Dspecies(99) = ratelaws(125) -ratelaws(323) -ratelaws(324);
    Dspecies(100) = 2.0*ratelaws(126) -ratelaws(142) +ratelaws(325) -ratelaws(326) -2.0*ratelaws(329);
    Dspecies(101) = 2.0*ratelaws(127) -ratelaws(149) +ratelaws(327) -ratelaws(328) -2.0*ratelaws(330);
    Dspecies(102) = ratelaws(128) -ratelaws(143) -ratelaws(331);
    Dspecies(103) = ratelaws(129) -ratelaws(150) -ratelaws(332);
    Dspecies(104) = ratelaws(130) -ratelaws(333);
    Dspecies(105) = ratelaws(131) -ratelaws(183) -ratelaws(334);
    Dspecies(106) = ratelaws(132) -ratelaws(184) -ratelaws(335);
    Dspecies(107) = ratelaws(133) -ratelaws(167) -ratelaws(336);
    Dspecies(108) = ratelaws(134) -ratelaws(168) -ratelaws(337);
    Dspecies(109) = ratelaws(135) -ratelaws(338) -ratelaws(340);
    Dspecies(110) = ratelaws(136) -ratelaws(339) -ratelaws(341);
    Dspecies(111) = ratelaws(186) +ratelaws(194) -ratelaws(342) -ratelaws(387) -ratelaws(394);
    Dspecies(112) = ratelaws(187) +ratelaws(198) -ratelaws(343) -ratelaws(388) -ratelaws(398) -ratelaws(404) +ratelaws(558);
    Dspecies(113) = ratelaws(188) +ratelaws(204) -ratelaws(344) -ratelaws(389) -ratelaws(396) -ratelaws(402) +ratelaws(556);
    Dspecies(114) = ratelaws(190) +ratelaws(195) -ratelaws(347) -ratelaws(391) -ratelaws(395);
    Dspecies(115) = ratelaws(191) +ratelaws(199) -ratelaws(348) -ratelaws(392) -ratelaws(399) -ratelaws(406) +ratelaws(559);
    Dspecies(116) = ratelaws(192) +ratelaws(206) -ratelaws(349) -ratelaws(393) -ratelaws(397) -ratelaws(403) +ratelaws(557);
    Dspecies(117) = ratelaws(200) +ratelaws(208) -ratelaws(384) -ratelaws(386) -ratelaws(400) -ratelaws(405) +ratelaws(554);
    Dspecies(118) = ratelaws(201) +ratelaws(209) -ratelaws(385) -ratelaws(390) -ratelaws(401) -ratelaws(407) +ratelaws(555);
    Dspecies(119) = ratelaws(210) -ratelaws(374) -ratelaws(414);
    Dspecies(120) = ratelaws(211) -ratelaws(375) -ratelaws(415);
    Dspecies(121) = ratelaws(212) +ratelaws(268) -ratelaws(364) -ratelaws(416) -ratelaws(446) +ratelaws(452) -ratelaws(474) +ratelaws(481) -ratelaws(483) -ratelaws(506) +ratelaws(587);
    Dspecies(122) = ratelaws(213) +ratelaws(277) -ratelaws(365) -ratelaws(417) -ratelaws(447) +ratelaws(454) -ratelaws(487) +ratelaws(494) -ratelaws(496) -ratelaws(521) +ratelaws(592);
    Dspecies(123) = ratelaws(216) -ratelaws(376) -ratelaws(428);
    Dspecies(124) = ratelaws(217) -ratelaws(377) -ratelaws(429);
    Dspecies(125) = ratelaws(218) +ratelaws(271) -ratelaws(366) -ratelaws(430) -ratelaws(448) +ratelaws(453) -ratelaws(475) +ratelaws(482) -ratelaws(484) -ratelaws(507) +ratelaws(588);
    Dspecies(126) = ratelaws(219) +ratelaws(280) -ratelaws(367) -ratelaws(431) -ratelaws(449) +ratelaws(455) -ratelaws(488) +ratelaws(495) -ratelaws(497) -ratelaws(522) +ratelaws(593);
    Dspecies(127) = ratelaws(226) +ratelaws(230) +ratelaws(232) -ratelaws(345) -ratelaws(436) -ratelaws(438) -ratelaws(440) -ratelaws(444) +ratelaws(572);
    Dspecies(128) = ratelaws(227) +ratelaws(231) +ratelaws(233) -ratelaws(350) -ratelaws(437) -ratelaws(439) -ratelaws(441) -ratelaws(445) +ratelaws(573);
    Dspecies(129) = ratelaws(234) -ratelaws(346) -ratelaws(442);
    Dspecies(130) = ratelaws(235) -ratelaws(351) -ratelaws(443);
    Dspecies(131) = ratelaws(246) +ratelaws(266) -ratelaws(368) -ratelaws(408) +ratelaws(418) -ratelaws(422) +ratelaws(432) -ratelaws(450) -ratelaws(476) -ratelaws(508) +ratelaws(584);
    Dspecies(132) = ratelaws(247) +ratelaws(275) -ratelaws(369) -ratelaws(409) +ratelaws(420) -ratelaws(423) +ratelaws(434) -ratelaws(451) -ratelaws(489) -ratelaws(523) +ratelaws(589);
    Dspecies(133) = ratelaws(252) +ratelaws(256) -ratelaws(352) -ratelaws(462) -ratelaws(479) -ratelaws(500) -ratelaws(502) -ratelaws(504) +ratelaws(596) +ratelaws(597) +ratelaws(598);
    Dspecies(134) = ratelaws(253) +ratelaws(261) -ratelaws(353) -ratelaws(463) -ratelaws(492) -ratelaws(515) -ratelaws(517) -ratelaws(519) +ratelaws(601) +ratelaws(602) +ratelaws(603);
    Dspecies(135) = ratelaws(254) +ratelaws(259) -ratelaws(354) -ratelaws(470) -ratelaws(485) -ratelaws(501) -ratelaws(503) -ratelaws(505) +ratelaws(510) +ratelaws(594) +ratelaws(595);
    Dspecies(136) = ratelaws(255) +ratelaws(264) -ratelaws(355) -ratelaws(471) -ratelaws(498) -ratelaws(516) -ratelaws(518) -ratelaws(520) +ratelaws(525) +ratelaws(599) +ratelaws(600);
    Dspecies(137) = ratelaws(257) +ratelaws(267) -ratelaws(356) -ratelaws(410) +ratelaws(419) -ratelaws(424) +ratelaws(433) -ratelaws(456) -ratelaws(464) +ratelaws(472) -ratelaws(480) -ratelaws(509) +ratelaws(574);
    Dspecies(138) = ratelaws(260) -ratelaws(357) -ratelaws(411) -ratelaws(425) +ratelaws(464) -ratelaws(472) -ratelaws(486) +ratelaws(501) -ratelaws(510) +ratelaws(562) +ratelaws(568);
    Dspecies(139) = ratelaws(262) +ratelaws(276) -ratelaws(358) -ratelaws(412) +ratelaws(421) -ratelaws(426) +ratelaws(435) -ratelaws(457) -ratelaws(465) +ratelaws(473) -ratelaws(493) -ratelaws(524) +ratelaws(575);
    Dspecies(140) = ratelaws(265) -ratelaws(359) -ratelaws(413) -ratelaws(427) +ratelaws(465) -ratelaws(473) -ratelaws(499) +ratelaws(516) -ratelaws(525) +ratelaws(563) +ratelaws(569);
    Dspecies(141) = ratelaws(269) -ratelaws(370) +ratelaws(408) -ratelaws(418) +ratelaws(446) -ratelaws(452) -ratelaws(477) -ratelaws(511) +ratelaws(585);
    Dspecies(142) = ratelaws(270) -ratelaws(360) +ratelaws(410) -ratelaws(419) -ratelaws(458) -ratelaws(466) +ratelaws(474) -ratelaws(481) -ratelaws(512) +ratelaws(576) +ratelaws(580);
    Dspecies(143) = ratelaws(272) -ratelaws(371) +ratelaws(422) -ratelaws(432) +ratelaws(448) -ratelaws(453) -ratelaws(478) -ratelaws(513) +ratelaws(586);
    Dspecies(144) = ratelaws(273) -ratelaws(361) +ratelaws(424) -ratelaws(433) -ratelaws(459) -ratelaws(467) +ratelaws(475) -ratelaws(482) -ratelaws(514) +ratelaws(577) +ratelaws(582);
    Dspecies(145) = ratelaws(278) -ratelaws(372) +ratelaws(409) -ratelaws(420) +ratelaws(447) -ratelaws(454) -ratelaws(490) -ratelaws(526) +ratelaws(590);
    Dspecies(146) = ratelaws(279) -ratelaws(362) +ratelaws(412) -ratelaws(421) -ratelaws(460) -ratelaws(468) +ratelaws(487) -ratelaws(494) -ratelaws(527) +ratelaws(578) +ratelaws(581);
    Dspecies(147) = ratelaws(281) -ratelaws(373) +ratelaws(423) -ratelaws(434) +ratelaws(449) -ratelaws(455) -ratelaws(491) -ratelaws(528) +ratelaws(591);
    Dspecies(148) = ratelaws(282) -ratelaws(363) +ratelaws(426) -ratelaws(435) -ratelaws(461) -ratelaws(469) +ratelaws(488) -ratelaws(495) -ratelaws(529) +ratelaws(579) +ratelaws(583);
    Dspecies(149) = ratelaws(284) +ratelaws(292) -ratelaws(378) -ratelaws(530) -ratelaws(536);
    Dspecies(150) = ratelaws(285) +ratelaws(293) -ratelaws(379) -ratelaws(531) -ratelaws(537);
    Dspecies(151) = ratelaws(286) +ratelaws(294) -ratelaws(380) -ratelaws(532) -ratelaws(538);
    Dspecies(152) = ratelaws(287) +ratelaws(295) -ratelaws(381) -ratelaws(533) -ratelaws(539);
    Dspecies(153) = ratelaws(288) -ratelaws(382) -ratelaws(534);
    Dspecies(154) = ratelaws(289) -ratelaws(383) -ratelaws(535);
    Dspecies(155) = ratelaws(386) +ratelaws(396) +ratelaws(404) -ratelaws(540) -ratelaws(554) -ratelaws(556) -ratelaws(558);
    Dspecies(156) = ratelaws(390) +ratelaws(397) +ratelaws(406) -ratelaws(542) -ratelaws(555) -ratelaws(557) -ratelaws(559);
    Dspecies(157) = ratelaws(411) +ratelaws(466) +ratelaws(483) +ratelaws(503) -ratelaws(544) -ratelaws(562) -ratelaws(580) -ratelaws(587) -ratelaws(594);
    Dspecies(158) = ratelaws(413) +ratelaws(468) +ratelaws(496) +ratelaws(518) -ratelaws(545) -ratelaws(563) -ratelaws(581) -ratelaws(592) -ratelaws(599);
    Dspecies(159) = ratelaws(425) +ratelaws(467) +ratelaws(484) +ratelaws(505) -ratelaws(546) -ratelaws(568) -ratelaws(582) -ratelaws(588) -ratelaws(595);
    Dspecies(160) = ratelaws(427) +ratelaws(469) +ratelaws(497) +ratelaws(520) -ratelaws(547) -ratelaws(569) -ratelaws(583) -ratelaws(593) -ratelaws(600);
    Dspecies(161) = ratelaws(444) -ratelaws(541) -ratelaws(572);
    Dspecies(162) = ratelaws(445) -ratelaws(543) -ratelaws(573);
    Dspecies(163) = ratelaws(456) +ratelaws(476) +ratelaws(500) -ratelaws(548) -ratelaws(560) +ratelaws(564) -ratelaws(566) +ratelaws(570) -ratelaws(574) -ratelaws(584) -ratelaws(596);
    Dspecies(164) = ratelaws(457) +ratelaws(489) +ratelaws(515) -ratelaws(549) -ratelaws(561) +ratelaws(565) -ratelaws(567) +ratelaws(571) -ratelaws(575) -ratelaws(589) -ratelaws(601);
    Dspecies(165) = ratelaws(458) +ratelaws(477) +ratelaws(502) -ratelaws(550) +ratelaws(560) -ratelaws(564) -ratelaws(576) -ratelaws(585) -ratelaws(597);
    Dspecies(166) = ratelaws(459) +ratelaws(478) +ratelaws(504) -ratelaws(551) +ratelaws(566) -ratelaws(570) -ratelaws(577) -ratelaws(586) -ratelaws(598);
    Dspecies(167) = ratelaws(460) +ratelaws(490) +ratelaws(517) -ratelaws(552) +ratelaws(561) -ratelaws(565) -ratelaws(578) -ratelaws(590) -ratelaws(602);
    Dspecies(168) = ratelaws(461) +ratelaws(491) +ratelaws(519) -ratelaws(553) +ratelaws(567) -ratelaws(571) -ratelaws(579) -ratelaws(591) -ratelaws(603);

end


end
```
